# Implementing the precautionary approach into fisheries management: Making the case for probability-based harvest control rules

**DOI:** 10.1101/2020.11.06.369785

**Authors:** Tobias K. Mildenberger, Casper W. Berg, Alexandros Kokkalis, Adrian R. Hordyk, Chantel Wetzel, Nis S. Jacobsen, André E. Punt, J. Rasmus Nielsen

## Abstract

The precautionary approach to fisheries management advocates for risk-averse management strategies that include biological reference points as well as decision rules and account for scientific uncertainty. In this regard, two approaches have been recommended: (i) harvest control rules (HCRs) with threshold reference points to safeguard against low stock biomass, and (ii) the *P*\* method, a ‘probability-based HCR’ that reduces the catch limit as a function of scientific uncertainty (i.e. process, model, and observation uncertainty). This study compares the effectiveness of these precautionary approaches in recovering over-exploited fish stocks with various life-history traits and under a wide range of levels of scientific uncertainty. We use management strategy evaluation based on a stochastic, age-based operating model with quarterly time steps and a stochastic surplus production model. The results show that the most effective HCR includes both a biomass threshold as well as the *P*\* method, and leads to high and stable long-term yield with a decreased risk of low stock biomass. For highly dynamics stocks, management strategies that aim for higher biomass targets than the traditionally used *B*_MSY_ result in higher long-term yield. This study makes the case for probability-based HCRs by demonstrating their benefit over deterministic HCRs and provides a list of recommendations regarding their definition and implementation.

## Introduction

Fisheries management success is challenged by high levels of uncertainty inherent to ecosystems and the management process (Garcia 2000). Uncertainty is defined as the “incompleteness of knowledge about the state or process (past, present, and future) of nature” (FAO 1995) and can arise from natural variability in the system, observation error in the data collection process, and the practical implementation of management policies (Francis and Shotton 1997; Rosenberg and Brault 1993). Uncertainty translates directly into risk in the fisheries management process (Fogarty et al. 1996), such as the probability of low stock biomass. For this reason, most fisheries management systems refer to the precautionary principle in their guidelines (e.g. Australian Government 2007; DFO 2009; EU 2013; U.S. Office of the Federal Register 2009). The precautionary approach to fisheries management recognises the potential negative consequences associated with high uncertainty and advocates for the use of predefined decision rules and conservative management actions (FAO 1995). In light of the precautionary principle, one of the key objectives of sustainable fisheries management is maximising expected returns (e.g. measured as the expected catch or revenue) from fisheries while minimising risks, such as the probability of low stock biomass (Dowling et al. 2013; Punt et al. 2001). This risk-yield trade-off predicts that expected returns associated with management tactics increase with the managers’ willingness to take risks (Little et al. 2016). However, larger yields are also linked to higher variability in yield from one year to the next (May et al. 1978). Borrowing the concept of effective portfolios from finance science and based on the risk-yield-variability trade-off, we define a management strategy as ‘effective’ if there are no alternative strategies with: (i) the same or higher expected return and a lower risk, and (ii) the same risk and a higher ore equal expected return. This study compares the effectiveness of various decision rules and conservative management actions in light of the precautionary approach to fisheries management.

Biological reference points such as the fishing mortality are (*F*_MSY_) and biomass (*B*_MSY_) corresponding to the maximum sustainable yield (MSY) remain key components of harvest control rules (HCRs) and concepts in precautionary fisheries management (Garcia 1996; Punt 2010). Apart from empirical HCRs that might be independent of any stock status indicator, HCRs usually link an indicator of the stock abundance, for example, the estimated stock status relative to biological reference points (e.g. *B/B*_MSY_), to specific management actions such as a total allowable catch (TAC) (Punt 2010). The stock status and reference points cannot be observed but rely on estimation using stock assessment methods and are therefore subject to estimation uncertainty. Estimation uncertainty does not only include the uncertainty associated with natural states and processes (process uncertainty) and the measurement thereof (observation uncertainty), but also uncertainty due to structure of the estimation model (model uncertainty) (Francis and Shotton 1997). These sources of uncertainty are also referred to as scientific uncertainty (e.g. Punt and Donovan 2007b). Stochastic estimation models allow the scientific uncertainty associated with current and future stock status to be quantified as probability or likelihood distributions. Two approaches for including the precautionary approach into fisheries management have been recommended and are used in this study: (i) HCRs with threshold reference points to safeguard against low stock biomass in the face of high uncertainty (e.g. Da-Rocha et al. 2016), and (ii) ‘probability-based HCRs’ that reduce the catch limit, such as the overfishing limit (OFL) or TAC, as a function of quantified or derived uncertainty of current or future stock status explicitly (e.g. Dankel et al. 2016; Shertzer et al. 2008; Wiedenmann et al. 2017).

Threshold reference points, in particular those related to stock biomass, play an important role in optimal harvesting theory (e.g. Lande et al. 2001). When predicted biomass (current or future) falls below the threshold, fishing effort is reduced or terminated. Biomass thresholds are the foundation for escapement strategies, where the survival (‘escapement’) of a certain stock biomass size is desired (Beddington and May 1977; Getz et al. 1987; Lett and Doubleday 1976). Both, the absolute threshold values and the definition of the threshold vary widely. For example, the threshold can be defined based on an inflection point in the stock-recruitment relationship (e.g. ICES 2018), on estimated historical biomass (e.g. ICES 2018), or as a fraction of *B*_MSY_ (e.g. 50%*B*_MSY_) (ICES 2017). This study evaluates the effect of alternative threshold levels defined as a fraction of *B*_MSY_ on the performance of the HCR.

In contrast to deterministic HCRs, probability-based rules quantify scientific uncertainty and reduce the catch limit using an uncertainty buffer (Prager and Shertzer 2010). The buffer can, for example, be derived by defining the acceptable risk (or probability) that predicted fishing mortality is above or below biological reference points (Caddy and McGarvey 1996). This method is formally known as the *P*\* method, and was later refined to include the uncertainty of the reference points (Prager et al. 2003) and to be used on the predicted catch distribution rather than the fishing mortality rate (Prager and Shertzer 2010). The distribution for the different quantities can be estimated within the assessment method, by means of simulation (Privitera-Johnson and Punt 2020), or be a predefined measure of the uncertainty based on stock characteristics such as the amount and quality of available data. For instance, Ralston et al. (2011) provides measures of the scientific uncertainty (standard deviations in log-space) associated with current spawning stock biomass derived from a meta-analysis. In contrast to previous studies (e.g. Dankel et al. 2016; Wiedenmann et al. 2017), we apply the *P*\* method to all components of the HCR and use assessment-based uncertainty.

Based on the risk-yield-variability trade-off, we evaluate the effectiveness of various probability-based and deterministic HCRs by focusing on (i) the comparison between biomass thresholds and the *P*\* method, (ii) the effect of scientific uncertainty on deterministic and probability-based HCRs, and (iii) the definition of probability-based HCRs using the *P*\* method for different components of the HCR (e.g. predicted catch or *F/F*_MSY_). We use management strategy evaluation (MSE) to compare the performances of the HCRs. MSEs simulate populations as well as the feedback between the population and the successive applications of management strategies in a closed loop (Punt et al. 2016; Smith 1994). We use a stochastic age-based operating model with quarter-year time steps to determine the population dynamics of three stocks with different life history traits. Then, we employ a stochastic production model to estimate both stock status and biological reference points with associated uncertainty. The HCR recommends a TAC based on estimated stock status that is used in the operating model to project the stock forward, i.e. closed-loop simulation framework. We show that the most effective HCR combines the biomass threshold with the *P*\* method and leads to high and stable long-term yield while reducing the risk of overfishing.

## Methods

### Operating model

The operating models were based on the life history characteristics of three marine fish species from different geographical regions in the North Atlantic: (i) anchovy (*Engraulis encrasicolus*; ICES stock code: ane.27.8) in the Bay of Biscay, representing a fast-growing species, (ii) haddock (*Melanogrammus aeglefinus*; ICES stock code: had.27.7.b-k) in the Celtic Seas, representing a species with intermediate parameters, and (iii) Greenland halibut (*Reinhardtius hippoglossoides*; ICES stock code: ghl.27.1-2) in the Northeast Arctic, representing a slow-growing species (Table 1). We simulated the population dynamics of the three species using an age-structured population model with quarterly time steps described in detail in the Supplementary Section A. We will refer to the three stocks as anchovy, haddock, and halibut.

**Table 1:** Parameter values for the anchovy, haddock, and halibut stocks representing species with fast-growing, intermediate and slow-growing life history parameters. Growth parameters (*L*_∞_, *K*) correspond to the von Bertalanffy growth equation (Bertalanffy 1938). Steepness (h) corresponds to the Beverton-Holt stock-recruitment relationship (Mace and Doonan 1988). Selectivity and maturity parameters correspond to a logistic function (Supplementary Equations A4 and A5). The references for the biological parameters of each stock are given in Supplementary Table A1.

| Parameter | Description | Stocks |  |  |
| --- | --- | --- | --- | --- |
|  |  | Anchovy<sup>1</sup> | Haddock | Halibut |
| $L_\infty$ | Asymptotic length [cm] | 18.69 | 76.9 | 120 |
| $K$ | Growth rate [1/yr] | 0.89 | 0.197 | 0.073 |
| $a_0$ | Age at length=0 [yr] | -0.02 | -0.092 | -0.1 |
| $a$ | Length-weight scaling factor [g/cm<sup>b</sup>] | $4.8e^{-3}$ | $6.45e^{-3}$ | $3.33e^{-3}$ |
| $b$ | Length-weight exponent | 3.134 | 3.108 | 3.249 |
| $a_{max}$ | Maximum age [yr] | 6 | 17 | 27 |
| $h$ | Steepness | 0.75 | 0.74 | 0.79 |
| $L_{m50}$ | Length at 50% maturity [cm] | | 22 | 71.2 |
| $L_{m95}$ | Length at 95% maturity [cm] | | 29 | 81.2 |
| $L_{s50}$ | Length at 50% selectivity [cm] | | 13.48 | 51 |
| $L_{s95}$ | Length at 95% selectivity [cm] | | 25.17 | 58.23 |
| $\sigma_R$ | Standard deviation of the recruitment deviations | 0.766 | 0.748 | 0.636 |
| $\rho_R$ | Coefficient of auto-correlated recruitment deviations | 0.435 | 0.404 | 0.437 |

<sup>1</sup> The maturity and selectivity parameters are missing for anchovy as these processes were defined by age for this stock (Supplementary Figure A1).

We initialised the MSE with 40 years of data referred to as the historical period. The historical period reflects the amount of relevant and standardised data available for many stocks (e.g. ICES 2019b). The stocks We assumed that fishing effort was increasing over time and identified the fishing mortality rate that lead to stock biomass of approximately 0.5*B*_MSY_ at the end of the historical period (Supplementary Figure A2). The over-exploited state enables the evaluation of the HCRs ability to recover stocks, and also amplifies the differences between HCRs. We added bias-corrected noise with lognormally distributed deviations to the historical fishing mortality rate: 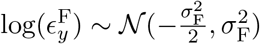 with *σ*_F_ = 0.15 (Carruthers et al. 2014). Finally, we run the projection period of 40 years, which is equal to the historical period and exceeds the maximum age of the longest lived of the three species (halibut: 27 yr).

### Assessment model

The HCRs evaluated in this study require the quantification of stock status (relative to fishery reference points) and thus the application of a stock assessment method. In line with previous studies investigating probability-based HCRs (Caddy and McGarvey 1996; Prager et al. 2003; Prager and Shertzer 2010), we applied a production model to estimate reference points and stock status. In particular and following the guidelines of ICES (ICES 2017), we used the stochastic production model in continuous time (SPiCT; Pedersen and Berg 2017) as the stock assessment method. SPiCT is a state-space re-parameterised version of the Pella-Tomlinson surplus production model (Fletcher 1978; Pella and Tomlinson 1969), i.e. quantifies uncertainty in the observation and process equations. Thus, SPiCT has the potential to derive the probability distributions of the three quantities important to fisheries management and that are part of the HCR: fishing mortality rate relative to *F_MSY_* at the start of the management year (*F_y_/F_MSY_*), the predicted biomass relative to *B*_*MSY*_ at the end of the management year *B*_*y*+1_/*B_MSY_*, and the predicted catch during the management year *C*_*y*+1_. The predicted catch in year *y* is estimated by:

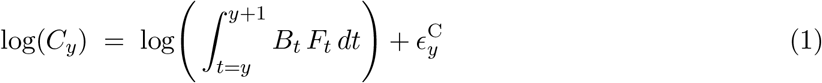

where *C*_*y*_ is the predicted annual catch, *B_t_* and *F_t_* are the exploitable biomass and fishing mortality rate at time t, respectively (Supplementary Table A2), and the observation error is 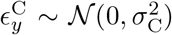. SPiCT approximates continuous time by means of the Forward Euler scheme (Iacus 2009), i.e. using small time-steps within a single year (Pedersen and Berg 2017). All model parameters (9 fixed parameters, Supplementary Table A3) are estimated by maximum likelihood using automatic differentiation, as implemented in Template Model Builder (**\{**)Kristensen2016}. The uncertainties of all quantities of SPiCT are estimated using the delta method assuming asymptotically normal distributions in log space (Kristensen et al. 2016; Pedersen and Berg 2017). We used the recommended default model configuration (Pedersen and Berg 2017), i.e. three vague prior distributions for parameter n defining the shape of the production curve 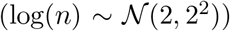 and for the hyper parameters 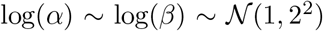, corresponding to the ratios of the standard deviations of observation to process noise terms: 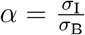 and 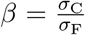 (c.f. Supplementary Table A2 and A3). For computational reasons, we decreased the number of time steps of the Forward Euler scheme from the default 16 per year to 4 per year. We evaluated the sensitivity to the assumed prior distributions and the decreased number of Euler time steps (Supplementary Section C).

### Data simulation

Required input data for a SPiCT assessment consists of a time series of catches and a relative abundance index (Pedersen and Berg 2017). We simulated annual catches and two time-series of abundance indices for the whole (40 yr) historical time period. Annual catch observations were calculated as the sum of quarterly catches in weight (Supplementary Equation A12). The abundance indices correspond to the exploitable stock biomass, i.e. the part of the total stock biomass that is vulnerable to the commercial gear. The timing of the two surveys at the start of the year (1^*st*^ quarter) and mid-year (3^*rd*^ quarter) correspond to the ICES International Bottom Trawl Surveys (IBTS) in the North Sea (ICES 2012). We simulated lognormal observation noise for the annual catches 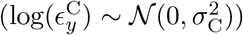 and the abundance indices 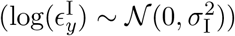 with *σ*_C_ = *σ*_I_ = 0.3 (Supplementary Equations A15 and A16). We evaluated the effect of different levels of observation noise on the performance of the HCRs by varying *σ*_C_ and *σ*_I_ within the range from 0.1 to 0.7, covering values used in other studies (e.g. Carruthers et al. 2014; Wiedenmann et al. 2017). Additionally, we explored the effect of implementation uncertainty by simulating lognormal distributed deviations on the realised fishing mortality rate in a sensitivity analysis (Supplementary Section C). The standard deviation (SD) of 0.15 is within the range of implementation uncertainty assumed by other studies (**\{**)Walsh2018b,Nieland2008,Fischer2020}.

### Harvest control rules

This study assumes that advice is given annually for the next fishing period (year *y* + 1) based on a stock assessment at the start of the same year using fishery-dependent and -independent information from all previous years (until then end of year *y*). In fact, many management systems include an intermediate year or assessment year, i.e. advice is given for year *y* + 1 based on an assessment in year *y* using data up until year *y* − 1 (abundance indices or seasonal catches might be available at the start of the assessment year *y*; c.f. timeline in Supplementary Fig A5) (e.g. ICES 2019b). In this case, assumptions about fishery and biological processes during the assessment year *y* are required to perform a short-term forecast and predict the catch in the management year *y* + 1. We explore the effect of having an intermediate year and two assumptions about the catch herein (continuation of the F-process or catch equals TAC from previous year) in a sensitivity analysis (Supplementary Section C).

We define the recommended TAC for any period (here year *y* + 1) as a fractile of the catch distribution predicted by the assessment model given a target fishing mortality rate for that period 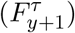:

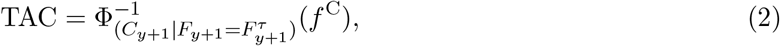

where Φ^−1^(*f*^C^) is the inverse distribution function of the predicted catch given the target fishing mortality rate and *f*^C^ ≤ 0.5 is the risk fractile for the predicted catch distribution. Note, that the risk fractile (*f*^C^) is identical to the *P*\* value of the *P*\* method, whereas *P*\* does not indicate which quantity it is used for, e.g. predicted catch or relative fishing mortality. Risk fractiles less than 0.5 are considered as they are more precautionary than the median and take into account the estimated uncertainty. The target fishing mortality rate 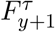 is defined by

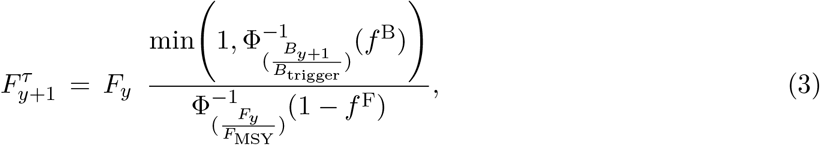

where the biomass threshold (BT) is defined as *B*_trigger_ = *xB*_MSY_, *x* > 0, and *f*^B^ ≤ 0.5 and *f*^F^ ≤ 0.5 are the risk fractiles of the distributions of the relative biomass and fishing mortality rate, respectively. Note that 1 − *f*^*F*^ is used for the *F/F*_MSY_ distribution (Eq. 3), i.e. a smaller risk fractile implies a larger fractile for this distribution. We define the inverse distribution function for all quantities in log space. In other words, this rule implies that the TAC is based on fishing at *F*_MSY_ if the stock biomass is equal or above the biomass threshold (*B*_trigger_). When the biomass is less than the BT, 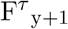 is set to 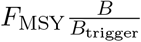, i.e. F is reduced linearly to zero as the estimated biomass decreases and *F* = 0 at *B* = 0.

A wide range of HCRs are nested within this HCR formulation: The ‘MSY rule’ or ‘fishing at *F*_MSY_’ is obtained by setting the numerator to 1, i.e. no BT is used, and by setting *f*^C^ = *f*^F^ = 0.5, i.e. the median of all distributions is used. The ‘hockey-stick MSY rule’ for surplus production models (ICES 2017) emerges if *x* = 0.5 and *f*^C^ = *f*^B^ = *f*^F^ = 0.5. The formulation can also define probability-based HCRs (Prager et al. 2003), by using any risk fractile (*f*^C^, *f*^B^, *f*^F^) smaller than 0.5. Figure 1 conceptualises a deterministic HCR (grey) in comparison to two probability-based HCR with *f* = 0.25 for all three quantities (orange) and the catch distribution only (blue). When approaching the limit where observations are perfect (no observation noise) and the model is perfect (no process noise), the recommended TAC is independent of the risk fractile chosen.

**Figure 1:**
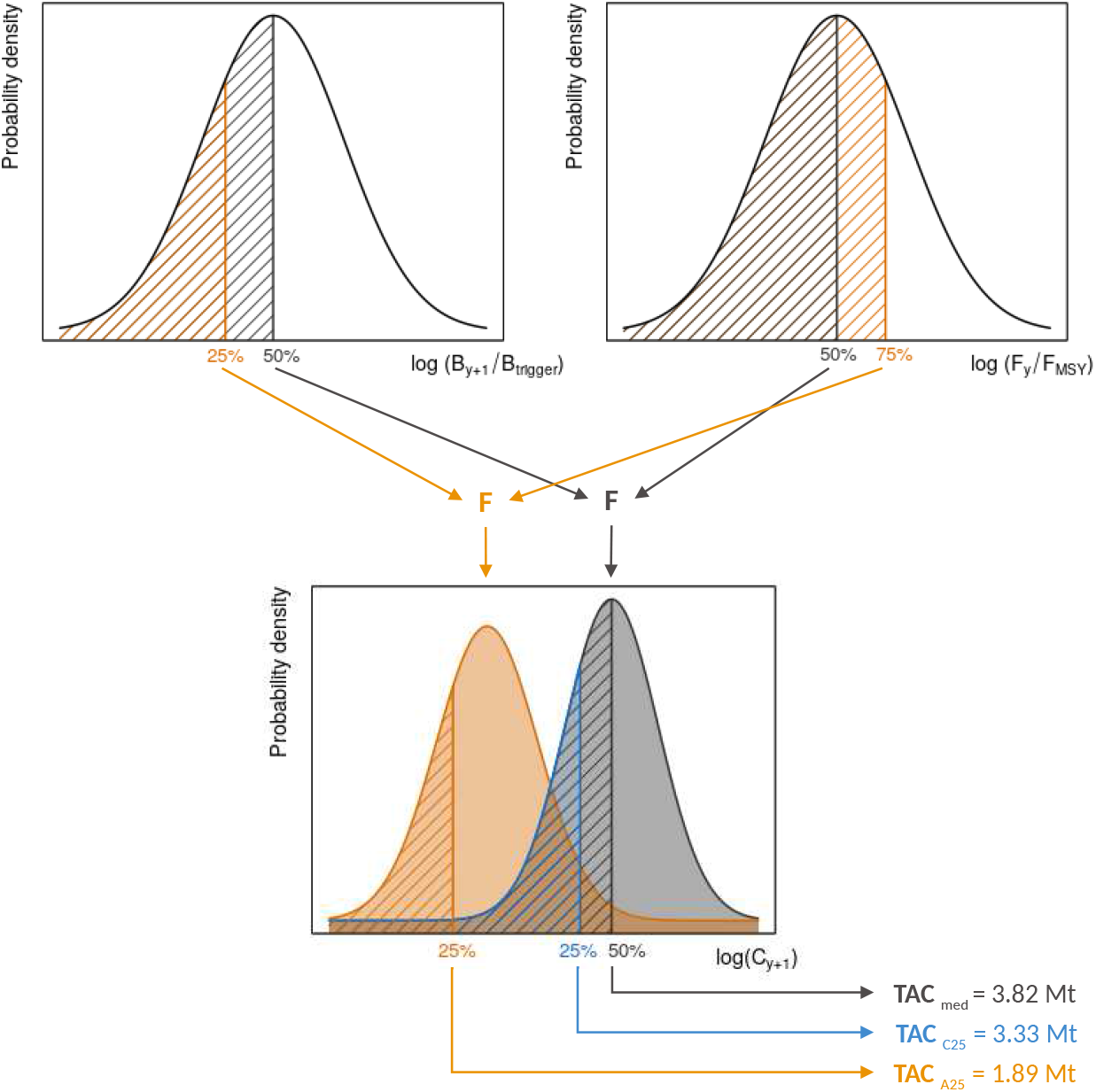
Probability densities in log space of three quantities relevant for fisheries management: *F_y_/F_MSY_*, *B*_*y*+1_*/B*_trigger_, and *C*_*y*+1_; and their use within three harvest control rules (HCR; different colours). Based on any percentile of the relative F and B distributions (upper panels), the target F is estimated, which leads to different predicted catch distributions (lower panel). The recommended TAC can be defined as any percentile of the predicted catch distribution. *TAC_med_* (grey) reflects a deterministic HCR based on the median of all three distributions. *TAC_C_*_25_ (blue) reflects a HCR based on the median of *F_y_/F*_MSY_ and *B_y_*_+1_*/B*_trigger_, and the 25th percentile of the *C*_*y*+1_ distribution. *TAC_A_*_25_ (orange) reflects a HCR based on the 25th percentile of *B*_*y*+1_*/B*_trigger_, predicted catch, and *F_y_/F*_MSY_ distribution.

For the purpose of this study, we defined 60 HCR variations based on equations 2 and 3. These HCRs aim to investigate the following:

- For the comparison of the effect of BTs and the P* method, we defined eight HCRs with different BTs with *x* ∈ [0.25, 2], including the recommendation of *x* = 0.5 for surplus production models (ICES 2017). These HCRs are denoted as BT_x_, e.g. BT_0.25_ or BT_1.5_. Further, we defined the MSY rule (*f*_0.5_) and six HCRs with different risk fractiles for the predicted catch distribution *f*^C^ ∈ [0.005, 0.45], e.g. 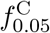 or 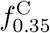.
- For the quantification of the effect of different levels of scientific uncertainty on the performance of deterministic and probability-based HCRs as well as all sensitivity analyses, we used the three HCRs defined above: *f*_0.5_, 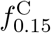, and BT_0.5_, and added a probability-based threshold rule 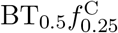, i.e. combining the threshold and *P*\* method.
- For the evaluation of probability-based rules accounting for the uncertainty of the different components of the HCR: *C*_*y*+1_, *F_y_/F*_MSY_, and *B*_*y*+1_*/B*_trigger_, we defined 18 HCRs without BT and 30 HCRs with a BT (*x* = 0.5; including HCRs defined above). To reduce the number of combinations and thus computations, we used the same risk fractile if several quantities are considered, e.g. *f*^C^ = *f*^F^ = 0.25. As above, the labels indicate if a BT was used (BT_0.5_), and which quantities (*f*^C^, *f*^F^, *f*^B^) and fractiles (*f* ∈ [0.005, 0.5]) were used. For example, 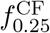 to a HCR with a risk fractile 0.25 for the quantities *C*_*y*+1_ and *F_y_/F*_MSY_, and 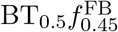 refers refers to a HCR with a BT with *x* = 0.5 and the risk fractile 0.45 for the quantities *F_y_/F*_MSY_ and *B_y_*_+1_*/B*_trigger_. The full list of all HCRs defined in this study is presented in Supplementary Table A4.

In addition to these HCRs, we defined two reference rules: (1) no fishing (*F* = 0) and (2) fishing with the ‘true’ *F*_MSY_ (*F* = *F*_MSY_), i.e. the catch depends on following quantities without observation uncertainty: numbers-at-age, selectivity-at-age, and *F*_MSY_.

### Performance metrics

We evaluated the performance of the HCRs based on following metrics:

- Risk of overfishing (*Prop*(*B* < *B*_lim_)) defined as the average proportion of simulation years in which the true biomass (i.e. B of the operating model) is below the limit reference biomass *B*_lim_ over replicates (ICES 2013), where *B*_lim_ is 20% of the virgin biomass (Dichmont et al. 2017). The 95% confidence intervals of the average risk were estimated by the Wilson score interval method for Binomial proportions (Wilson 1927).
- Yield defined as the median catch relative to the catch obtained with the reference rule *F* = *F*_MSY_ over simulation years and replicates. The 95% simulation intervals of relative yield were estimated by the modified Cox method (Olsson 2005).
- Annual Absolute Variability in yield (AAV) defined as the median annual differences in yield over replicates (Punt 2003) :

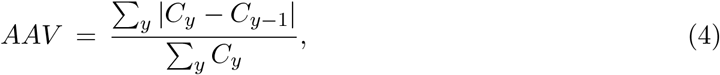

where *C*_*y*_ is the catch during year y.

For the comparison of the performance of HCRs among scenarios with different levels of scientific uncertainty, we estimated the difference in risk and yield associated with low observation uncertainty (SD=0.1) from the other scenarios (SD ∈ [0.1, 0.7]). We estimated the 95% confidence intervals for the risk and yield differences according to Newcombe and Altman (2000) and Price and Bonett (2002), respectively. In addition to estimating the metrics for the whole management period, i.e. years 41-80 for risk and yield and years 42-80 for AAV to avoid the effect of the last historical catch, we define short-term risk and yield corresponding to the first 5 years of the projection period (years 41-45) and long-term risk and yield corresponding to the last 15 years of the projection period (years 66-80).

We conducted 500 replicates for each stock and excluded any replicate with one or more nonconverged SPiCT assessment(s) in any year from the calculation of the performance metrics, i.e. the performance metrics are based only on replicates in which all SPiCT assessments converged. Furthermore, for the comparison of scenarios with different levels of scientific uncertainty, we only used replicates that converged in all years across HCRs and scenarios. Thus, to account for the lower sample size, in these specific scenarios, we conducted 2000 replicates for each stock. We evaluated the stability of all performance metrics against the number of replicates and results indicated that the number of replicates for each stock was sufficient.

## Results

The number of converged replicates varied between 51% and 86% corresponding to 255 to 430 replicates depending on the HCR and species. On average, 77%, 80%, and 81% of the replicates converged for the baseline scenario for anchovy, haddock, and halibut, respectively (Supplementary Table B1). First, we focus first on the probability-based rules with the risk fractile on the predicted catch distribution (e.g. 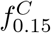) and HCR with a biomass threshold (e.g. *BT*_0.75_).

### Biomass thresholds vs P* method

The two quantitative precautionary approaches evaluated in this study, biomass thresholds and the *P*\* method, reduce the risk of overfishing defined as *Prop*(*B* < *B*_lim_) in comparison to HCRs without any of the approaches. The absolute risk reduction varies among the stocks with different life-history traits and depends on the time since start of the management as well as the threshold and the risk fractile, with higher thresholds and lower fractiles leading to larger risk aversion. For haddock and halibut, the risk-yield trade-off defined by the thresholds and risk fractiles describes a proportional relationship on the short-term, implying that any risk reduction comes at the immediate cost of loss in yield (Fig. 2). Over the long-term, however, the use of certain (larger) thresholds and (smaller) fractiles reduces risk without any appreciable loss in yield (Fig. 2). While for anchovy, the short-term risk-yield trade-off resembles the long-term trade-off of the other stocks, the long-term trade-off implies an increase of yield in combination with a decrease in risk for certain thresholds and risk fractiles.

**Figure 2:**
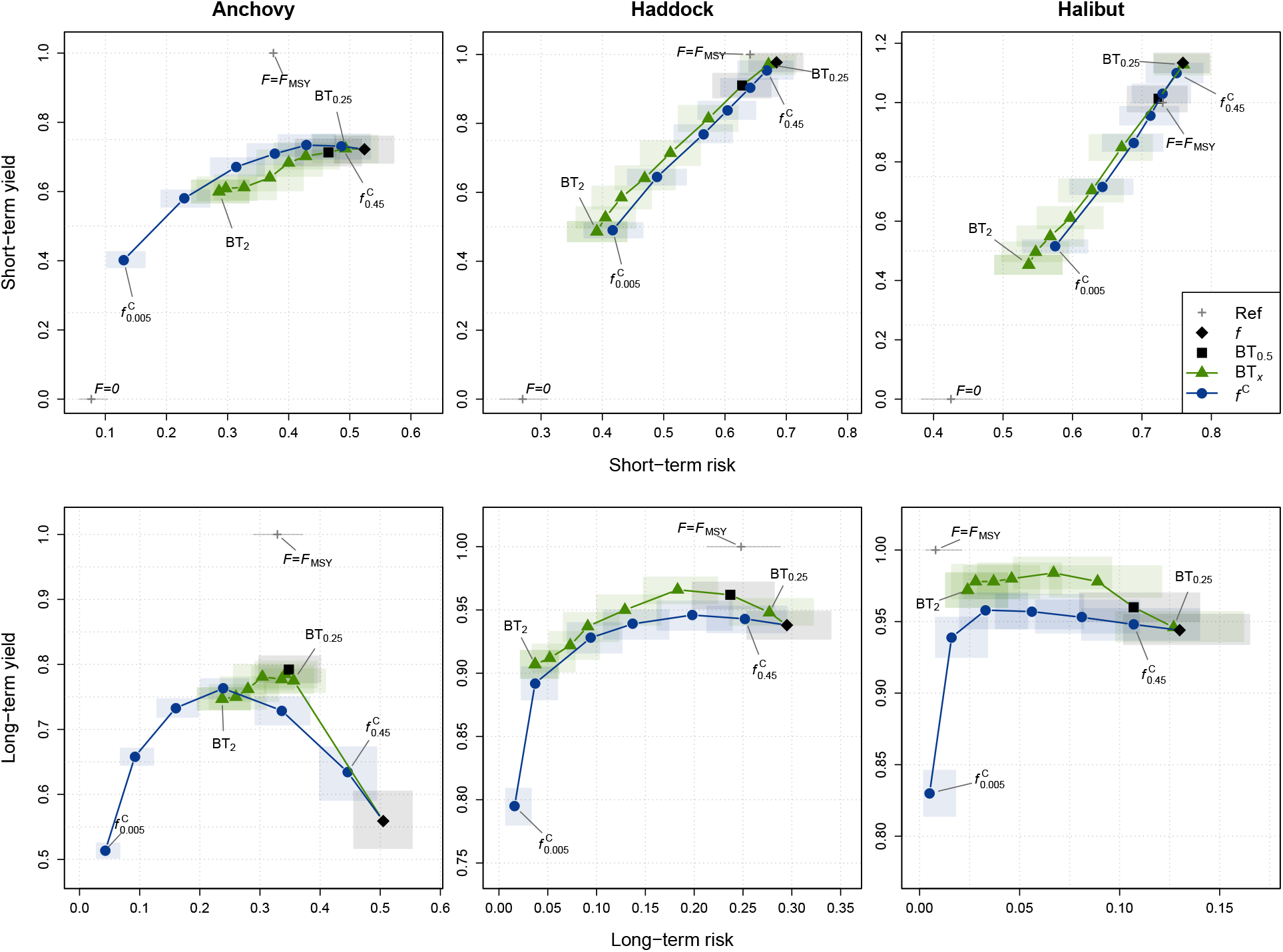
Risk-yield trade-off over the short term (years 41-45; upper row) and long term (years 66-80; lower row) for anchovy, haddock, and halibut (columns). Starting at the black diamond (MSY rule) the HCRs with an uncertainty buffer (blue circles) are: 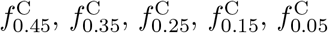 and 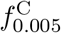, while the HCRs with a biomass threshold (green triangles) are: BT_0.25_, BT_0.5_, BT_0.75_, BT_1_, BT_1.25_, BT_1.5_, BT_1.75_, and BT_2_. These sequences are connected by lines and the first and last HCRs in that sequence are labelled. The two grey crosses represent the two reference rules: fishing at true F_MSY_ (F = F_MSY_) and no fishing (F = 0). Note that the ‘no fishing rule’ is not included in the long-term plots to reduce the range of the y axis. The shaded areas represent the 95% confidence intervals of the yield and risk.

Due to the over-exploited conditions at the start of the projection period in the last historical year (year 40), for haddock and halibut, all HCRs show high absolute risk levels above 0.4 in the short-term. Even the reference rule without fishing (*F* = 0) shows a high absolute risk with levels of 0.28 and 0.44 for haddock and halibut, respectively (Fig. 2). The long-term risk levels of all HCRs are significantly lower and below 0.35 for haddock and below 0.15 for halibut. In contrast, the short- and long-term risk levels of all HCRs are similar for anchovy and less than 0.4. The anchovy stock shows a similar short- and long-term yield between 0.4 and 0.8 for all HCRs, while the other stocks show a high short-term yield close or even higher (halibut) than the yield of the *F* = *F*_MSY_ reference rule. In particular, on the short-term fishing at *F*_MSY_ might not lead to the highest yield. On the long-term, the yield is between 0.8 and 1 for all HCRs for haddock and halibut.

Overall, biomass thresholds have a slightly more efficient risk-yield trade-off than the risk fractiles, i.e. they reach more desirable locations in the risk-yield space (higher yield for same risk or same yield with lower risk), in particular considering the long-term trade-off. This can be attributed to the fact that the rules leads to *B*_MSY_ the fastest rather than fishing at *F*_MSY_ leads to the theoretical highest yield, i.e. F potentially larger than *F*_MSY_ whenever *B* > *B*_MSY_, *F* = 0 when *B* < *B*_MSY_, and only *F* = *F*_MSY_ when *B* = *B*_MSY_ (e.g. Hening et al. 2019). Thus, it is to be expected that threshold rules (with thresholds ≤ *B*_MSY_) can give a higher yield than the MSY rule and the *P*\* method. This is confirmed by the time series plots, which show that the threshold rules lead to a large reduction in recommended TAC (and thus F) at the start of the projection period when the stock is highly depleted leading to fast stock recovery, but recommend higher TACs than the risk fractiles when the stock is recovered (Supplementary Figure B1).

For the *P*\* method, the yield-variability trade-off evaluated over the whole projection period is similar to the risk-yield trade-off, i.e. the larger the risk fractiles the lower the variability in yield (Fig. 3). By contrast, a larger threshold leads to a higher variability in yield up to a certain threshold of 1.5, 1.25, and 1.0 times *B*_MSY_ for the three stocks respectively. With even higher thresholds, the variability in yield decreases again. The same pattern is true for the TAC recommended by the assessment model (Supplementary Figure B2). The absolute values of AAV vary among species with values up to 0.6 for anchovy to up to 0.16 for halibut.

**Figure 3:**
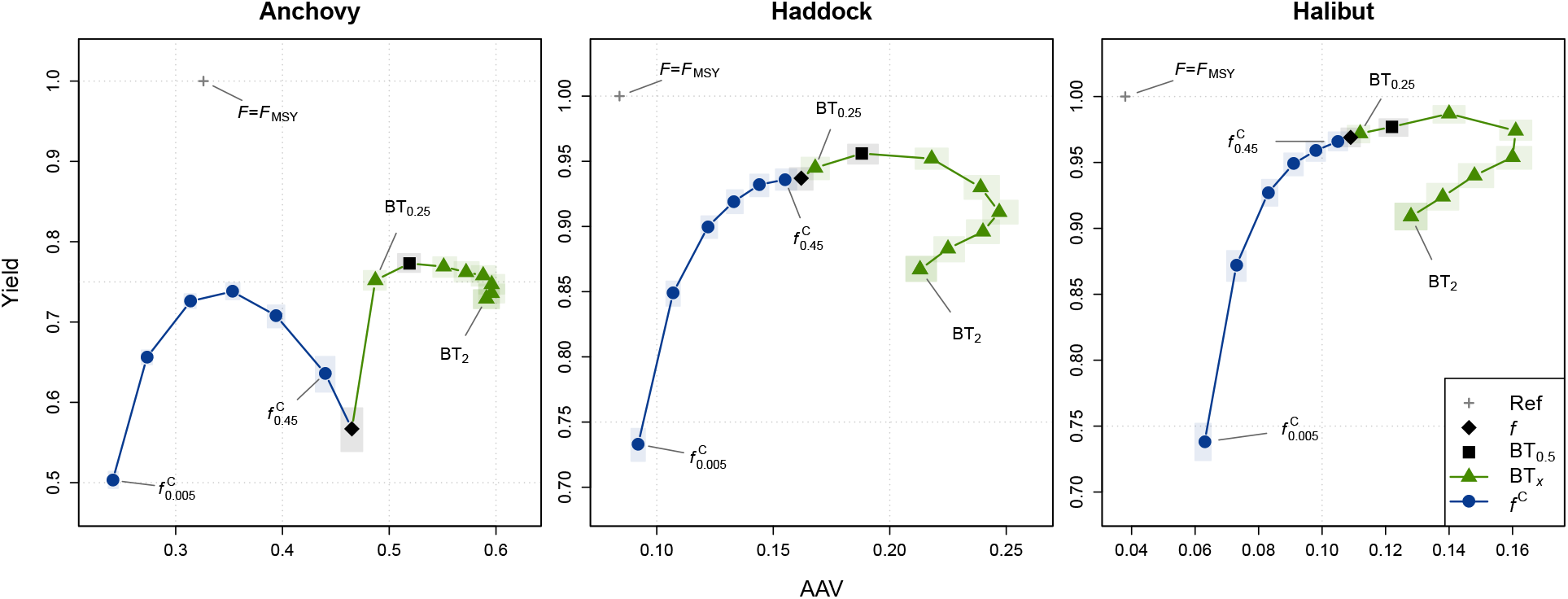
Yield-variability trade-off for the full projection period (years 41-80 for yield and years 42-80 for AAV) for anchovy, haddock, and halibut. Starting at the black diamond (MSY rule) the HCRs with an uncertainty buffer (blue circles) are: 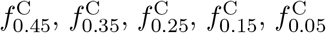 and 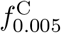, while the HCRs with a biomass threshold (green triangles) are: BT_0.25_, BT_0.5_, BT_0.75_, BT_1_, BT_1.25_, BT_1.5_, BT_1.75_, and BT_2_. These sequences are connected by lines and the first and last HCRs in that sequence are labelled. The two grey crosses represent the two reference rules: fishing at true F_MSY_ (F = F_MSY_) and no fishing (F = 0). The shaded areas represent the 95% confidence intervals of the yield and AAV.

### Probability-based and deterministic HCRs vs. scientific uncertainty

Although biomass thresholds and the *P*\* method reduce risk as a function of the threshold and risk fractile, there is a clear advantage of probability-based rules (here: *P*\* method) over deterministic rules (here: all rules without the *P*\* adjustment, e.g. the MSY rule). While increasing scientific uncertainty leads to higher risk levels for deterministic rules, probability-based rules quantify and account for the increased uncertainty leading to similar risk levels for a range of scientific uncertainty (Fig. 4). The expected decrease in yield associated with deterministic and probability-based rules due to larger uncertainty depends on the species. While for anchovy the decrease in yield is larger for deterministic rules, deterministic rules provide a relative constant yield for halibut (Fig. 4). Biomass threshold rules show a similar sensitivity to scientific uncertainty as the MSY rule, although with a lower relative loss in yield (green lines in Fig. 4). The sensitivity to risk of fractile rules is to a large extent defined by the fractile. While the 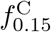 rule shows almost no increase in risk, the 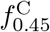 shows a similar sensitivity than the MSY rule. On the other side, the higher the fractile the larger the loss in yield. When considering both risk and yield, the rule with the biomass threshold and the *P*\* method combined seems to perform best (low increase in risk and low decrease in yield; yellow line in Fig. 4).

**Figure 4:**
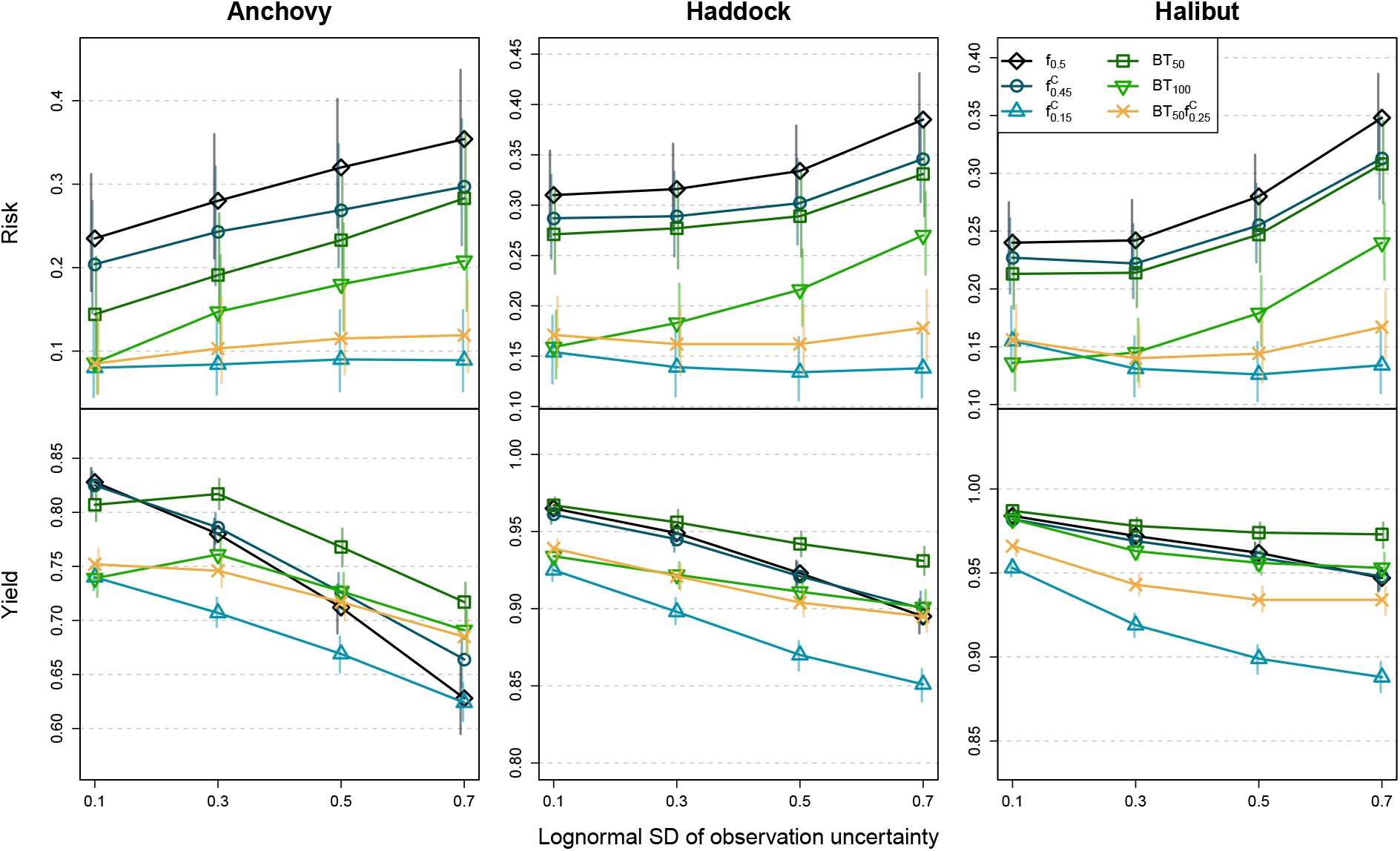
Risk and yield associated with various HCRs for each observation uncertainty scenario (SD [0.1, 0.7]). Vertical lines represent the 95% confidence intervals. Risk and yield are evaluated over the full projection period (years 41-80).

For all harvest rules, increased scientific uncertainty leads to larger variability in stock biomass. However, the distributional properties of the stock biomass differ markedly between deterministic and probability-based rules and explain their differing sensitivity to scientific uncertainty: while the median biomass trajectory of the deterministic rules (e.g. *f*_0.5_ or *BT*_100_) overlap for different scientific uncertainty levels, the distribution is wider for larger scientific uncertainty (comparing SD = 0.1 to SD = 0.7; Fig. 5). Similarly, probability-based rules (e.g. 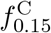) show a wider biomass distribution for larger scientific uncertainty, however the median biomass is larger. Consequently, the lower bounds of the biomass distribution (e.g. 10th percentile in Fig. 5) are similar or even larger for increased scientific uncertainty, leading to similar risk levels for various uncertainty levels. The differing distributional properties of the stock biomass can be attributed to the TAC recommended by the rules. Increased scientific uncertainty leads to a higher variability in TAC and a lower median TAC recommended by the deterministic rules (e.g. *f*_0.5_ and *BT*_0.5_). This implies a wider and more positively skewed fishing mortality distribution, ultimately leading to a wider biomass distribution (Supplementary Figures B4-B7). In contrast, probability-based rules (e.g. 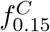) also recommend a lower median TAC with increased scientific uncertainty. However the TAC is less variable and the fishing mortality rate does not exhibit the large positive outliers, leading to a larger median stock biomass (Supplementary Figure B6).

**Figure 5:**
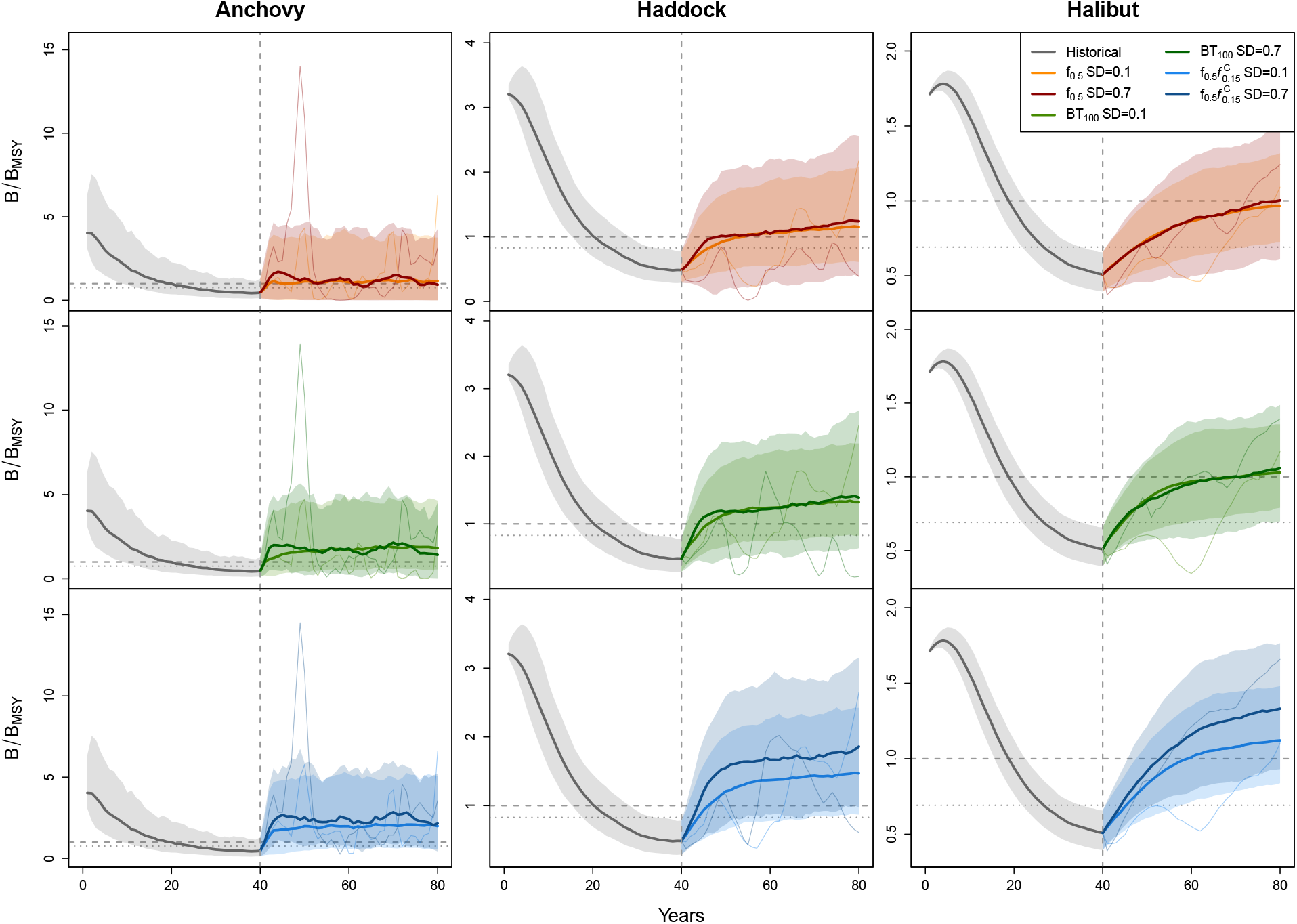
Biomass relative to *B*_MSY_ for the MSY rule (*f*_0.5_, top row), a threshold rule (BT_100_, middle row), and a fractile rule (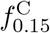, bottom row) and for two levels of scientific uncertainty (SD = 0.1, lighter color, SD = 0.7, darker color). Dotted line corresponds to *B*_lim_*/B*_MSY_. Shaded area extends from the 10th and 90th percentile. Finer lines correspond the trajectory of a single simulation for each rule.

The results for the comparison across observation uncertainty levels are based on replicates which converged across all HCRs and uncertainty levels. Thus, a large proportion of 93%, 78%, and 68% of the 2000 replicates was rejected for anchovy, haddock, and halibut, respectively, corresponding to 137, 448, and 639 replicates for the three stocks, respectively.

### Definition of probability-based HCRs

So far, we only considered the probability-based rules with the risk fractile on the predicted catch distribution (e.g. 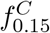), however, the *P*\* method can be applied to any quantity or multiple quantities of the HCR at the same time. The results show that independent of the quantities considered, the risk fractiles lead to a similar risk-yield trade-off as described above for the *f*^C^ rules (Fig. 6). In fact, the predicted yield achieved at chosen risk levels is very similar across the different rule types (varying between 72% to 79%, 87% to 92%, and 89% to 97% for anchovy, haddock, and halibut, respectively; Supplementary Table B6). Nevertheless, the absolute risk level and yield depends strongly on the quantities considered (Fig. 6; c.f. threshold and risk fractile required to reach chosen risk level in Supplementary Table B6). In that regard, three general patterns become apparent: (i) The more quantities considered the larger the effect of the risk fractile, i.e. *f*^CF^ and 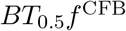 show the largest risk reduction for a given risk fractile. (ii) The risk fractile on *F/F*_MSY_ has a larger effect on the risk reduction than the risk fractile on the predicted catch distribution (e.g. 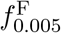 vs. 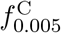). (iii) Using the risk fractile on the *B/B*_trigger_ distribution alone (*f*^B^) leads to the smallest risk reduction. In addition, *f*^B^ rules show a less consistent pattern for the three stocks: While they lead to the highest yield for haddock and halibut, they lead to much lower yield than the other threshold fractile rules for anchovy (Fig. 6). This can be attributed to the fact that for anchovy, the distribution of the recommended TAC is wider and positively skewed, leading to a lower median yield in comparison to another rule 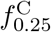 with a similar risk level (Supplementary Figure B9). For all species, *f*^B^ rules lead to increased variability in yield with decreasing risk fractiles (lower row in Fig. 6). This implies that the effect of a smaller risk fractile on the biomass relative to the threshold is comparable to a larger threshold (Fig. 3), leading to higher variability in TAC recommendations. For all other rules, smaller risk fractiles lead to lower variability in yield. However, the results indicate that very small risk fractiles (below 0.05) can lead to a sharp and significant increase in AAV (Fig. 6).

**Figure 6:**
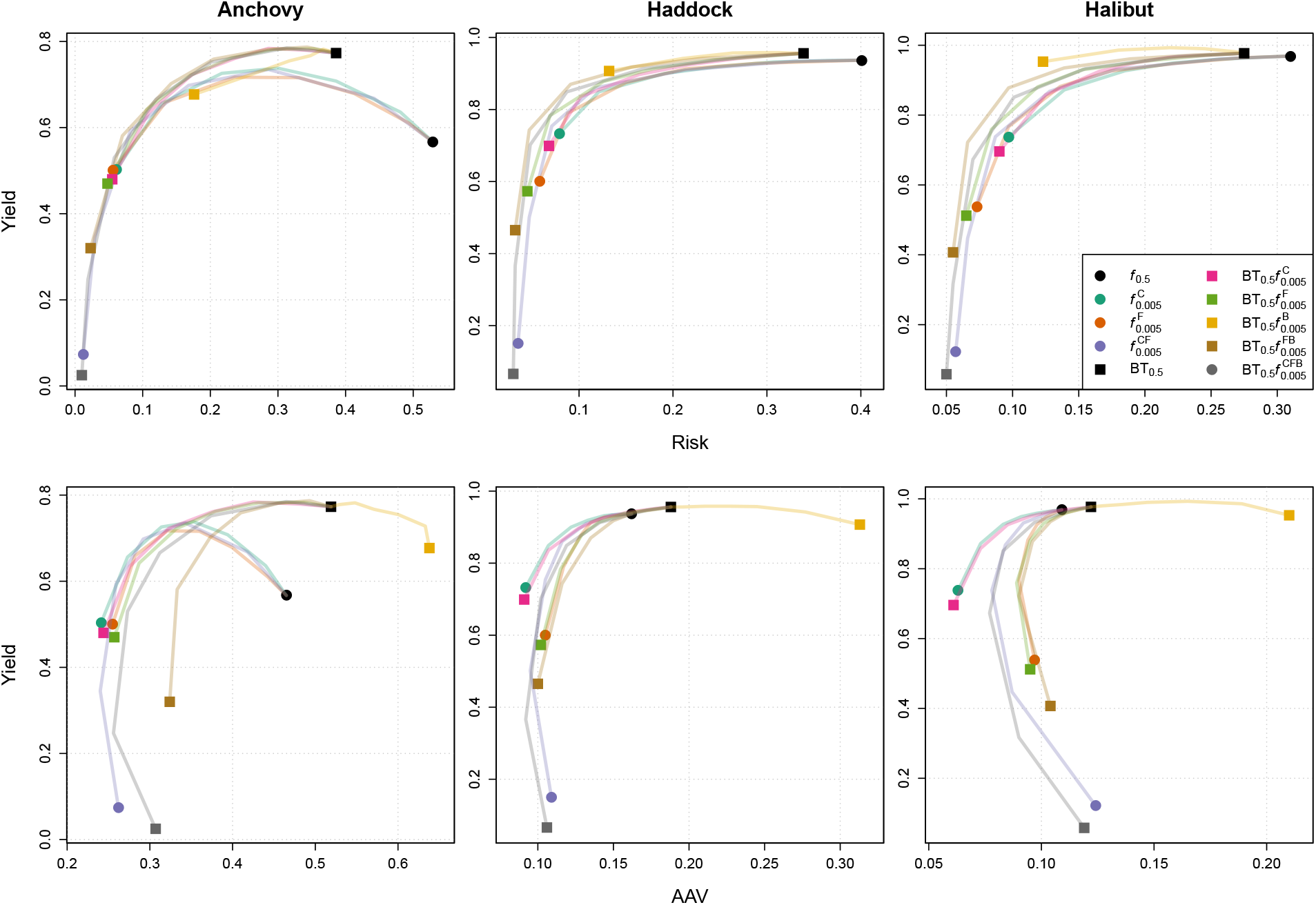
Trade-off between risk and yield (upper row) and between inter-annal absolute variability in yield (AAV) and yield (lower row) for all HCRs and anchovy, haddock, and halibut (columns) evaluated over the whole projection period (years 41-80, years 42-80 for AAV). Starting at the black diamond (MSY rule), the coloured lines connect HCRs without a biomass threshold and with decreasing risk fractiles. Similarly, starting at the black square (BT_0.5_ rule), the coloured lines connect HCRs with a biomass threshold and decreasing risk fractiles. The different symbols represent the HCR with the lowest risk fractile of 0.005. Considered risk fractiles are: 0.45, 0.35, 0.25, 0.15, 0.05, 0.005.

The consistent risk-yield trade-off for the fractile rules with and without threshold (excluding *f*^*B*^) confirm that the results from above, indicating that the threshold improves the efficiency of HCRs at the expense of larger variability in yield. Furthermore, the results reveal that the *P*\* method can be combined with threshold rule counteracting the increase in variability and leading to further risk reduction (Fig. 6).

### Sensitivity analysis

The results of the sensitivity analysis confirm that the risk-yield-variability trade-off is not sensitive to the Euler time step, the assumed prior distributions, or the assumptions in the intermediate year. Only the *f*_0.5_ rule for anchovy indicates slightly lower yield and higher risk with *dt*_Euler_ = 8 and when changing the assumptions in the intermediate year (e.g. assuming last year’s TAC). However, the qualitative results of the trade-off are not affected by that, i.e. the probability-based and threshold rules are still more efficient when assuming an intermediate year in which the model predicts the catch for the following year by assuming last year’s TAC or continuing the F process. Consequently, the results can be extrapolated to a management system that does not assume that the assessment takes place at the start of the year and just before the management period starts. Furthermore, the results indicate that the risk-yield trade-off is robust against implementation uncertainty (SD = 0.15). However, implementation uncertainty leads to higher variability in yield for all HCRs.

In contrast, the risk-yield-variability trade-off is sensitive to the level of process uncertainty, with lower levels leading to larger yields, lower risk levels, and lower variability in yield. These differences were the largest for anchovy. The length of the time series had the largest effect on the risk-yield-variability trade-off. Reducing the time series length from 40 to 20 years leads to higher risk and variability in yield for all species. Interestingly, the probability-based rules (e.g. 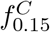 and 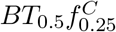) mitigate these negative effects, in particular, for haddock and halibut. For halibut, probability-based rules do not indicate any loss in yield and for haddock, they show a smaller increase in risk and loss in yield.

While the priors do not affect the trade-off, they have a large effect on the convergence rate, as does the time series length. For instance, removing all priors reduced the convergence rate from 66-86% to 2-14% for anchovy (19-33% for haddock and 26-35% for halibut) and reducing the time series length to 20 years shrunk the convergence rate from 82-85% to 3-4% for halibut (37-71% for anchovy and 21-27% for haddock; Supplementary Table C1). The low convergence rate can partly be attributed to the fact that we only used replicates with converged assessments in all years for the specific scenario and the baseline scenario combined. Considering the number of not converged assessments instead of simulations, the convergence rate was 46-98% across all scenarios and stocks (Supplementary Table C2).

For the assessment and management of the anchovy stock in the Bay of Biscay, several steepness parameters (h) of the stock-recruitment relationship are considered (*h* = 0.75 and *h* = 0.9; ICES 2019a). In this study, a higher steepness parameter (*h* = 0.9 instead of *h* = 0.75) implies a substantially larger yield and slightly lower risk for the MSY rule. For all other rules, it leads to slightly higher yield and risk (Supplementary Figure C4). The results indicate that the effect is mitigated by probability-based rules. The AAV is not affected substantially (Supplementary Table C6).

The results of the sensitivity analysis are presented in the Supplementary Section C.

## Discussion

We presented harvest control rules (HCRs) that incorporate the precautionary approach into fisheries management. We evaluated their performance for species with contrasting life history traits and under various levels of scientific uncertainty within an MSE framework. This study builds upon other studies evaluating the performance of HCRs in MSEs (e.g. Carruthers et al. 2014; Dankel et al. 2016; Dichmont et al. 2017; Walsh et al. 2018; Wetzel and Punt 2011; Wiedenmann et al. 2013).

### Biomass thresholds

The two precautionary approaches considered in this study, biomass thresholds and the *P*\* method, reduce the risk of overfishing and lead to faster stock recovery for over-exploited stocks. While the risk reduction is tightly linked to a short-term loss in yield, over the longer term, the probability-based rules lead to similar or even higher yields in comparison to deterministic rules. This finding is in line with previous studies demonstrating that biomass-based and probability-based control rules can maintain high average yield while reducing risk of low biomass (Benson et al. 2016; Irwin et al. 2008; Punt et al. 2008; Wiedenmann et al. 2017).

Threshold reference points, and in particular biomass thresholds, are an important component of the implementation of the precautionary approach into fisheries management (FAO 1995; Punt et al. 2008). Biomass thresholds reduce the risk of overfishing by reducing fishing mortality if biomass falls below a certain threshold. The F reduction is usually linearly to zero (or some lower fishing mortality rate), while the biomass at which *F* = 0 might vary. Overall, thresholds are effective in recovering over-exploited stocks and return high long-term yields as they recommend fishing at *F*_MSY_ if biomass is recovered (or high in general). On the downside and in particular for short-lived species, thresholds can lead to higher variability in recommended TAC and thus yield, with the largest variability for thresholds close to *B*_MSY_. Due to the variability in the recruitment success and dependent on the life history traits and gear selectivity, the stock will fluctuate around the target biomass level *B*_MSY_. If the biomass threshold (here *B*_trigger_) is defined at a level close to the target, the threshold is triggered more often than when defined by a smaller fraction of the target, leading to larger variability in TAC and yield.

If, on the other hand, the threshold is defined at a level much higher than *B*_MSY_, the target is triggered more often, however, the slope of the ascending part of the control rule is smaller, implying a lower variability in the target fishing mortality rate and thus the recommended TAC (Supplementary Figure A7). While from an ecological perspective, high variability in yield is not concerning and has in fact been shown to be a major component contributing to effective and adaptive management (Charles 1998), from a social and economical standpoint, it is more problematic for many reasons. For example, fishers might not have an alternative source of income and some running costs of the fishing vessels and production facilities are independent on the yield.

The results of this study confirm that ICES’s definition of the biomass threshold as *B*_trigger_ = 50%*B*_MSY_ for surplus production models (ICES 2017) is a meaningful compromise of the risk-yield-variability trade-off, which leads to higher long-term yield and lower risk, but slightly higher variability than the MSY rule without any threshold for all species. However, the results also indicate that higher thresholds could be considered which return similar yield while leading to further risk reduction. While thresholds defined as a fraction of *B*_MSY_ might fulfil one aspect of the precautionary approach, this definition does not account for the level of scientific uncertainty. Thus, rules with a threshold but without the *P*\* adjustment constitute deterministic rules that lead to higher risks with higher scientific uncertainty. In the case of high scientific uncertainty, Da-Rocha et al. (2016) argue that a higher biomass threshold should be used. While our results confirm that a higher threshold leads to more precautionary management in the face of high uncertainty, a higher threshold might affect the variability in yield substantially. Instead and as demonstrated, the *P*\* method can be combined with the threshold rules, resulting in effective and precautionary management. Alternatively, the threshold could also be defined in a probabilistic way, e.g. dependent on the estimated biomass distribution. This is for example being implemented for data-rich stocks within ICES, where *B*_trigger_ is defined as the lower 5th percentile of the spawning stock biomass distribution when fishing at *F*_MSY_ (ICES 2018).

### Deterministic vs. probability-based rules

In contrast to deterministic rules, such as fishing at *F*_MSY_, probability-based rules, such as the *P*\* method, account for scientific uncertainty in a quantitative and transparent manner. Probability-Based rules reduce the risk of overfishing without significant loss in long-term (here defined as 25 years after the first application of the rules) yield for species with different life-history traits. This finding is in line with recent simulation studies evaluating probability-based HCRs (Dankel et al. 2016; Wiedenmann et al. 2017). Over the short term (here defined as the first 5 years after start of the management), the risk reduction associated with probability-based rules is linked to an immediate loss in yield. While this relationship is almost proportional for medium- and long-lived species, our results indicate that the loss in short-term yield is less pronounced for short-lived species.

Interestingly, deterministic and probability-based rules with intermediate thresholds and risk fractiles showed higher long-term yields than the MSY rule (*f*_0.5_), in particular for the anchovy stock. This pattern was also described by Wiedenmann et al. (2017). We consider three factors that might have contributed to the higher yield of the more conservative rules: (i) varying recovery rates, (ii) the concept of pretty good yield, and (iii) the inadequacy of the MSY concept for stocks with large fluctuations in population size.

First, conservative rules recommend a low TAC thus leading to faster recovery of the over-exploited stocks to levels above *B*_MSY_ than less conservative rules. Dependent on the period considered for the calculation of the performance metrics, the faster stock recovery might lead to higher yields. The results indicate that the effect of faster recovery is more marked for longer-lived species: While the median biomass of the anchovy stock reaches levels above *B*_MSY_ within a few years after the first application of the MSY rule, it takes 10-20 years for haddock, and more than 40 years for halibut (upper row in Fig. 5). On the other hand, the more conservative rule 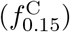 leads to a similar recovery rate for anchovy, but a much faster rate for haddock (5-10 years dependent on the scientific noise) and halibut (12-20 years; lower row in Fig. 5).

Secondly, many stocks show high yields across a wide range of population sizes and fishing mortality rates. For instance, Hilborn (2010) argues that a yield of 80% of MSY can be achieved over a wide range of biomass levels (20-50% of unfished biomass), a concept the author refers to as ‘pretty good yield’. For the stocks in this study, fishing mortality rates of 0.5 to around 2 times *F*_MSY_ lead to yield greater than 80% of MSY or more (dependent on the stock; Supplementary Figure A5). Thus, and in light of the concept of pretty good yield, it is not to be expected that conservative harvesting necessarily leads to a substantial loss in (long-term) yield.

Lastly, the MSY concept is derived from deterministic population models and is based on a constant harvest rate that does not take fluctuations in population size into account. For stocks with fluctuating population sizes, a HCR with a variable fishing mortality rate that follows recruitment success, might lead to a higher long-term yield than a constant harvest rate. The results of this study reveal that the biomass, fishing mortality rate, and yield are all strongly positively skewed (Supplementary Figures B11-B13), which can be attributed to the assumption of log-normally distributed deviations on the number of recruits in the operating model. The skewedness is greatest for the short-lived species (anchovy), which is characterised by few age classes (6) and a high gear selectivity for young age classes. Thus, the recruitment fluctuations have proportionally the largest effect on stock biomass, fishing mortality rate, and yield for this stock, which also shows the largest difference in long-term yield among HCRs. Due to the positively skewed distributions of those quantities, the median yield of the 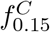 rule is closer to the true MSY of the operating model than the *f*_0.5_ rule even though the median fishing mortality rate of *f*_0.5_ is closer to the true *F/F*_MSY_ of the operating model (Supplementary Figures B12-B14). Assuming that the log-normally distributed recruitment deviations are realistic as indicated by meta-analyses (e.g. Thorson et al. 2014) and as assumed in almost all MSEs (e.g. Carruthers et al. 2014; Fischer et al. 2020; Wiedenmann et al. 2017), the results confirm findings of previous studies that for stocks with strongly fluctuating populations size, fishing at *F*_MSY_ is neither precautionary nor does it result in the highest yield (e.g. Lande et al. 2001). Theoretically, the pattern of higher long-term yield with more conservative rules could also be caused by biased estimates and stock status, i.e. fishing with a rate lower than the overestimated *F*_MSY_ is closer to the ‘true’ *F*_MSY_ (of the operating model) and thus returns a higher yield. However, in this study, we can exclude bias as a reason explaining the higher yield as the bias in the stock status is conservative for all stocks, i.e *F/F*_MSY_ is overestimated and *B/B*_MSY_ is underestimated (Supplementary Table B10).

Furthermore, probability-based rules lead to consistently low risk levels over a wide range of scientific uncertainty while the risk increases significantly with increasing scientific uncertainty for deterministic rules. This finding confirms the results by Dankel et al. (2016). We focused the main analysis on varying the observation uncertainty covering all levels assumed by Carruthers et al. (2014) and Wiedenmann et al. (2017) to evaluate the effect of increasing scientific uncertainty on the performance of the various rules. However, the information-content of the data (Bentley and Stokes 2009) is not only dependent on the level of observation uncertainty, but also on the quantity of available data and the contrast in the data in terms of periods of high and low biomass levels (Hilborn and Walters 1992). Reducing available data from a 40 year time series to 20 years decreases the performance of all rules. In this case, the shorter time series does not only reduce the number of data points, but also its contrast, as the shorter time series lacks information about the historical period of low exploitation rate and high stock biomass (Supplementary Figure A2). While the shorter time series leads to a larger proportion of non-converged assessments for all HCRs, for deterministic rules and in particular deterministic rules without a biomass threshold, it also leads to lower yield and higher risk. Probability-based rules mitigate the effect of higher scientific uncertainty in terms of lower quantity or quality of available information. Thus, probability-based rules have the natural incentive to reduce the scientific uncertainty, e.g. by improving data sampling programs (Punt and Donovan 2007a).

Process uncertainty, another component of scientific uncertainty, arising from natural variability, e.g. in the recruitment process, affects all HCRs, deterministic and probability-based, in the same way: Assuming independent log-normally distributed recruitment deviations with a SD of 0.15, leads to higher yield and lower risk. This can be attributed to the fact that process uncertainty leads to limited predictability of future states (Charles 2001). The results show that the effect of the process uncertainty depends on life-history, indicating that the effect is larger for shorter lived species. The population of a short-lived species is characterised by fewer cohorts and thus, higher fluctuations in the number of recruits (here the definition of higher process error) have a larger proportional effect on the dynamics of the total stock biomass.

The last component of scientific uncertainty, model uncertainty describes the incomplete knowledge of nature’s processes and states and in an MSE context, refers to the structural differences between the operating and assessment model. The operating model in this study is an age-based model while the assessment model is a surplus production model without any length or age structure. The results reveal that SPiCT underestimates *B/B*_MSY_ and overestimates *F/F*_MSY_, corresponding to a conservative bias, i.e. leading to more precautionary management actions. The bias is similar for the two quantities (relative F and B), the highest for the short-lived species (anchovy), and the lowest for long-lived species (halibut; Supplementary Table B10 and B11). While these results might indicate a general tendency for the relation of the accuracy of SPiCT estimates to the life history parameters, the biases are specific to the assumptions of this study and based only on three stocks. The biases can not only be attributed to the fundamental differences between the operating and estimation model (age-based vs. biomass-pool model, auto-correlated recruitment deviations in the operating model, fixed exploitation pattern vs. random walk process for F, density-dependence, etc.), but also to the limited amount of data available to the assessment and high observation uncertainty. Despite the conservative bias, all rules based on the SPiCT assessment lead to a relatively high maximum long-term yield corresponding to 95-98% of the yield achieved when fishing at constant true *F*_MSY_. Furthermore, we do not expect that the relative results of this study are affected by the bias, as all HCRs show similar biases. Small differences between HCRs might be due to the fact that more conservative rules lead to lower TACs and thus to greater contrast in the future biomass and thus more information content in the input data (Hilborn and Walters 1992). The results show that, while the bias of the SPiCT estimated stock status remains constant for different levels of scientific uncertainty, the precision increases with lower scientific uncertainty (Supplementary Table B10 and B11).

Another important source of uncertainty in fisheries management systems is implementation uncertainty, which represents uncertainty in the efficiency of management decisions that are designed to ensure that catch limits are not exceeded. This uncertainty might not only have large negative implications on the performance of the management strategy, but it is also generally underrepresented in fisheries related papers (< 0.06%; Fulton et al. 2011). Although we excluded implementation uncertainty in the main analysis to isolate the performance of the HCRs rather than the management of a specific stock, i.e. the actual catch corresponds to the recommended TAC, we explored the effect of unbiased log-normally distributed implementation noise (*SD* = 0.15) in the sensitivity analysis. Interestingly, the results do not indicate significant differences in the risk-yield trade-off (Supplementary Figures C1-C3). However, implementation uncertainty affects the variability in yield, with larger uncertainty leading to larger variability (13-53% higher variability; Supplementary Tables C3-C5).

The main disadvantage of probability-based HCRs is potentially forgone catch (Little et al. 2016), in particular given that the estimated stock status might be biased. Another potential disadvantage of more conservative rules described previously, could not be confirmed in this study. Specifically Wiedenmann et al. (2017) indicated that more conservative buffers (corresponding to smaller risk fractiles or higher thresholds in this study) lead to less frequent overfishing but when overfishing did occur the magnitude of it was higher. Our results indicate that overfishing occurs less frequently, and with a lower magnitude if it occurs, when using a conservative buffer (Supplementary Figure B3). This discrepancy could be due to the differences in the approaches, e.g. Wiedenmann et al. (2017) used a statistical catch-at-age (SCAA) model that differs markedly from the SPiCT model used here.

### Recommendations concerning probability-based

Ultimately, the definition of the probability-based rule and choice of the risk fractile (*P*\* value) is a policy decision and depends on management objectives as well as the managers’ risk tolerance. Ideally, and if expertise and resources allow, stock assessors should present the trade-off between risk, yield, and yield variability for a range of harvest control rules associated with probability-based rules with different threshold and risk fractiles to the managers. Nevertheless, based on the results from this study and in line with previous literature, a list of recommendations can be compiled guiding the selection and implementation of probability-based rules.

1. While generic MSEs are important and useful. The single best harvest control rule for a specific stock and given specific management objectives should in the best case be determined from a stock-specific MSE. The simulations should include scenarios with realistic levels of scientific uncertainty as this greatly affects the performance of the different HCRs (in particular deterministic vs. probability-based rules).
2. The probability-based rule should include both the *P*\* method and a biomass threshold. These ‘probability-based threshold rules’ are the more effective than rules without threshold or without the *P*\* method.
3. The probability-based rule should never lead to more risk-prone management decisions (prone to overfishing) than the deterministic alternative. This means for example that the risk fractile should not exceed 0.5 as mandated by the U.S. MagnussonStevens Fishery Conservation and Management Act and National Standards (Federal Register 2009).
4. The probability-based rule should recommend the catch limit (e.g. TAC) based on a specific risk fractile for the predicted catch distribution corresponding to the *f ^C^* and BT_0.5_*f ^C^* rules here. The results indicate that the risk-yield-variability trade-off is similar between HCRs with or without a biomass threshold and with risk fractiles on different quantities. Due to the high variability in yield associated with the risk fractile on the biomass relative to the threshold (here: BT_0.5_*f*^*B*^), we do not recommend to account for the uncertainty of the quantity *B/B*_trigger_.
5. The choice of the risk fractile should depend on the quantity (quantities) it is applied to. For the *f*^C^ and 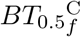 rules, we recommend use of a risk fractile in the range of 0.15 to 0.45. Risk fractiles in this range reduce the risk by up to 55-59% (for anchovy; 52% for haddock and 40% for halibut) and the variability in yield by up to 30-35% (for anchovy; 8-9% for haddock and 18-24% for halibut) without a detrimental loss in expected yield in comparison to corresponding rules using the median (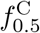 and 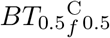; Supplementary Table B7-B9). In fact, for anchovy, rules with fractiles in this range lead to an increase in expected yield by up to 28% (for 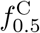). For the other rule and stocks, the loss in expected yield is only 4-6%. These values are based on the metrics corresponding to the entire management period for all species and HCRs. This recommended range of the risk fractiles is in line with the recommendation by Prager and Shertzer (2010), who suggested a range of 0.25-0.45 and the recommendation by Council and Service (2016), which restricts the risk fractile from exceeding 0.45. Our results indicate that very small risk fractiles (< 0.005) are too restrictive and can lead to sharp increases in the variability in yield. As stated in the above, the best performing risk fractile should be determined by stock-specific MSEs and in agreement with all stake-holders. However, we acknowledge the benefit of probability-based rules and that it might not always be possible to apply stock-specific MSEs. This might for example be due to a lack of expertise, information, or resources. For these cases exclusively and until available data indicate otherwise, we recommend use of a probability-based HCR with a biomass threshold equal to 0.5*B*_MSY_ and the 0.35 risk fractile for the predicted catch distribution 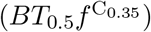.

Note that these recommendations are based on MSEs using SPiCT as the assessment model and three different operating models. While we expect our conclusions and recommendations to be valid for other combinations of assessment model and stock dynamics, further research is needed to confirm this.

### Future research

This study did not consider a limit reference point in terms of biomass (*B*_lim_) in the HCRs. A biomass limit reference point implies that fishing is terminated if biomass falls below the reference biomass. For instance, the ‘40-10 rule’ is a popular HCR with a biomass threshold and limit reference point, implemented e.g. in Canada (DFO 2009). The rule involves reducing the fishing mortality rate linearly to zero when the biomass is below 40% and above 10% of the virgin biomass and to terminate fishing if the biomass is below 10% of the virgin biomass (Punt and Ralston 2007). Future research should investigate the implications of the inclusion of *B*_lim_ in probability-based rules. Furthermore, the performance of a probabilistic definition of the biomass threshold or limit point should be evaluated.

In this study, we excluded all non-converged SPiCT assessments to isolate the performance of the actual HCRs rather than constant catch rules (the alternative HCR). However, non-convergence of assessment models (incl. SPiCT) is a common issue in fisheries management and as relevant as alternative approaches are not necessarily more precautionary (ICES 2013; Punt et al. 2020). Albeit, the proportion of replicates where all SPiCT assessments converged was relatively high (around 80%), some scenarios showed a low convergence rate of only 3-51%. In particular, the scenario with the shorter time series and the scenario without priors led to low convergence rates. Future work should thus be allocated to explore alternative applications of the method, e.g. trend-based rules, and the implications of modifications to the assessment model, such as fixing the shape of the production curve, which might help to increase the convergence rate.

Lastly, future work should explore the bias in the stock status estimated by SPiCT. It would be interesting to know if other applications confirm the conservative bias and how the bias is related to the life history traits of the species.

## Conclusion

Identifying harvest control rules that are robust to uncertainty is essential for precautionary and effective fisheries management. This study showed that biomass thresholds as well as the *P*\* method reduce the risk of overfishing without loss in long-term yield. In contrast to deterministic rules, probability-based rules (e.g. the *P*\* method) show similar or lower risk levels while maintaining high long-term yields in the face of high scientific uncertainty. This study provides guidelines for the definition and implementation of probability-based rules and concludes that the most efficient and robust HCR should include both a biomass threshold as well as the *P*\* method with a risk fractile (*P*\* value) between 0.15 and 0.45 for the distribution of the predicted catch. Furthermore, for stocks exhibiting large fluctuations in population size, the results of this study support the notion that the MSY related reference points *F*_MSY_ and *B*_MSY_ are not precautionary nor efficient as target reference points and should be treated as limit or threshold reference points (Mace and Mace 2001; Wiedenmann et al. 2017) or combined with precautionary approaches as presented in this study. Our MSE framework overcomes some of the limitations of other frameworks and can directly be used for stock-specific MSE, which will remain a crucial component of precautionary fisheries management in the future. The results of this study indicate ways to make current management strategies more robust and incorporate the precautionary approach into fisheries management.

## Acknowledgements

This work was mainly funded by the EMFF project “ManDaLiS - Improving the management basis for Danish data-limited stocks” (Ref 33113-B-16-085), which is funded by the European Maritime and Fisheries Fund and the Danish Fisheries Agency. TKM received financial support from the University of Washington School of Aquatic and Fishery Sciences, the Otto Mønsted Foundation, and the Idella Foundation. The authors would like to express their gratitude to all participants of the ICES workshops “WKLife VII - IX” for comments on the methodology and interpretation of the results of this study during the workshops.

## Data Availability Statement

The code for the operating model is available at https://github.com/tokami/mse. The code for the assessment model is available at https://github.com/tokami/spict/tree/probHCR. Additional R scripts supporting this study are available at https://github.com/tokami/pubs/tree/master/probHCR.

