## Supplementary Information for "Implementing the precautionary approach into fisheries management: Making the case for probability-based harvest control rules"

### Supplementary Section A: Methodology

This section includes all equations governing the population dynamics in the operating model, more information about the assumptions of the simulation study, the assessment model, and harvest control rules, as well as a table with all scenarios of this study.

#### Operating model

We used an age-structured operating model with annual time steps to simulate the population dynamics. The equations governing the population dynamics are presented in the following:

Growth in length is modelled by means of the von Bertalanffy growth function (Bertalanffy 1938):

$$L_a = L_\infty(1 - \exp(-k(a - a_0))) \tag{A1}$$

where $L_\infty$ is the asymptotic length in cm, $k$ is the growth rate in $yr^{-1}$, and $a_0$ is the age at length = 0cm. Weight at age is calculated by the power law:

$$w_a = aL_a^b \tag{A2}$$

with the two parameters $a$ in g/cm and unit-less $b$. Mortality at length is calculated by means of the length-based empirical formula by Gislason et al. 2010:

$$M_L = \exp(0.55 - 1.61\ln(L_a) + 1.44\ln(L_\infty) + \ln(k)) \tag{A3}$$

where $M_L$ for $L \in [0, 10]$ is equal to $\exp(0.55 - 1.61\ln(10) + 1.44\ln(L_\infty) + \ln(k))$, accounting for the low sample size of fish under 10cm in Gislason's meta study (Gislason et al. 2010). Maturity at length is modelled by the logistic function:

$$m_L = \frac{1}{1 + \exp(-\ln(19)\frac{L - L_{m50}}{L_{m95} - L_{m50}})} \tag{A4}$$

where $\mathrm{L_{m50}}$ and $\mathrm{L_{m95}}$ correspond to the length in cm where 50% and 95% of the individuals, respectively, are mature. Equally, selectivity at length is modelled by:

$$\zeta_L = \frac{1}{1 + \exp(-\ln(19)\frac{L - L_{s50}}{L_{s95} - L_{s50}})} \tag{A5}$$

where $L_{s50}$ and $L_{s95}$ correspond to the lengths in cm at which the probability of capture is 50% and 95%, respectively.

Accounting for fish growth within a year, these length-based processes (length, weight, natural mortality, maturity, and selectivity) were converted into age-based processes by season by means of a stochastic seasonal age-length key. The key defines the proportion of fish of a certain length contributing to the different age groups per season (Rudd and Thorson 2018):

$$p_{l,a,s} = \begin{cases} \phi(\frac{l-L_{a,s}}{L_{a,s}CV_L}) & \text{for } l = 1 \\ \phi(\frac{l-L_{a,s}}{L_{a,s}CV_L}) - \phi(\frac{l-1-L_{a,s}}{L_{a,s}CV_L}) & \text{for } 1 < l < L \\ 1 - \phi(\frac{l-1-L_{a,s}}{L_{a,s}CV_L}) & \text{for } l = L \end{cases} \tag{A6}$$

where $L_{a,s}$ is the length at age a in season s, $l \in [1, L]$ are the mid lengths of the defined length bins, and $CV_L$ is the coefficient of variation per length.

The population dynamics are governed by:

$$N_{a,y,s} = \begin{cases} R_y & \text{for } a = 0, y \geq 1, s = 1 \\ N_{a-1,y,s}\exp(-M_{a-1,y,s} - F_y\zeta_{a-1,s}) & \text{for } 0 < a < A, y = 1, s = 1 \\ \frac{N_{a-1,y,s}\exp(-M_{a-1,y,s} - F_y\zeta_{a-1,s})}{1-\exp(-M_{a-1,y,s} - F_y\zeta_{a-1,s})} & \text{for } a = A, y = 1, s = 1 \\ N_{a,y-1,s}\exp(-M_{a,y,s-1} - F_{y,s-1}\zeta_{a-1,s-1}) & \text{for } a < A, y \geq 1, 1 < s \leq S \\ N_{a-1,y-1,S}\exp(-M_{a-1,y-1,S} - F_{y-1,S}\zeta_{a-1,S}) & \text{for } 0 < a < A, y > 1, s = 1 \\ (N_{a-1,y,s-1} + N_{a,y,s-1})\exp(-M_{a-1,y,s-1} - F_{y,s-1}\zeta_{a-1,s-1}) & \text{for } a = A, y > 1, s = 1 \end{cases} \tag{A7}$$

where $R_y$ are the number of recruits based on the Beverton-Holt stock recruitment relationship (Beverton and Holt 1957) parameterised with the steepness parameter h:

$$R_y = \frac{4hR_0SSB_{y,s-1}}{SSB_0(1-h) + SSB_{y,s-1}(5h-1)} * \exp(\tau_y^R) \tag{A8}$$

where $SSB_0$ is the unfished spawning stock biomass:

$$SSB_0 = \sum_{a=0}^{A} R_0 \exp(-\sum_{a=0}^{a} M_a) w_a m_a \tag{A9}$$

and $\tau_y^R$ are auto-correlated recruitment deviations (Thorson et al. 2014):

$$\tau_y^R = \begin{cases} \epsilon_y^R & \text{for } y = 1 \\ \rho\tau_{y-1}^R + \sqrt(1-\rho^2)\epsilon_y^R & \text{for } y > 1 \end{cases} \tag{A10}$$

where $\epsilon_{Ry} \sim N(-\frac{\sigma_R^2}{2}, \sigma_{\mathrm{R}}{}^2)$ are biased-corrected recruitment deviations. The catch in numbers by age, year and season is calculated by:

$$C_{a,y,s} = \frac{F_y\zeta_{a,s}}{(M + F_y\zeta_{a,s})}\, N_{a,y,s}\, (1 - \exp(-M - F_y\zeta_{a,s}))) \tag{A11}$$

Annual catch in weight $S^w$ is defined by:

$$\mathrm{C}_y^w = \sum_{a=0}^{A} C_{a,y,s} w_{a,s} \tag{A12}$$

Total stock biomass $B$ is defined by:

$$\mathrm{B}_y = \sum_{a=0}^{A} N_{a,y,s} w_{a,s} \tag{A13}$$

Spawning stock biomass is defined by:

$$\text{SSB}_t = \sum_{a=0}^{A} N_{a,y,s} w_{a,s} m_{a,s} \tag{A14}$$

Annual catch observations are calculated with:

$$\text{Cobs}_y = \text{C}_y^w \epsilon_y^C \tag{A15}$$

where $\epsilon_C \sim \text{Lognormal}(0, \sigma_C^2)$ is the catch observation noise. Survey observations can be computed independent of seasons by:

$$I_t = q \sum_{a=0}^{A} \exp(\ln(N_{a,y,s}) - (M_{a,s} + F_y \zeta_{a,s})\delta_t) w_{a,s} \zeta_{a,s} * \epsilon_y^I \tag{A16}$$

where t defines any time point independent of seasons, $\delta_t$ is the time period from the closest season to the time point t, q is the catchability coefficient, and $\epsilon_I \sim \text{Lognormal}(0, \sigma_I^2)$ describes the survey observation noise.

Natural mortality at length was calculated using the empirical formula by Gislason et al. (2010) and the growth parameters of each stock. The length-based processes, i.e. natural mortality, maturity, weight at length, and gear selectivity (Fig. A1), were transferred to corresponding ages at each time step assuming a stochastic age-length key as described by Rudd and Thorson (2018) with a bin size of 1 cm and a coefficient of variation of 10% (Fig. A1). For all stocks, the reparameterised Beverton and Holt stock-recruitment relationship was assumed, where steepness represents the fraction of unfished recruitment that results when the spawning biomass is reduced to 20% of the unfished level (Beverton and Holt 1957; Mace and Doonan 1988). Recruitment at virgin biomass ($R_0$) of $1e6$ and an age of recruitment to the population of zero was assumed. Spawning was assumed to occur at the beginning of each year.

We estimated stochastic reference points for each stock based on optimising long-term surplus production over a range of fishing mortality rates assuming process uncertainty (Fig. A4) and defined the biomass limit reference point $B_{\text{lim}}$ as 20% of the virgin biomass (Dichmont et al. 2017).

The Total Allowable Catch (TAC) resulting from the assessment model in combination with the HCR was removed from the population given that enough biomass was available (the upper limit of the search space for the fishing mortality rate was $5\text{yr}^{-1}$ for all species). If the TAC exceeded the vulnerable biomass at the mid of the next time step, the TAC was reduced to 75% of the exploitable biomass at the midpoint of the time step. The catchability of both simulated surveys was set to 0.05.

Table A1: References for life history parameter values for the dab, haddock, and Greenland halibut stocks.

| Parameter | Description | Stocks | | |
|---|---|---|---|---|
| | | **Dab** | **Haddock** | **Greenland halibut** |
| $L_\infty$ | Asymptotic length [cm] | Jardim et al. 2015 | ICES 2017 | Jardim et al. 2015 |
| $K$ | Growth rate [1/yr] | Jardim et al. 2015 | ICES 2017 | Jardim et al. 2015 |
| $a_0$ | Age at length=0 [yr] | Jardim et al. 2015 | ICES 2017 | Jardim et al. 2015 |
| $a$ | Length-weight scalar [g/cm] | ICES 2019b | ICES 2019a | Jardim et al. 2015 |
| $b$ | Length-weight exponent | ICES 2019b | ICES 2019a | Jardim et al. 2015 |
| $a_{max}$ | Maximum age [yr] | Froese and Pauly 2019 | Froese and Pauly 2019 | Froese and Pauly 2019 |
| $h$ | Steepness | Myers et al. 1999 | Myers et al. 1999 | Myers et al. 1999 |
| $L_{m50}$ | Length at 50% maturity [cm] | ICES 2019b | ICES 2017 | Rickman et al. 2000 |
| $L_{m95}$ | Length at 95% maturity [cm] | ICES 2019b[1] | ICES 2017 | Rickman et al. 2000 |
| $L_{s50}$ | Length at 50% selectivity [cm] | ICES 2019b[2] | ICES 2019a[3] | ICES 2013 |
| $L_{s95}$ | Length at 95% selectivity [cm] | ICES 2019b[2] | ICES 2019a[3] | ICES 2013 |
| $\sigma_R$ | SD of the recruitment deviations | Thorson et al. 2014 | Thorson et al. 2014 | Thorson et al. 2014 |
| $\rho_R$ | Coeffcient of auto-correlated recruitment deviations | Thorson et al. 2014 | Thorson et al. 2014 | Thorson et al. 2014 |

[1] $L_{m95}$ for dab was calculated from $L_{m50}$ and width of maturity ogive.
[2] $L_{s50}$ and $L_{s95}$ for dab were not given directly, but set based on the information that individuals smaller than 12cm were not counted to explitable biomass in spict assessment in (ICES 2019b).
[3] $L_{s50}$ and $L_{s95}$ for haddock were not given directly, but inferred from the fishing mortality at age given in (p. 295 in ICES 2019a).
[4] $L_{s95}$ for Greenland halibut was not given directly, but inferred from the graph depicting probability of capture vs length in (ICES 2013).

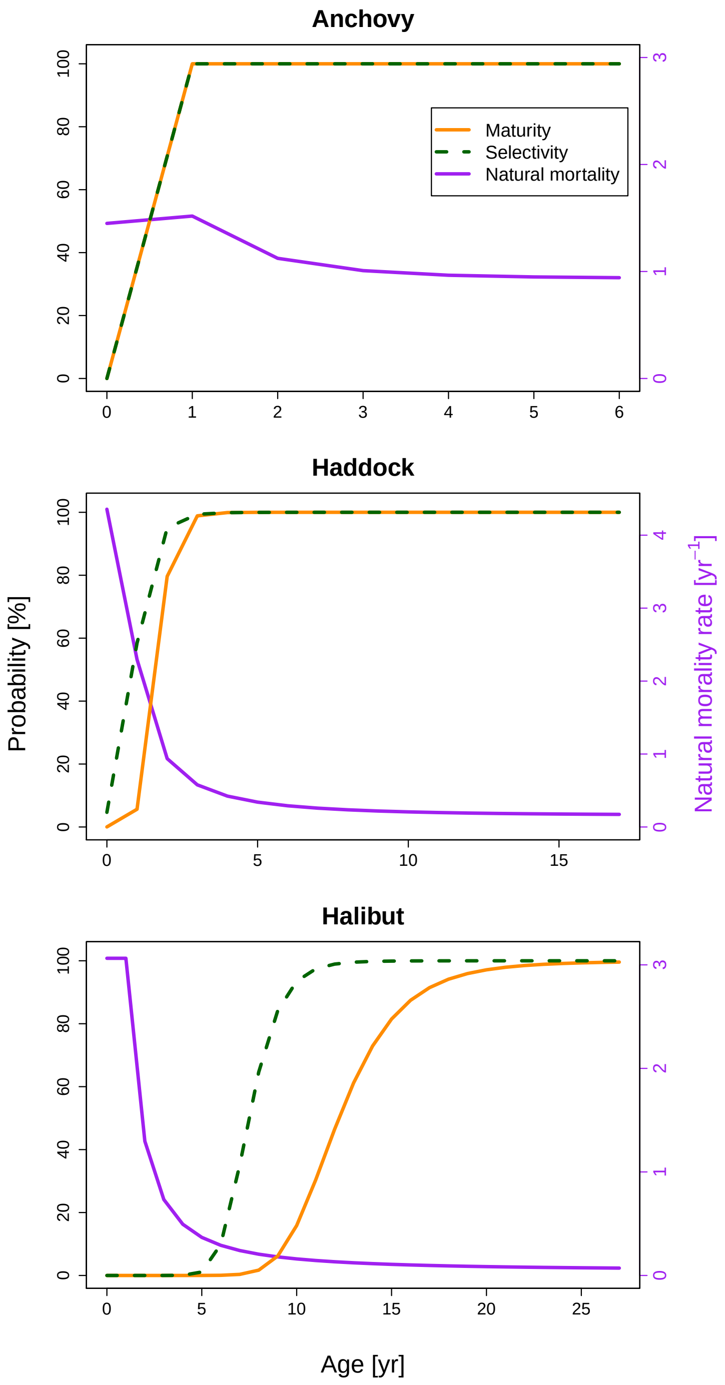


Figure A1: Maturity, gear selectivity, and natural mortality by age for all species.

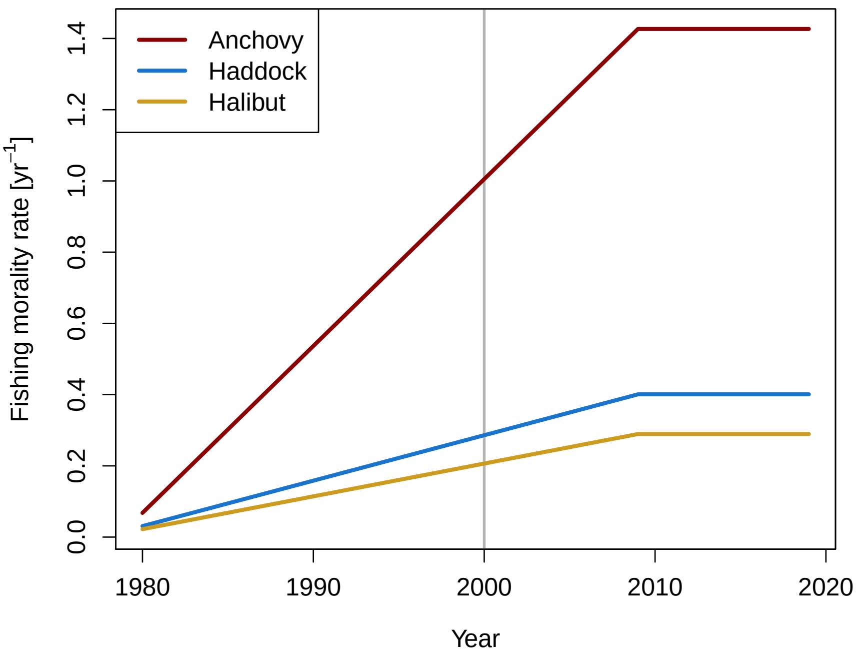


Figure A2: Fishing mortality rate (F) during the 40 historical years for the three species. The grey vertical line indicates where the time series was cut for scenario 14 (scenario with time series of 20yr), i.e. the historical F pattern for scenario 14 covers the period from year 2000 to 2019.

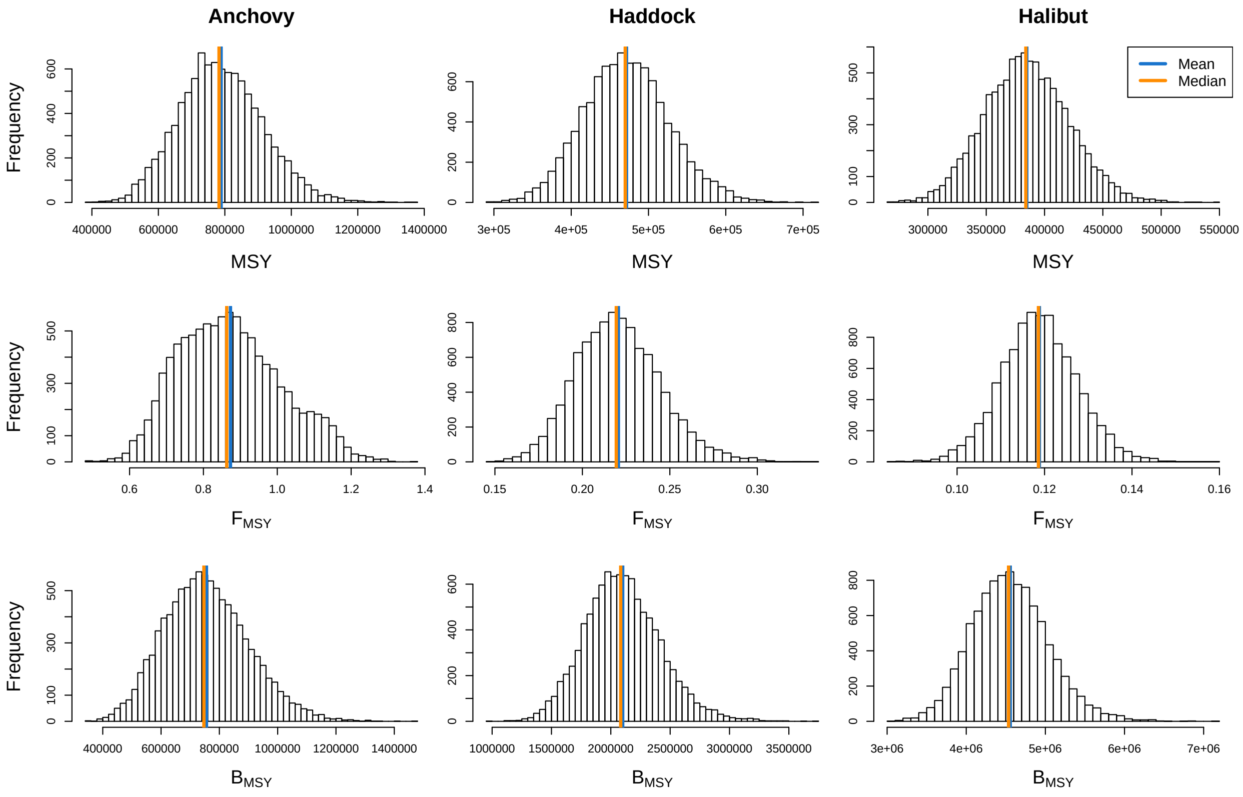


Figure A3: Posterior distributions of the simulated reference points (MSY, $F_{\mathrm{MSY}}$, $B_{\mathrm{MSY}}$) for all species. The two vertical lines represent the mean (solid) and median (dashed).

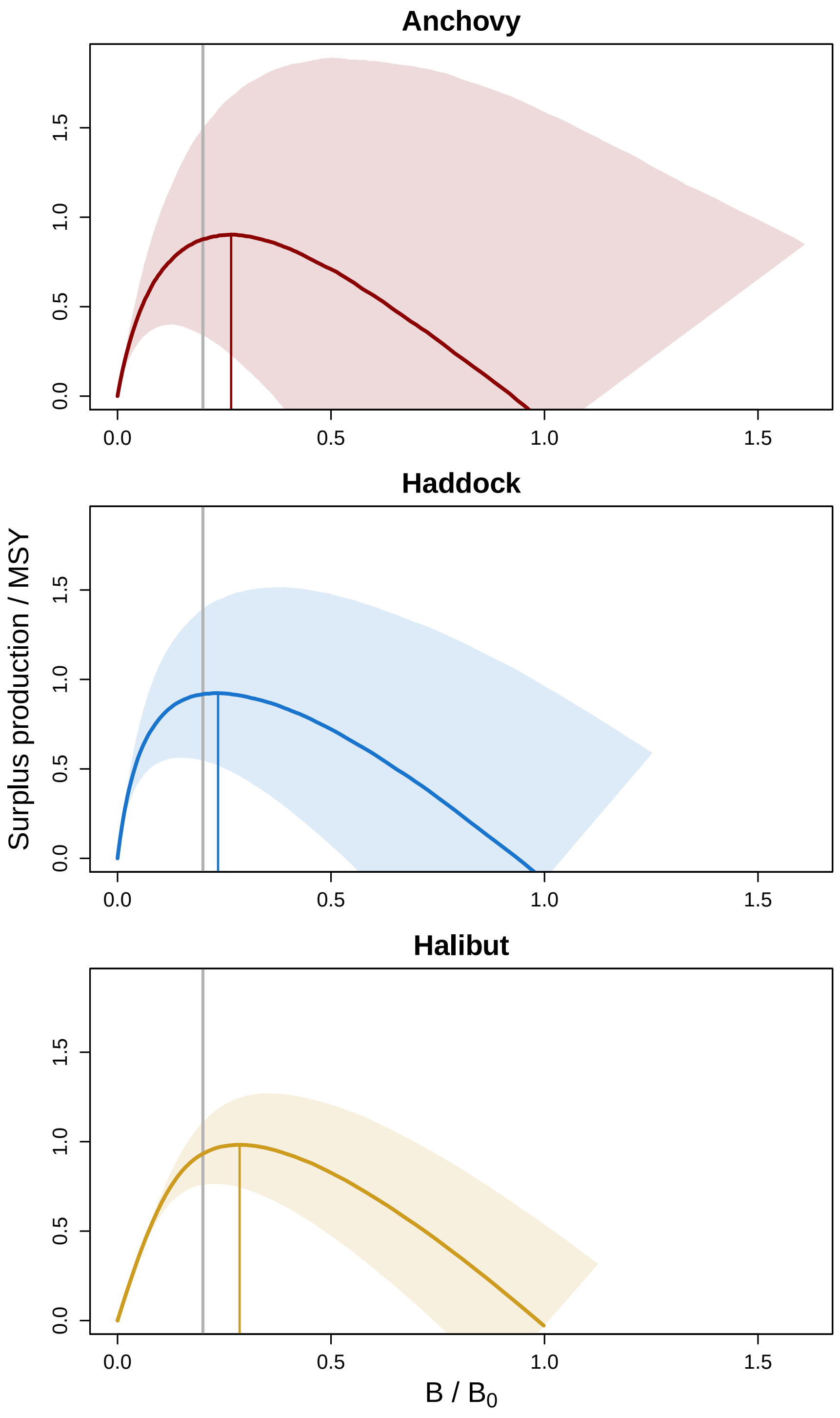


Figure A4: Total stock biomass relative to virgin biomass ($B_0$) against the surplus production relative to MSY for anchovy, haddock, and halibut. The solid lines represent the median relationship and the area extends from the 10th to the 90th percentile. The vertical lines indicate the total stock biomass (B) at which the stock shows the highest surplus production ($B_{\mathrm{MSY}}/B_0$).

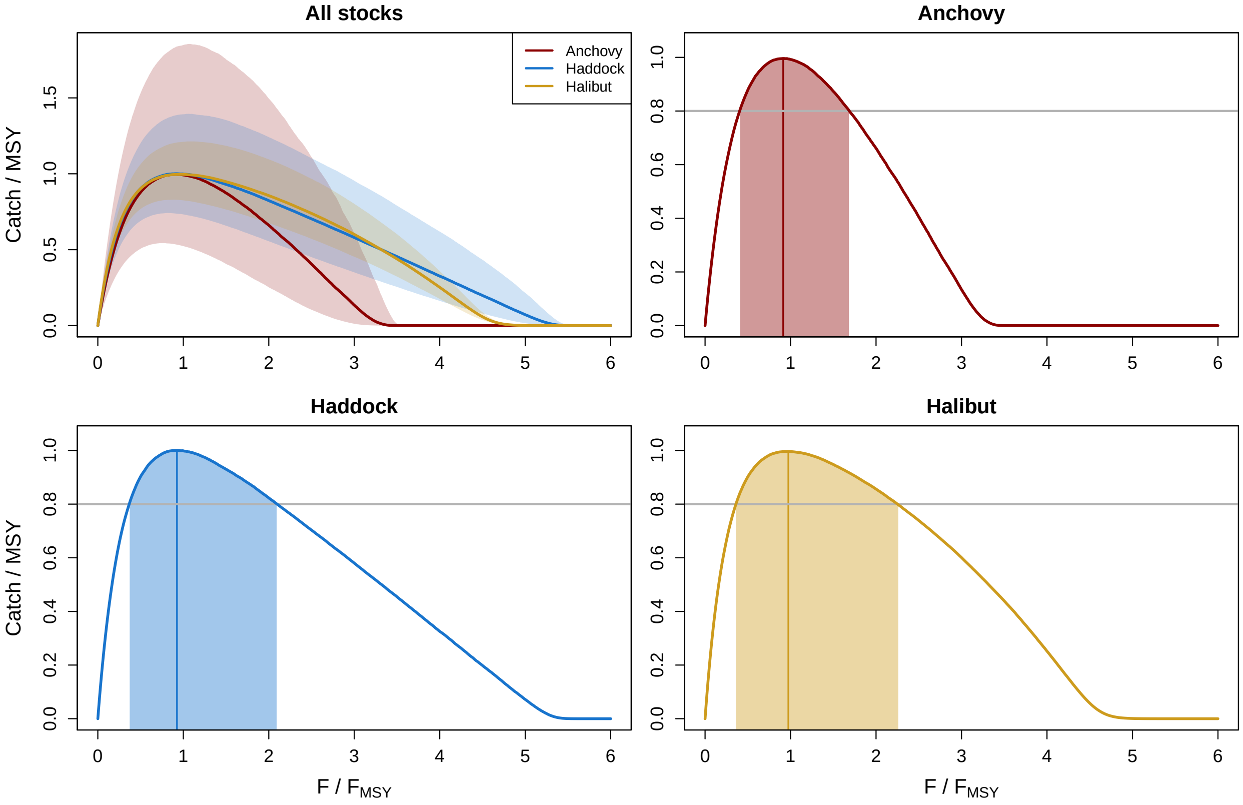


Figure A5: Fishing mortality rate relative to $F_{\mathrm{MSY}}$ against catch relative to MSY for all stocks together and individually. The solid lines represent the median and the shaded area in the plot with all species extends from the 10th and 90th percentile. Shaded area in the plots indicate fishing mortality rate which leads to yields of at least 80% of MSY (grey horizonatal line).

Table A2: Equations of the Stochastic surplus Production model in Continuous Time (SPiCT). For a description of the parameters, please refer to Table A3; for more information, please refer to Pedersen and Berg 2017.

| No | Equation | Description |
|---|---|---|
| 1 | $dB_t = \left(\gamma\, m \frac{B_t}{K} - \gamma\, m \left[\frac{B_t}{K}\right]^n F_t\, B_t\right) dt + \sigma_B\, B_t\, dW_t$ | Biomass process |
| 2 | $F_t = S_t\, G_t \exp(H_{j(t)})$ | Fishing mortality (F) process |
| 3 | $d \log(G_t) = \sigma_F\, dV_t$ | Diffusion component of the F process |
| 4 | $S_t = \exp(D_{s(t)})$ | Seasonal component of the F process |
| 5 | $I_t = q\, B_t \cdot e^{\nu_t}$ | Abudance index observations |
| 6 | $C_t = \int_t^{t+\Delta} F_s\, B_s\, ds \cdot e^{\varepsilon_t}$ | Catch observations |

Table A3: Description of the model parameters of the Stochastic surplus Production model in Continuous Time (SPiCT). The number of estimated parameters in this study adds up to 9 including the 8 parameters in the table plus an additional catchability coefficient for the second abundance index. For more information, please refer to Pedersen and Berg 2017.

| Parameter | Description | Estimated? |
|---|---|---|
| $K$ | Carrying capacity | Yes |
| $m$ | Productivity parameter (= MSY) | Yes |
| $n$ | Shape parameter of the production curve | Yes |
| $\gamma$ | Gamma $\gamma = n^{n/(n-1)}/(n-1)$ | No |
| $W_t$ | Brownian motion of the biomass process | No |
| $\sigma_B$ | Standard deviation of biomass process noise | Yes |
| $V_t$ | Brownian motion of the fishing mortality (F) process | No |
| $\sigma_F$ | Standard deviation of F process noise | Yes |
| $D_{s(t)}$ | Cyclic B-spline with a period of 1 year | No |
| $s(t)$ | Mapping from t to the proportion of the current year that has passed | No |
| $q$ | Catchability for each abundance index | Yes |
| $\nu_t$ | Index observation errors $\nu_t \sim N(0, \sigma_I^2)$ | No |
| $\sigma_I$ | Standard deviation of the index observation error | Yes |
| $\Delta_t$ | Time interval length of catch observations (typically a year or quarter of a year) | No |
| $\varepsilon_t$ | Catch observation errors $\varepsilon_t \sim N(0, \sigma_C^2)$ | No |
| $\sigma_C$ | Standard deviation of the catch observation error | Yes |

The default model configuration of SPiCT includes three vague priors on the shape parameter of the production curve $log(n) \sim$ N(2,2)\$ and on two hyper parameters $log(\alpha) \sim N(1,2)$ and \$log($\beta$) $\sim$ N(1,2). The prior on $log(n)$ corresponds to the symmetrical surplus production model (Schaefer 1954). The hyper parameters are the ratios of the standard deviations (SD) of the observation to process noise terms: $log(\alpha) = log(\sigma_I) - log(\sigma_B)$ and $log(\beta) = log(\sigma_F) - log(\sigma_C)$ (c.f. Table A2 and A3). The priors correspond to equal SDs of the observation and process noise terms for the catches and indices respectively (Pedersen and Berg 2017).

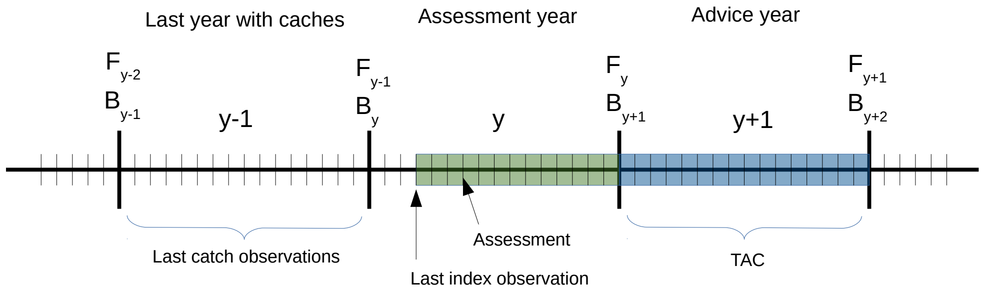


Figure A6: A timeline defining the 'continuous' time quantities of SPiCT in relation to the discrete time of assessment and advice years in fisheries management. The small vertical bars represent the time steps of the Forward Euler scheme (in this graph: 16 time steps per year). The green area depicts the projection period between the last observation (here: index observation) and the start of the management period. The blue area depicts the period for which the TAC is going to be calculated

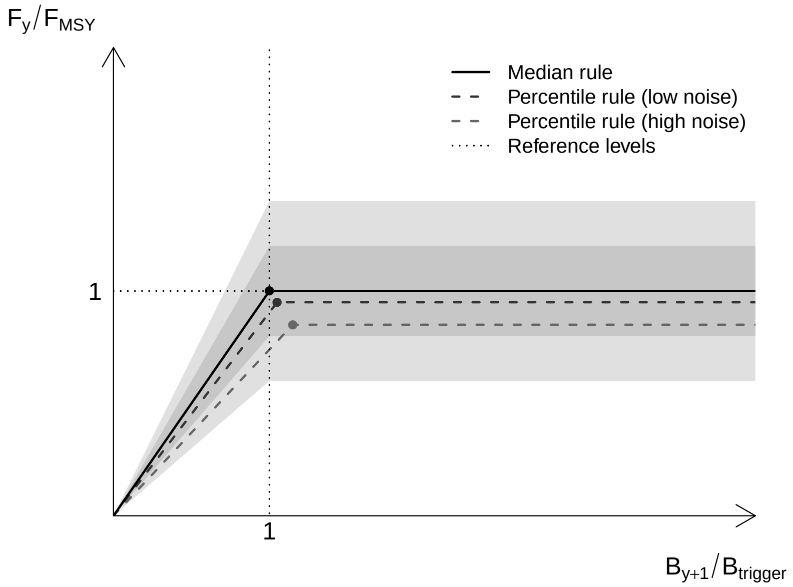


Figure A7: Target fishing mortality rate based on the hockey-stick MSY harvest control rule (HCR; ICES 2018): Recommended total allowable catch (TAC) is equal to the predicted catch with $F = F_{\text{MSY}}$ for $B \geq B_{trigger}$ and F linearly decreased to 0 for $B \rightarrow 0$. The grey rectangles depict the uncertainty around the relative state variables $B_{y+1}/B_{trigger}$ and $F_y/F_{\text{MSY}}$. The dark-grey dashed line illustrates the HCR as defined with uncertainty buffers $< 0.5$ for the relative state variables. While the median rule (black solid line) remains unchanged with increasing scientific uncertainty, the probability-based rules lead to a more risk averse trajectory (grey dashed lines).

Table A4: The code and definition of all HCRs used in this study.

| **Code** | **True** $F/F_{MSY}$ | **Biomass threshold** | **Risk fractile for** $C_{y+1}$ | **Risk fractile for** $F_y/F_{\mathbf{MSY}}$ | **Risk fractile for** $B_{y+1}/B_{\mathbf{trigger}}$ |
|---|---|---|---|---|---|
| Reference rules | | | | | |
| $F = 0$ | 0% | | | | |
| $F = F_{\text{MSY}}$ | 100% | | | | |
| Deterministic rules | | | | | |
| $f_{0.5}$ | | 0 | 0.5 | 0.5 | 0.5 |
| $\text{BT}_{0.25}$ | | 0.25 | 0.5 | 0.5 | 0.5 |
| $\text{BT}_{0.5}$ | | 0.5 | 0.5 | 0.5 | 0.5 |
| $\text{BT}_{0.75}$ | | 0.75 | 0.5 | 0.5 | 0.5 |
| $\text{BT}_{1}$ | | 1 | 0.5 | 0.5 | 0.5 |
| $\text{BT}_{1.25}$ | | 1.25 | 0.5 | 0.5 | 0.5 |
| $\text{BT}_{1.5}$ | | 1.5 | 0.5 | 0.5 | 0.5 |
| $\text{BT}_{1.75}$ | | 1.75 | 0.5 | 0.5 | 0.5 |
| $\text{BT}_{2}$ | | 2 | 0.5 | 0.5 | 0.5 |
| Probability-based rules | | | | | |
| $f^{\text{C}}$ rules | | | | | |
| $f^{\text{C}}_{0.45}$ | | | 0.45 | 0.5 | |

*Continue on the next page*

Table A4: All 69 Harvest Control rules (HCRs) used in this study (cont.).

| **Code** | **True** $F/F_{\mathbf{MSY}}$ | **Biomass threshold** | **Risk fractile for** $C_{y+1}$ | **Risk fractile for** $F_y/F_{\mathbf{MSY}}$ | **Risk fractile for** $B_{y+1}/B_{\mathbf{trigger}}$ |
|---|---|---|---|---|---|
| $f^{\mathrm{C}}_{0.35}$ | | | 0.35 | 0.5 | |
| $f^{\mathrm{C}}_{0.25}$ | | | 0.25 | 0.5 | |
| $f^{\mathrm{C}}_{0.15}$ | | | 0.15 | 0.5 | |
| $f^{\mathrm{C}}_{0.05}$ | | | 0.05 | 0.5 | |
| $f^{\mathrm{C}}_{0.005}$ | | | 0.005 | 0.5 | |
| $f^{\mathrm{F}}$ rules | | | | | |
| $f^{\mathrm{F}}_{0.45}$ | | | 0.5 | 0.45 | |
| $f^{\mathrm{F}}_{0.35}$ | | | 0.5 | 0.35 | |
| $f^{\mathrm{F}}_{0.25}$ | | | 0.5 | 0.25 | |
| $f^{\mathrm{F}}_{0.15}$ | | | 0.5 | 0.15 | |
| $f^{\mathrm{F}}_{0.05}$ | | | 0.5 | 0.05 | |
| $f^{\mathrm{F}}_{0.005}$ | | | 0.5 | 0.005 | |
| $f^{\mathrm{CF}}$ rules | | | | | |
| $f^{\mathrm{CF}}_{0.45}$ | | | 0.45 | 0.45 | |
| $f^{\mathrm{CF}}_{0.35}$ | | | 0.35 | 0.35 | |
| $f^{\mathrm{CF}}_{0.25}$ | | | 0.25 | 0.25 | |
| $f^{\mathrm{CF}}_{0.15}$ | | | 0.15 | 0.15 | |
| $f^{\mathrm{CF}}_{0.05}$ | | | 0.05 | 0.05 | |
| $f^{\mathrm{CF}}_{0.005}$ | | | 0.005 | 0.005 | |
| $\mathrm{BT}_{0.5}f^{\mathrm{C}}$ rules | | | | | |
| $\mathrm{BT}_{0.5}f^{\mathrm{C}}_{0.45}$ | | 0.5 | 0.45 | 0.5 | 0.5 |
| $\mathrm{BT}_{0.5}f^{\mathrm{C}}_{0.35}$ | | 0.5 | 0.35 | 0.5 | 0.5 |
| $\mathrm{BT}_{0.5}f^{\mathrm{C}}_{0.25}$ | | 0.5 | 0.25 | 0.5 | 0.5 |
| $\mathrm{BT}_{0.5}f^{\mathrm{C}}_{0.15}$ | | 0.5 | 0.15 | 0.5 | 0.5 |
| $\mathrm{BT}_{0.5}f^{\mathrm{C}}_{0.05}$ | | 0.5 | 0.05 | 0.5 | 0.5 |
| $\mathrm{BT}_{0.5}f^{\mathrm{C}}_{0.005}$ | | 0.5 | 0.005 | 0.5 | 0.5 |
| $\mathrm{BT}_{0.5}f^{\mathrm{F}}$ rules | | | | | |
| $\mathrm{BT}_{0.5}f^{\mathrm{F}}_{0.45}$ | | 0.5 | 0.5 | 0.45 | 0.5 |
| $\mathrm{BT}_{0.5}f^{\mathrm{F}}_{0.35}$ | | 0.5 | 0.5 | 0.35 | 0.5 |
| $\mathrm{BT}_{0.5}f^{\mathrm{F}}_{0.25}$ | | 0.5 | 0.5 | 0.25 | 0.5 |
| $\mathrm{BT}_{0.5}f^{\mathrm{F}}_{0.15}$ | | 0.5 | 0.5 | 0.15 | 0.5 |
| $\mathrm{BT}_{0.5}f^{\mathrm{F}}_{0.05}$ | | 0.5 | 0.5 | 0.05 | 0.5 |
| $\mathrm{BT}_{0.5}f^{\mathrm{F}}_{0.005}$ | | 0.5 | 0.5 | 0.005 | 0.5 |
| $\mathrm{BT}_{0.5}f^{\mathrm{B}}$ rules | | | | | |
| $\mathrm{BT}_{0.5}f^{\mathrm{B}}_{0.45}$ | | 0.5 | 0.5 | 0.5 | 0.45 |
| $\mathrm{BT}_{0.5}f^{\mathrm{B}}_{0.35}$ | | 0.5 | 0.5 | 0.5 | 0.35 |
| $\mathrm{BT}_{0.5}f^{\mathrm{B}}_{0.25}$ | | 0.5 | 0.5 | 0.5 | 0.25 |
| $\mathrm{BT}_{0.5}f^{\mathrm{B}}_{0.15}$ | | 0.5 | 0.5 | 0.5 | 0.15 |
| $\mathrm{BT}_{0.5}f^{\mathrm{B}}_{0.05}$ | | 0.5 | 0.5 | 0.5 | 0.05 |
| $\mathrm{BT}_{0.5}f^{\mathrm{B}}_{0.05}$ | | 0.5 | 0.5 | 0.5 | 0.005 |
| $\mathrm{BT}_{0.5}f^{\mathrm{FB}}$ rules | | | | | |
| $\mathrm{BT}_{0.5}f^{\mathrm{FB}}_{0.45}$ | | 0.5 | 0.5 | 0.45 | 0.45 |
| $\mathrm{BT}_{0.5}f^{\mathrm{FB}}_{0.35}$ | | 0.5 | 0.5 | 0.35 | 0.35 |
| $\mathrm{BT}_{0.5}f^{\mathrm{FB}}_{0.25}$ | | 0.5 | 0.5 | 0.25 | 0.25 |
| $\mathrm{BT}_{0.5}f^{\mathrm{FB}}_{0.15}$ | | 0.5 | 0.5 | 0.15 | 0.15 |
| $\mathrm{BT}_{0.5}f^{\mathrm{FB}}_{0.05}$ | | 0.5 | 0.5 | 0.05 | 0.05 |
| $\mathrm{BT}_{0.5}f^{\mathrm{FB}}_{0.005}$ | | 0.5 | 0.5 | 0.005 | 0.005 |

*Continue on the next page*

Table A4: All 69 Harvest Control rules (HCRs) used in this study (cont.).

| **Code** | **True** $F/F_{\mathbf{MSY}}$ | **Biomass threshold** | **Risk fractile for** $C_{y+1}$ | **Risk fractile for** $F_y/F_{\mathbf{MSY}}$ | **Risk fractile for** $B_{y+1}/B_{\mathbf{trigger}}$ |
|---|---|---|---|---|---|
| $\mathrm{BT}_{0.5}f^{\mathrm{CFB}}$ rules | | | | | |
| $\mathrm{BT}_{0.5}f^{\mathrm{CFB}}_{0.45}$ | | 0.5 | 0.45 | 0.45 | 0.45 |
| $\mathrm{BT}_{0.5}f^{\mathrm{CFB}}_{0.35}$ | | 0.5 | 0.35 | 0.35 | 0.35 |
| $\mathrm{BT}_{0.5}f^{\mathrm{CFB}}_{0.25}$ | | 0.5 | 0.25 | 0.25 | 0.25 |
| $\mathrm{BT}_{0.5}f^{\mathrm{CFB}}_{0.15}$ | | 0.5 | 0.15 | 0.15 | 0.15 |
| $\mathrm{BT}_{0.5}f^{\mathrm{CFB}}_{0.05}$ | | 0.5 | 0.05 | 0.05 | 0.05 |
| $\mathrm{BT}_{0.5}f^{\mathrm{CFB}}_{0.005}$ | | 0.5 | 0.005 | 0.005 | 0.005 |

#### Scenarios

Table A5: The definition of the 14 scenarios.

| Code | Observation noise (SD) | Time series length | Euler time steps | Intermediate year | Process noise | Implementation noise | Priors | Number of replicates |
|---|---|---|---|---|---|---|---|---|
| Baseline scenario | | | | | | | | |
| S1 | 0.3 | 40 | 4 | no | spec dep | no | all | 500 |
| Observation noise scenarios | | | | | | | | |
| S2 | 0.1 | 40 | 4 | no | spec dep | no | all | 2000 |
| S3 = Baseline | | | | | | | | 2000 |
| S4 | 0.5 | 40 | 4 | no | spec dep | no | all | 2000 |
| S5 | 0.7 | 40 | 4 | no | spec dep | no | all | 2000 |
| Sensitivity scenarios | | | | | | | | |
| S6 | 0.3 | 40 | 4 | no | spec dep | no | noise ratios | 500 |
| S7 | 0.3 | 40 | 4 | no | spec dep | no | no | 500 |
| S8 | 0.3 | 40 | 8 | no | spec dep | no | all | 500 |
| S9 | 0.3 | 40 | 4 | const F | spec dep | no | all | 500 |
| S10 | 0.3 | 40 | 4 | const C | spec dep | no | all | 500 |
| S11 | 0.3 | 20 | 4 | no | spec dep | no | all | 500 |
| S12 | 0.3 | 40 | 4 | no | $\sigma_R = 0.15, \rho_R = 0$ | no | all | 500 |
| S13 | 0.3 | 40 | 4 | no | spec dep | $\sigma_{Imp} = 0.15$ | all | 500 |
| S14[1] | 0.3 | 40 | 4 | no | spec dep | no | all | 500 |

[1] Scenario S14 exists only for anchovy and assumes a higher steepness parameter (h) for the stock-recruitment relationship of 0.9 instead of 0.75.

84 ## Supplementary Section B: Complementary results

85 ### Biomass Thresholds vs. $P^*$ method

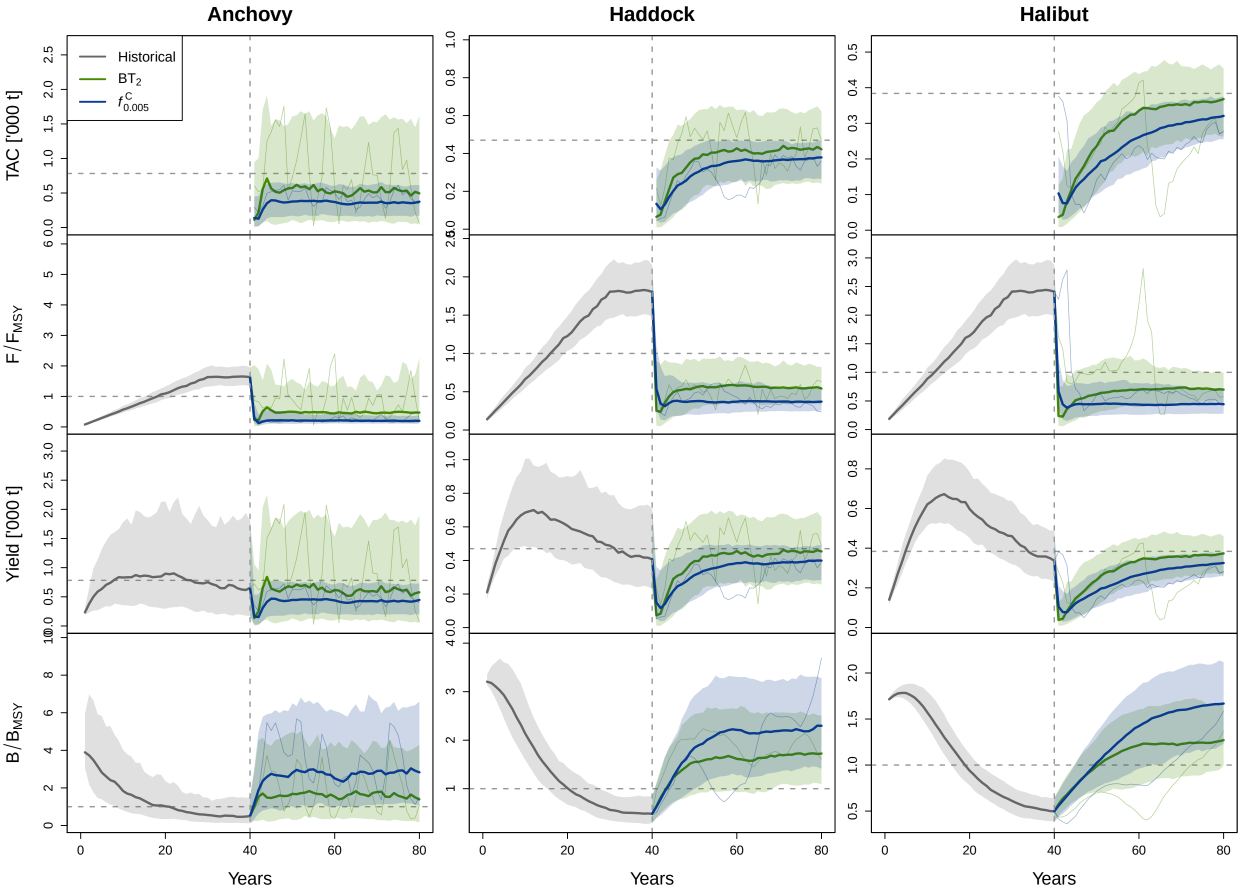


Figure B1: Trajectories of TAC, $F/F_{\text{MSY}}$, yield, and $B/B_{\text{MSY}}$ over historical and projection period. The thick coloured lines represent the median trajectories for the two HCRs, while the thin coloured lines represent the trajectory of a single replicate. The shaded area represents the 10th and 90th percentiles of the specific distributions. The vertical dashed line separates the historical (left of it) from the projection period (right to it). The horizontal dashed lines represent the reference points, i.e. 1 for the relative fishing mortality rate and biomass and MSY for TAC and yield.

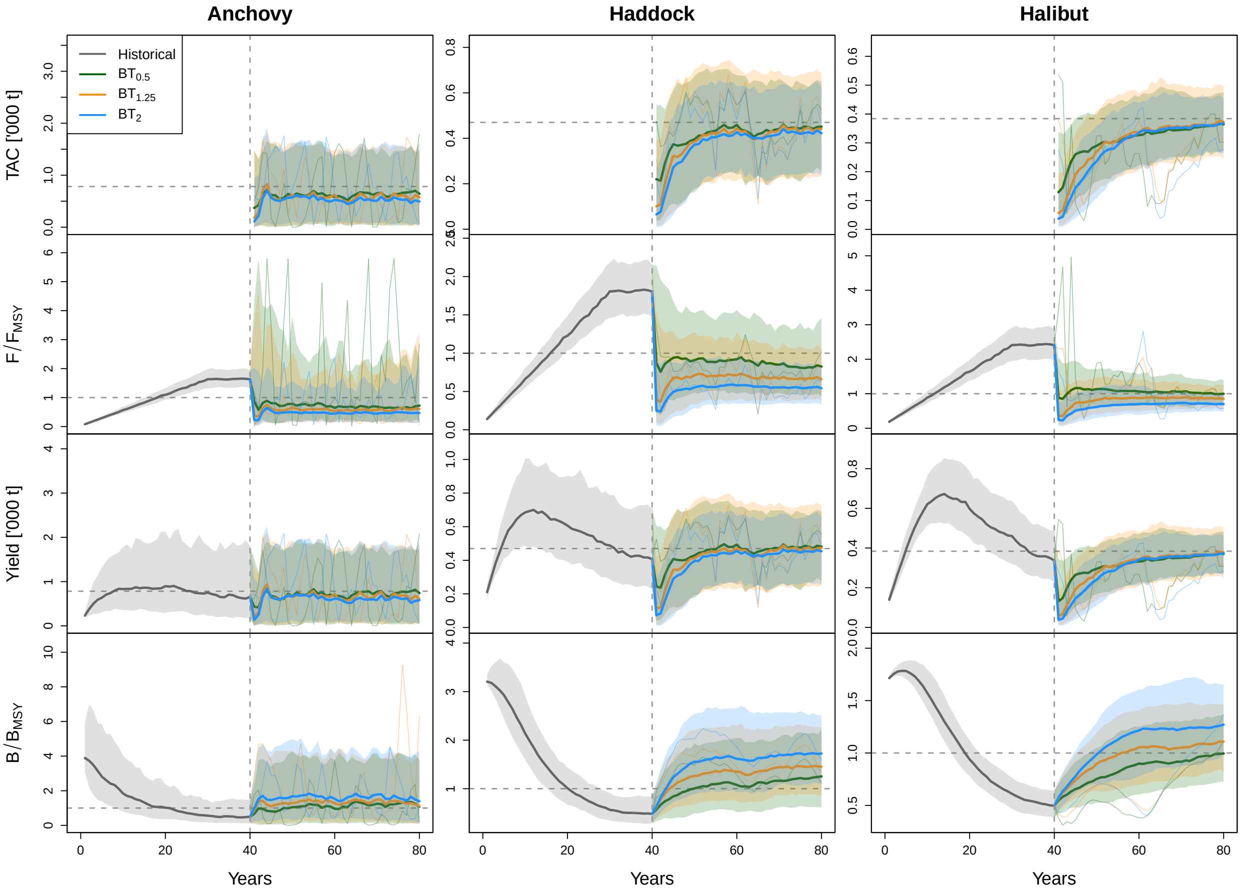


Figure B2: Trajectories of TAC, $F/F_{\mathrm{MSY}}$, yield, and $B/B_{\mathrm{MSY}}$ over historical and projection period. The thick coloured lines represent the median trajectories for the two HCRs, while the thin coloured lines represent the trajectory of a single replicate. The shaded area represents the 10th and 90th percentiles of the specific distributions. The vertical dashed line separates the historical (left of it) from the projection period (right to it). The horizontal dashed lines represent the reference points, i.e. 1 for the relative fishing mortality rate and biomass and MSY for TAC and yield.

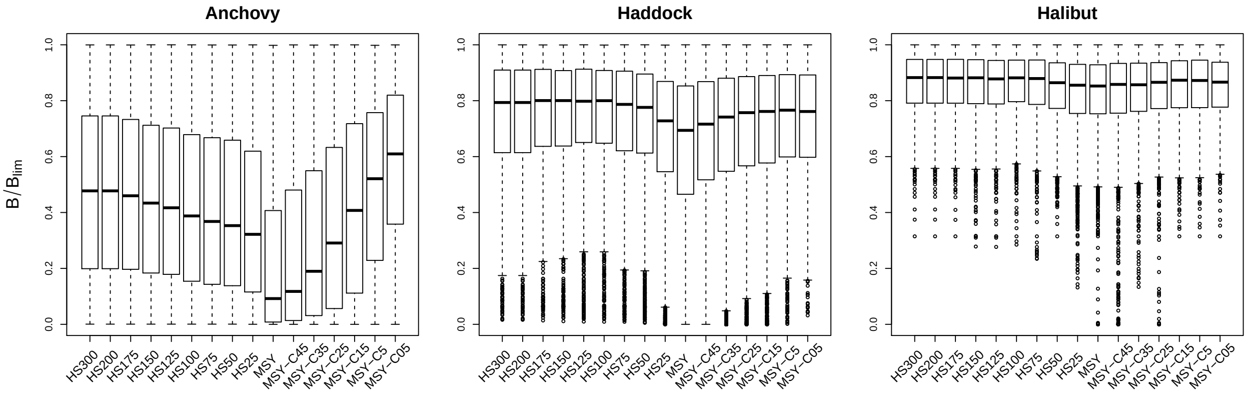


Figure B3: Relative biomass ($B/B_{\mathrm{lim}}$) of all replicates with biomass below $B_{\mathrm{lim}}$ for the threshold and $f^C$ rules.

Table B1: Proportion of converged replicates for each HCR and stock in percent.

| HCR | Anchovy | Haddock | Halibut |
|---|---|---|---|
| $f_{0.5}$ | 81 | 85 | 85 |
| $BT_{0.25}$ | 67 | 82 | 85 |
| $BT_{0.50}$ | 66 | 79 | 82 |
| $BT_{0.75}$ | 65 | 80 | 82 |
| $BT_{1.00}$ | 64 | 79 | 81 |
| $BT_{1.25}$ | 64 | 78 | 81 |
| $BT_{1.50}$ | 64 | 79 | 81 |
| $BT_{1.75}$ | 67 | 77 | 80 |
| $BT_{2.00}$ | 70 | 77 | 79 |
| $f^{C}_{0.45}$ | 82 | 84 | 86 |
| $f^{C}_{0.35}$ | 83 | 82 | 86 |
| $f^{C}_{0.25}$ | 84 | 82 | 86 |
| $f^{C}_{0.15}$ | 86 | 83 | 86 |
| $f^{C}_{0.05}$ | 85 | 82 | 84 |
| $f^{C}_{0.005}$ | 86 | 81 | 83 |
| $f^{F}_{0.45}$ | 82 | 85 | 84 |
| $f^{F}_{0.35}$ | 84 | 83 | 85 |
| $f^{F}_{0.25}$ | 85 | 82 | 82 |
| $f^{F}_{0.15}$ | 86 | 81 | 82 |
| $f^{F}_{0.05}$ | 86 | 80 | 81 |
| $f^{F}_{0.005}$ | 86 | 78 | 80 |
| $f^{CF}_{0.45}$ | 82 | 84 | 85 |
| $f^{CF}_{0.35}$ | 85 | 83 | 82 |
| $f^{CF}_{0.25}$ | 86 | 80 | 82 |
| $f^{CF}_{0.15}$ | 87 | 81 | 82 |
| $f^{CF}_{0.05}$ | 86 | 78 | 80 |
| $f^{CF}_{0.005}$ | 86 | 76 | 77 |
| $BT_{50}f^{C}_{0.45}$ | 67 | 81 | 83 |
| $BT_{50}f^{C}_{0.35}$ | 73 | 80 | 82 |
| $BT_{50}f^{C}_{0.25}$ | 77 | 81 | 83 |
| $BT_{50}f^{C}_{0.15}$ | 80 | 81 | 83 |
| $BT_{50}f^{C}_{0.05}$ | 84 | 79 | 83 |
| $BT_{50}$-$f^{C}_{0.005}$ | 85 | 78 | 81 |
| $BT_{50}f^{F}_{0.45}$ | 68 | 82 | 83 |
| $BT_{50}f^{F}_{0.35}$ | 74 | 81 | 81 |
| $BT_{50}f^{F}_{0.25}$ | 78 | 79 | 82 |
| $BT_{50}f^{F}_{0.15}$ | 82 | 78 | 81 |
| $BT_{50}f^{F}_{0.05}$ | 84 | 77 | 80 |
| $BT_{50}f^{F}_{0.005}$ | 86 | 77 | 77 |
| $BT_{50}f^{B}_{0.45}$ | 62 | 80 | 83 |
| $BT_{50}f^{B}_{0.35}$ | 59 | 80 | 81 |
| $BT_{50}f^{B}_{0.25}$ | 55 | 80 | 79 |
| $BT_{50}f^{B}_{0.15}$ | 51 | 79 | 79 |
| $BT_{50}f^{B}_{0.05}$ | 53 | 77 | 79 |
| $BT_{50}f^{B}_{0.005}$ | 64 | 75 | 78 |
| $BT_{50}f^{FB}_{0.45}$ | 65 | 81 | 81 |
| $BT_{50}f^{FB}_{0.35}$ | 73 | 81 | 81 |
| $BT_{50}f^{FB}_{0.25}$ | 76 | 79 | 82 |
| $BT_{50}f^{FB}_{0.15}$ | 83 | 76 | 79 |
| $BT_{50}f^{FB}_{0.05}$ | 84 | 74 | 77 |
| $BT_{50}f^{FB}_{0.005}$ | 86 | 74 | 75 |
| $BT_{50}f^{CFB}_{0.45}$ | 69 | 83 | 81 |
| $BT_{50}f^{CFB}_{0.35}$ | 78 | 80 | 81 |
| $BT_{50}f^{CFB}_{0.25}$ | 84 | 77 | 80 |
| $BT_{50}f^{CFB}_{0.15}$ | 85 | 75 | 78 |
| $BT_{50}f^{CFB}_{0.05}$ | 86 | 74 | 76 |
| $BT_{50}f^{CFB}_{0.005}$ | 86 | 75 | 73 |

Table B2: Risk, yield, and AAV of all threshold rules.

| Species | HCR | Yield | Risk | AAV |
|---|---|---|---|---|
| Anchovy | $f_{0.5}$ | 0.567 | 0.529 | 0.465 |
| Anchovy | $BT_{0.25}$ | 0.752 | 0.414 | 0.487 |
| Anchovy | $BT_{0.50}$ | 0.773 | 0.386 | 0.519 |
| Anchovy | $BT_{0.75}$ | 0.769 | 0.367 | 0.551 |
| Anchovy | $BT_{1.00}$ | 0.762 | 0.346 | 0.572 |
| Anchovy | $BT_{1.25}$ | 0.758 | 0.312 | 0.588 |
| Anchovy | $BT_{1.50}$ | 0.747 | 0.28 | 0.596 |
| Anchovy | $BT_{1.75}$ | 0.736 | 0.254 | 0.596 |
| Anchovy | $BT_{2.00}$ | 0.729 | 0.235 | 0.591 |
| Haddock | $f_{0.5}$ | 0.937 | 0.401 | 0.162 |
| Haddock | $BT_{0.25}$ | 0.945 | 0.381 | 0.168 |
| Haddock | $BT_{0.50}$ | 0.956 | 0.339 | 0.188 |
| Haddock | $BT_{0.75}$ | 0.952 | 0.286 | 0.218 |
| Haddock | $BT_{1.00}$ | 0.93 | 0.224 | 0.239 |
| Haddock | $BT_{1.25}$ | 0.911 | 0.176 | 0.247 |
| Haddock | $BT_{1.50}$ | 0.896 | 0.145 | 0.24 |
| Haddock | $BT_{1.75}$ | 0.883 | 0.119 | 0.225 |
| Haddock | $BT_{2.00}$ | 0.867 | 0.101 | 0.213 |
| Halibut | $f_{0.5}$ | 0.969 | 0.31 | 0.109 |
| Halibut | $BT_{0.25}$ | 0.972 | 0.31 | 0.112 |
| Halibut | $BT_{0.50}$ | 0.977 | 0.275 | 0.122 |
| Halibut | $BT_{0.75}$ | 0.987 | 0.236 | 0.14 |
| Halibut | $BT_{1.00}$ | 0.974 | 0.192 | 0.161 |
| Halibut | $BT_{1.25}$ | 0.954 | 0.161 | 0.16 |
| Halibut | $BT_{1.50}$ | 0.94 | 0.144 | 0.148 |
| Halibut | $BT_{1.75}$ | 0.924 | 0.126 | 0.138 |
| Halibut | $BT_{2.00}$ | 0.909 | 0.118 | 0.128 |

Table B3: Risk of all HS rules for the scenarios with different scientific uncertainty evaluated over the whole time period.

| Species | HCR | Risk (SD=0.1) | Risk (SD=0.3) | Risk (SD=0.5) | Risk (SD=0.7) |
|---|---|---|---|---|---|
| Anchovy | $f_{0.5}$ | 0.261 | 0.322 | 0.368 | 0.408 |
| Anchovy | $f^{C}_{0.45}$ | 0.232 | 0.28 | 0.317 | 0.353 |
| Anchovy | $f^{C}_{0.15}$ | 0.095 | 0.104 | 0.113 | 0.118 |
| Anchovy | $BT_{0.50}$ | 0.166 | 0.229 | 0.286 | 0.341 |
| Anchovy | $BT_{50}f^{C}_{0.25}$ | 0.099 | 0.122 | 0.142 | 0.157 |
| Haddock | $f_{0.5}$ | 0.319 | 0.328 | 0.347 | 0.401 |
| Haddock | $f^{C}_{0.45}$ | 0.296 | 0.301 | 0.314 | 0.362 |
| Haddock | $f^{C}_{0.15}$ | 0.162 | 0.151 | 0.144 | 0.147 |
| Haddock | $BT_{0.50}$ | 0.279 | 0.287 | 0.302 | 0.348 |
| Haddock | $BT_{50}f^{C}_{0.25}$ | 0.18 | 0.172 | 0.172 | 0.187 |
| Halibut | $f_{0.5}$ | 0.245 | 0.251 | 0.292 | 0.362 |
| Halibut | $f^{C}_{0.45}$ | 0.23 | 0.231 | 0.266 | 0.326 |
| Halibut | $f^{C}_{0.15}$ | 0.159 | 0.138 | 0.133 | 0.141 |
| Halibut | $BT_{0.50}$ | 0.217 | 0.223 | 0.257 | 0.32 |
| Halibut | $BT_{50}f^{C}_{0.25}$ | 0.16 | 0.146 | 0.153 | 0.175 |

Table B4: Yield of all HS rules for the scenarios with different scientific uncertainty evaluated over the whole time period.

| Species | HCR | Rel. yield (SD=0.1) | Rel. yield (SD=0.3) | Rel. yield (SD=0.5) | Rel. yield (SD=0.7) |
|---|---|---|---|---|---|
| Anchovy | $f_{0.5}$ | 0.823 | 0.766 | 0.684 | 0.588 |
| Anchovy | $f^{C}_{0.45}$ | 0.823 | 0.778 | 0.708 | 0.633 |
| Anchovy | $f^{C}_{0.15}$ | 0.75 | 0.715 | 0.676 | 0.639 |
| Anchovy | $BT_{0.50}$ | 0.815 | 0.813 | 0.758 | 0.694 |
| Anchovy | $BT_{50}f^{C}_{0.25}$ | 0.763 | 0.753 | 0.728 | 0.696 |
| Haddock | $f_{0.5}$ | 0.965 | 0.947 | 0.92 | 0.888 |
| Haddock | $f^{C}_{0.45}$ | 0.961 | 0.943 | 0.918 | 0.894 |
| Haddock | $f^{C}_{0.15}$ | 0.925 | 0.897 | 0.867 | 0.849 |
| Haddock | $BT_{0.50}$ | 0.968 | 0.956 | 0.94 | 0.927 |
| Haddock | $BT_{50}f^{C}_{0.25}$ | 0.939 | 0.92 | 0.903 | 0.892 |
| Halibut | $f_{0.5}$ | 0.983 | 0.971 | 0.96 | 0.944 |
| Halibut | $f^{C}_{0.45}$ | 0.981 | 0.968 | 0.957 | 0.945 |
| Halibut | $f^{C}_{0.15}$ | 0.953 | 0.92 | 0.9 | 0.889 |
| Halibut | $BT_{0.50}$ | 0.987 | 0.978 | 0.974 | 0.971 |
| Halibut | $BT_{50}f^{C}_{0.25}$ | 0.965 | 0.943 | 0.934 | 0.933 |

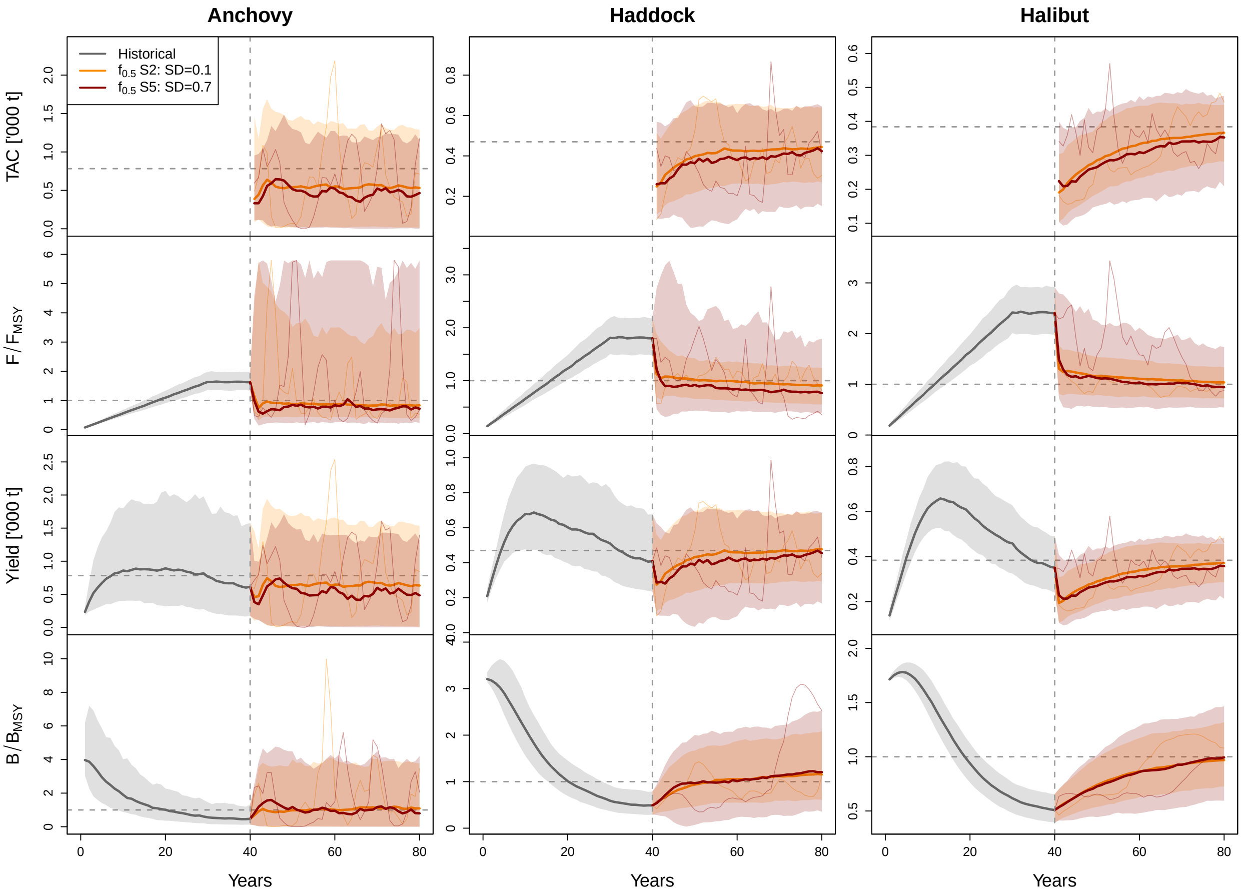


Figure B4: Trajectories of TAC, $F/F_{\text{MSY}}$, yield, and $B/B_{\text{MSY}}$ over historical and projection period for the MSY rule. The thick coloured lines represent the median trajectories for the two HCRs, while the thin coloured lines represent the trajectory of a single replicate. The shaded area represents the 10th and 90th percentiles of the specific distributions. The vertical dashed line separates the historical (left of it) from the projection period (right to it). The horizontal dashed lines represent the reference points, i.e. 1 for the relative fishing mortality rate and biomass and MSY for TAC and yield.

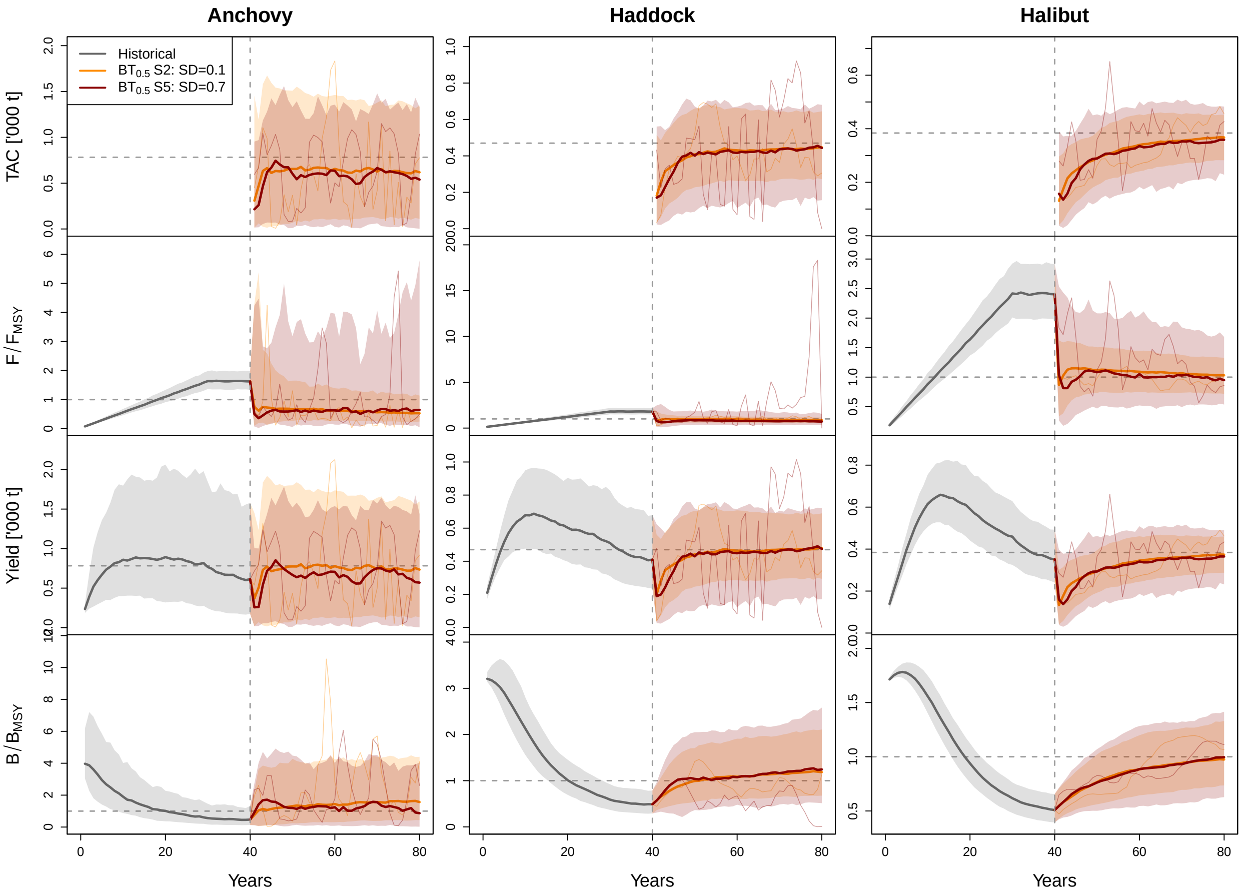


Figure B5: Trajectories of TAC, $F/F_{\text{MSY}}$, yield, and $B/B_{\text{MSY}}$ over historical and projection period for the $\text{BT}_{0.5}$ rule. The thick coloured lines represent the median trajectories for the two HCRs, while the thin coloured lines represent the trajectory of a single replicate. The shaded area represents the 10th and 90th percentiles of the specific distributions. The vertical dashed line separates the historical (left of it) from the projection period (right to it). The horizontal dashed lines represent the reference points, i.e. 1 for the relative fishing mortality rate and biomass and MSY for TAC and yield.

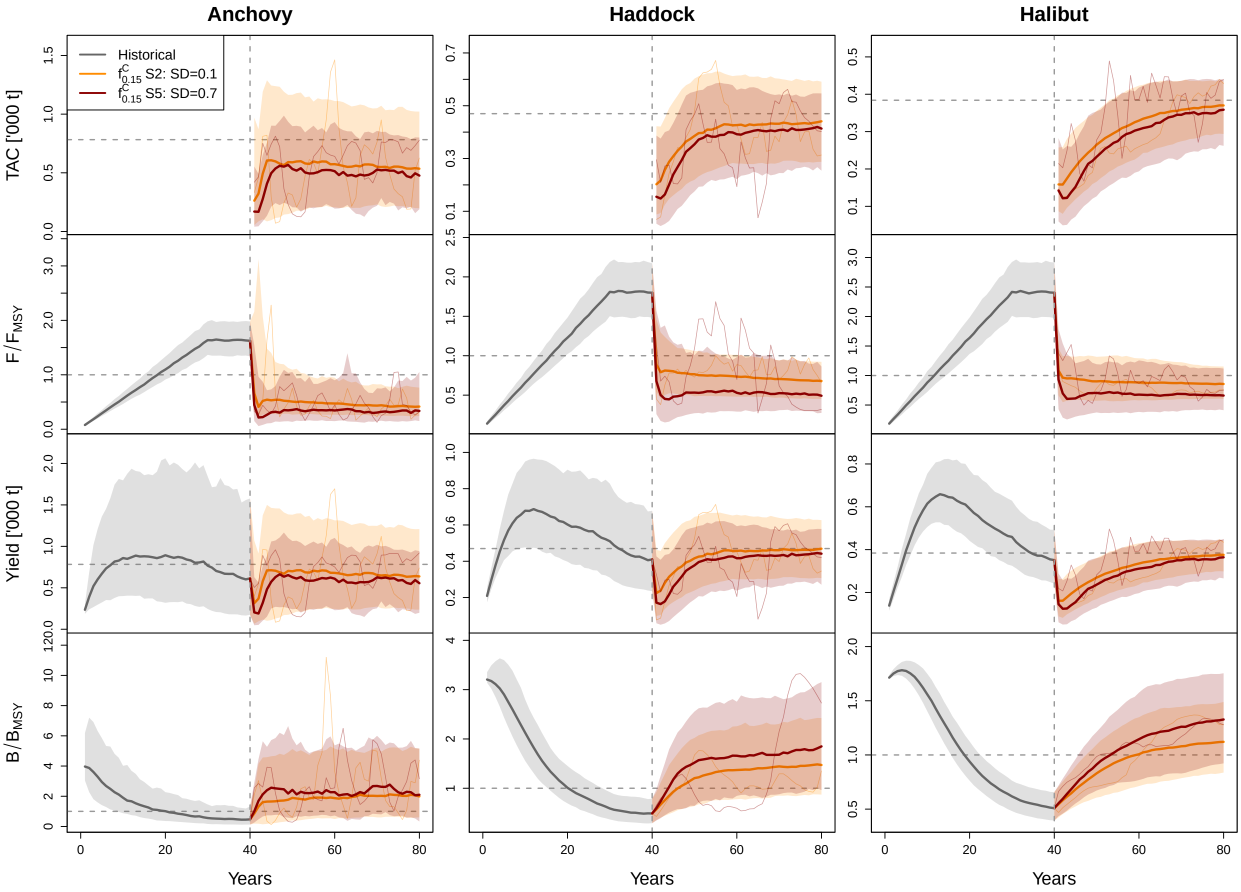


Figure B6: Trajectories of TAC, $F/F_{\mathrm{MSY}}$, yield, and $B/B_{\mathrm{MSY}}$ over historical and projection period for the $f^C_{15}$. The thick coloured lines represent the median trajectories for the two HCRs, while the thin coloured lines represent the trajectory of a single replicate. The shaded area represents the 10th and 90th percentiles of the specific distributions. The vertical dashed line separates the historical (left of it) from the projection period (right to it). The horizontal dashed lines represent the reference points, i.e. 1 for the relative fishing mortality rate and biomass and MSY for TAC and yield.

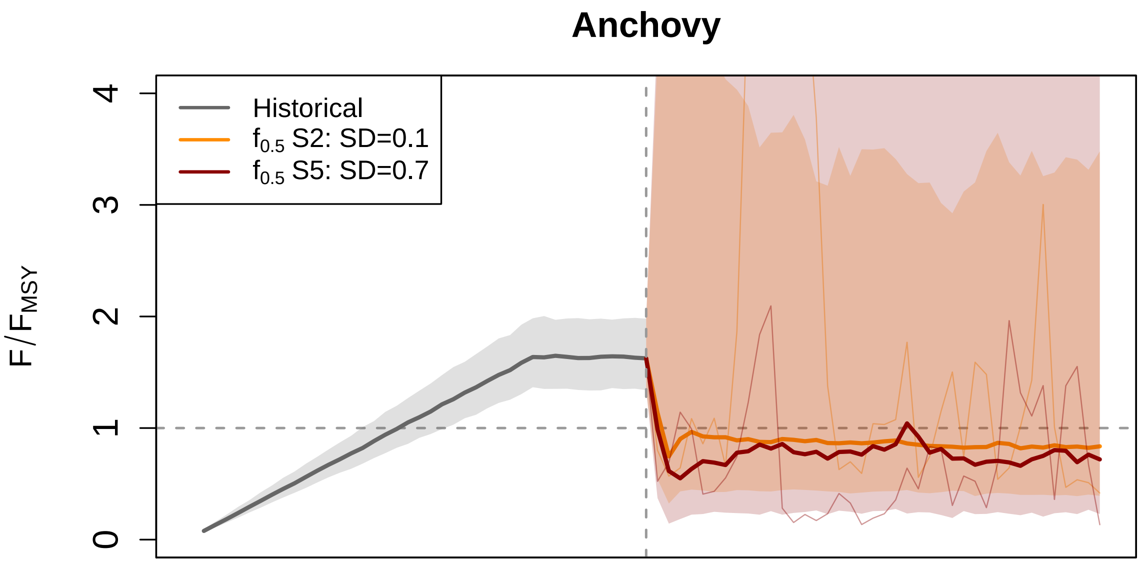


Figure B7: Trajectories of TAC, $F/F_{\mathrm{MSY}}$ over historical and projection period for the MSY rule (adjusted range of y axis). The thick coloured lines represent the median trajectories for the two HCRs, while the thin coloured lines represent the trajectory of a single replicate. The shaded area represents the 10th and 90th percentiles of the specific distributions. The vertical dashed line separates the historical (left of it) from the projection period (right to it). The horizontal dashed lines represent the reference point.

Table B5: Annual absolute variability in yield (AAV) of all HS rules for the scenarios with different scientific uncertainty evaluated over the whole time period.

| Species | HCR | AAV (SD=0.1) | AAV (SD=0.3) | AAV (SD=0.5) | AAV (SD=0.7) |
|---|---|---|---|---|---|
| Anchovy | $f_{0.5}$ | 0.33 | 0.369 | 0.383 | 0.397 |
| Anchovy | $f^{C}_{0.45}$ | 0.318 | 0.351 | 0.359 | 0.369 |
| Anchovy | $f^{C}_{0.15}$ | 0.259 | 0.273 | 0.265 | 0.26 |
| Anchovy | $\mathrm{BT}_{0.50}$ | 0.377 | 0.439 | 0.462 | 0.482 |
| Anchovy | $\mathrm{BT}_{50}f^{C}_{0.25}$ | 0.289 | 0.318 | 0.32 | 0.32 |
| Haddock | $f_{0.5}$ | 0.114 | 0.146 | 0.172 | 0.199 |
| Haddock | $f^{C}_{0.45}$ | 0.112 | 0.142 | 0.166 | 0.19 |
| Haddock | $f^{C}_{0.15}$ | 0.098 | 0.114 | 0.125 | 0.135 |
| Haddock | $\mathrm{BT}_{0.50}$ | 0.128 | 0.171 | 0.206 | 0.243 |
| Haddock | $\mathrm{BT}_{50}f^{C}_{0.25}$ | 0.108 | 0.131 | 0.15 | 0.167 |
| Halibut | $f_{0.5}$ | 0.068 | 0.098 | 0.126 | 0.153 |
| Halibut | $f^{C}_{0.45}$ | 0.067 | 0.095 | 0.121 | 0.147 |
| Halibut | $f^{C}_{0.15}$ | 0.058 | 0.074 | 0.091 | 0.106 |
| Halibut | $\mathrm{BT}_{0.50}$ | 0.075 | 0.109 | 0.145 | 0.184 |
| Halibut | $\mathrm{BT}_{50}f^{C}_{0.25}$ | 0.064 | 0.085 | 0.108 | 0.129 |

### All Harvest Control Rules (HCRs)

Table B6: Predicted yield and annual absolute variability in yield (AAV) for selected risk levels as well as the biomass threshold or risk fractile required to reach those risk levels for different HCR types and all species. All metrics are estimated over the whole management period.

| Species | HCR | Risk | Yield | AAV | Threshold/risk fractile |
|---|---|---|---|---|---|
| Anchovy | BT | 0.25 | 0.73 | 0.60 | 2.07 |
| Anchovy | $f^{\mathrm{C}}$ | 0.25 | 0.73 | 0.33 | 0.19 |
| Anchovy | $f^{\mathrm{F}}$ | 0.25 | 0.72 | 0.33 | 0.16 |
| Anchovy | $f^{\mathrm{FB}}$ | 0.25 | 0.75 | 0.33 | 0.32 |
| Anchovy | $\mathrm{BT}_{0.5}f^{\mathrm{C}}$ | 0.25 | 0.77 | 0.39 | 0.28 |
| Anchovy | $\mathrm{BT}_{0.5}f^{\mathrm{F}}$ | 0.25 | 0.77 | 0.39 | 0.28 |
| Anchovy | $\mathrm{BT}_{0.5}f^{\mathrm{B}}$ | 0.25 | 0.72 | 0.64 | 0.03 |
| Anchovy | $\mathrm{BT}_{0.5}f^{\mathrm{FB}}$ | 0.25 | 0.78 | 0.44 | 0.31 |
| Anchovy | $\mathrm{BT}_{0.5}f^{\mathrm{CFB}}$ | 0.25 | 0.79 | 0.41 | 0.40 |
| Haddock | BT | 0.15 | 0.90 | 0.24 | 1.69 |
| Haddock | $f^{\mathrm{C}}$ | 0.15 | 0.89 | 0.11 | 0.09 |
| Haddock | $f^{\mathrm{F}}$ | 0.15 | 0.87 | 0.13 | 0.15 |
| Haddock | $f^{\mathrm{FB}}$ | 0.15 | 0.88 | 0.12 | 0.29 |
| Haddock | $\mathrm{BT}_{0.5}f^{\mathrm{C}}$ | 0.15 | 0.89 | 0.12 | 0.13 |
| Haddock | $\mathrm{BT}_{0.5}f^{\mathrm{F}}$ | 0.15 | 0.90 | 0.13 | 0.20 |
| Haddock | $\mathrm{BT}_{0.5}f^{\mathrm{B}}$ | 0.15 | 0.92 | 0.31 | 0.01 |
| Haddock | $\mathrm{BT}_{0.5}f^{\mathrm{FB}}$ | 0.15 | 0.91 | 0.15 | 0.24 |
| Haddock | $\mathrm{BT}_{0.5}f^{\mathrm{CFB}}$ | 0.15 | 0.89 | 0.13 | 0.33 |
| Halibut | BT | 0.15 | 0.95 | 0.15 | 1.64 |
| Halibut | $f^{\mathrm{C}}$ | 0.15 | 0.89 | 0.08 | 0.07 |
| Halibut | $f^{\mathrm{F}}$ | 0.15 | 0.91 | 0.10 | 0.19 |
| Halibut | $f^{\mathrm{FB}}$ | 0.15 | 0.91 | 0.09 | 0.29 |
| Halibut | $\mathrm{BT}_{0.5}f^{\mathrm{C}}$ | 0.15 | 0.91 | 0.08 | 0.10 |
| Halibut | $\mathrm{BT}_{0.5}f^{\mathrm{F}}$ | 0.15 | 0.93 | 0.10 | 0.24 |
| Halibut | $\mathrm{BT}_{0.5}f^{\mathrm{B}}$ | 0.15 | 0.97 | 0.20 | 0.02 |
| Halibut | $\mathrm{BT}_{0.5}f^{\mathrm{FB}}$ | 0.15 | 0.95 | 0.11 | 0.27 |
| Halibut | $\mathrm{BT}_{0.5}f^{\mathrm{CFB}}$ | 0.15 | 0.93 | 0.10 | 0.34 |

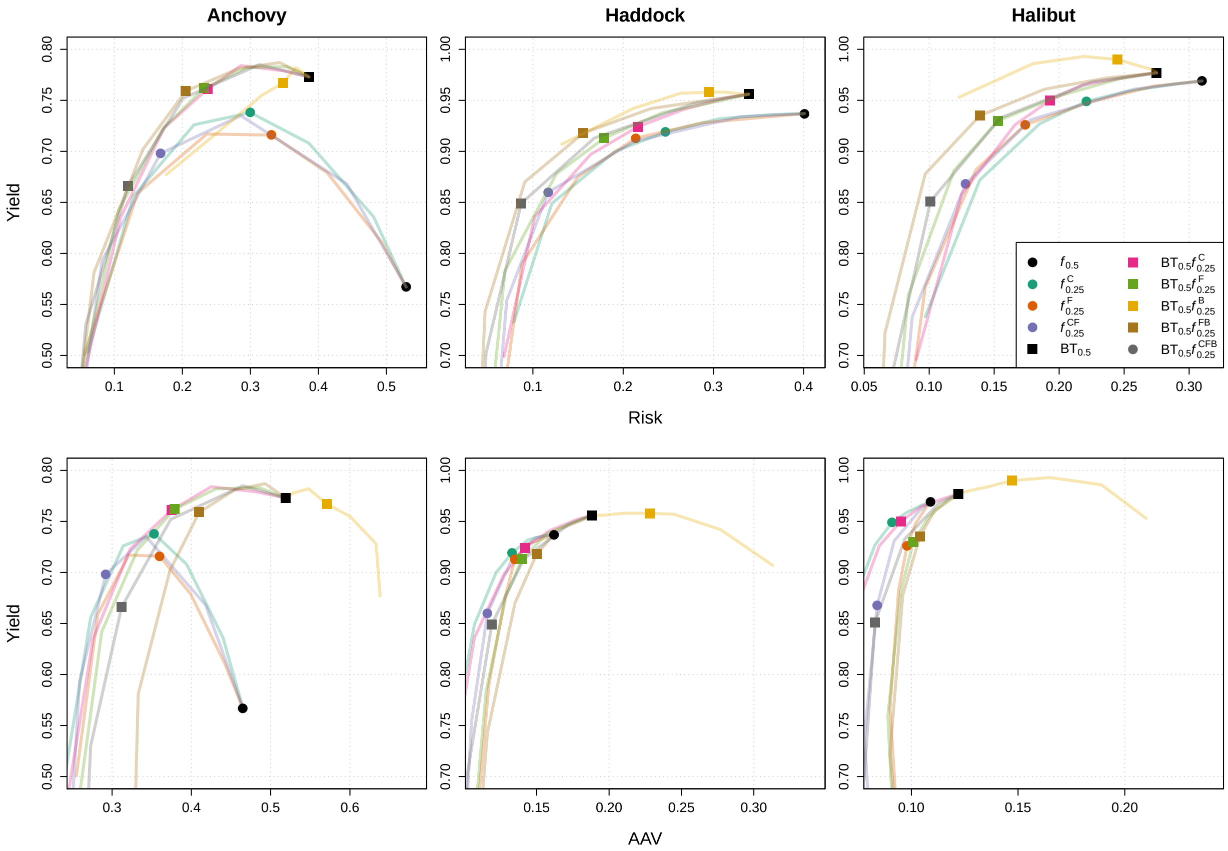


Figure B8: Zoomed into trade-off between risk and relative yield (upper row) and between inter-annal variability in yield (AAV) and yield (lower row) for all HCRs and anchovy, haddock, and halibut (columns) evaluated over the whole projection period (year 41-80). Starting at the black diamond (MSY rule), the lines connect HCRs without a biomass threshold and with increasing buffers. Similarly, starting at the black square ($BT_{0.5}$ rule), the lines connect HCRs with a biomass threshold and increasing buffers. Repeated symbols connected with lines represent the different quantities considered for the uncertainty buffer and include following risk fractiles: 0.45, 0.35, 0.25, 0.15, 0.05, 0.005.

Table B7: Risk of all fractile rules evaluated over the whole projection period (years 41-80).

| Species | Risk fractile | $f^{C}$ | $f^{F}$ | $f^{FB}$ | $BT_{0.5}f^{C}$ | $BT_{0.5}f^{F}$ | $BT_{0.5}f^{B}$ | $BT_{0.5}f^{FB}$ | $BT_{0.5}f^{CFB}$ |
|---|---|---|---|---|---|---|---|---|---|
| Anchovy | 0.5 | 0.529 | 0.529 | 0.529 | 0.386 | 0.386 | 0.386 | 0.386 | 0.386 |
| Anchovy | 0.45 | 0.481 | 0.493 | 0.442 | 0.349 | 0.353 | 0.376 | 0.343 | 0.313 |
| Anchovy | 0.35 | 0.386 | 0.413 | 0.286 | 0.286 | 0.294 | 0.367 | 0.282 | 0.201 |
| Anchovy | 0.25 | 0.3 | 0.331 | 0.168 | 0.237 | 0.232 | 0.348 | 0.205 | 0.12 |
| Anchovy | 0.15 | 0.217 | 0.236 | 0.082 | 0.173 | 0.171 | 0.317 | 0.142 | 0.058 |
| Anchovy | 0.05 | 0.128 | 0.134 | 0.029 | 0.109 | 0.106 | 0.269 | 0.07 | 0.019 |
| Anchovy | 0.005 | 0.061 | 0.056 | 0.012 | 0.055 | 0.048 | 0.176 | 0.023 | 0.01 |
| Haddock | 0.5 | 0.401 | 0.401 | 0.401 | 0.339 | 0.339 | 0.339 | 0.339 | 0.339 |
| Haddock | 0.45 | 0.365 | 0.363 | 0.33 | 0.313 | 0.31 | 0.334 | 0.301 | 0.279 |
| Haddock | 0.35 | 0.307 | 0.288 | 0.213 | 0.267 | 0.244 | 0.313 | 0.231 | 0.168 |
| Haddock | 0.25 | 0.247 | 0.214 | 0.117 | 0.216 | 0.179 | 0.295 | 0.156 | 0.087 |
| Haddock | 0.15 | 0.191 | 0.15 | 0.071 | 0.164 | 0.126 | 0.264 | 0.091 | 0.048 |
| Haddock | 0.05 | 0.121 | 0.088 | 0.047 | 0.102 | 0.069 | 0.211 | 0.047 | 0.032 |
| Haddock | 0.005 | 0.079 | 0.058 | 0.035 | 0.068 | 0.045 | 0.132 | 0.032 | 0.03 |
| Halibut | 0.5 | 0.31 | 0.31 | 0.31 | 0.275 | 0.275 | 0.275 | 0.275 | 0.275 |
| Halibut | 0.45 | 0.289 | 0.273 | 0.258 | 0.254 | 0.246 | 0.272 | 0.237 | 0.225 |
| Halibut | 0.35 | 0.253 | 0.223 | 0.178 | 0.218 | 0.199 | 0.261 | 0.189 | 0.155 |
| Halibut | 0.25 | 0.221 | 0.174 | 0.128 | 0.193 | 0.153 | 0.245 | 0.139 | 0.101 |
| Halibut | 0.15 | 0.185 | 0.136 | 0.087 | 0.166 | 0.119 | 0.219 | 0.097 | 0.07 |
| Halibut | 0.05 | 0.139 | 0.097 | 0.066 | 0.125 | 0.084 | 0.18 | 0.066 | 0.055 |
| Halibut | 0.005 | 0.097 | 0.073 | 0.057 | 0.09 | 0.065 | 0.123 | 0.055 | 0.05 |

Table B8: Yield of all fractile rules evaluated over the whole projection period (years 41-80).

| Species | Risk fractile | $f^{C}$ | $f^{F}$ | $f^{FB}$ | $BT_{0.5}f^{C}$ | $BT_{0.5}f^{F}$ | $BT_{0.5}f^{B}$ | $BT_{0.5}f^{FB}$ | $BT_{0.5}f^{CFB}$ |
|---|---|---|---|---|---|---|---|---|---|
| Anchovy | 0.5 | 0.567 | 0.567 | 0.567 | 0.773 | 0.773 | 0.773 | 0.773 | 0.773 |
| Anchovy | 0.45 | 0.636 | 0.611 | 0.668 | 0.779 | 0.783 | 0.778 | 0.787 | 0.785 |
| Anchovy | 0.35 | 0.708 | 0.679 | 0.735 | 0.784 | 0.782 | 0.782 | 0.781 | 0.752 |
| Anchovy | 0.25 | 0.738 | 0.716 | 0.698 | 0.761 | 0.762 | 0.767 | 0.759 | 0.666 |
| Anchovy | 0.15 | 0.726 | 0.717 | 0.593 | 0.723 | 0.722 | 0.755 | 0.702 | 0.53 |
| Anchovy | 0.05 | 0.656 | 0.659 | 0.345 | 0.635 | 0.642 | 0.728 | 0.581 | 0.247 |
| Anchovy | 0.005 | 0.503 | 0.501 | 0.074 | 0.48 | 0.47 | 0.677 | 0.32 | 0.025 |
| Haddock | 0.5 | 0.937 | 0.937 | 0.937 | 0.956 | 0.956 | 0.956 | 0.956 | 0.956 |
| Haddock | 0.45 | 0.936 | 0.934 | 0.934 | 0.952 | 0.95 | 0.956 | 0.951 | 0.947 |
| Haddock | 0.35 | 0.932 | 0.928 | 0.91 | 0.941 | 0.938 | 0.958 | 0.942 | 0.913 |
| Haddock | 0.25 | 0.919 | 0.913 | 0.86 | 0.924 | 0.913 | 0.958 | 0.918 | 0.849 |
| Haddock | 0.15 | 0.9 | 0.875 | 0.754 | 0.897 | 0.878 | 0.957 | 0.87 | 0.702 |
| Haddock | 0.05 | 0.849 | 0.79 | 0.501 | 0.836 | 0.783 | 0.942 | 0.743 | 0.366 |
| Haddock | 0.005 | 0.733 | 0.601 | 0.15 | 0.699 | 0.573 | 0.907 | 0.465 | 0.066 |
| Halibut | 0.5 | 0.969 | 0.969 | 0.969 | 0.977 | 0.977 | 0.977 | 0.977 | 0.977 |
| Halibut | 0.45 | 0.966 | 0.964 | 0.961 | 0.973 | 0.971 | 0.98 | 0.972 | 0.968 |
| Halibut | 0.35 | 0.959 | 0.948 | 0.931 | 0.965 | 0.954 | 0.984 | 0.961 | 0.933 |
| Halibut | 0.25 | 0.949 | 0.926 | 0.868 | 0.95 | 0.93 | 0.99 | 0.935 | 0.851 |
| Halibut | 0.15 | 0.927 | 0.882 | 0.738 | 0.926 | 0.881 | 0.993 | 0.878 | 0.673 |
| Halibut | 0.05 | 0.872 | 0.767 | 0.447 | 0.858 | 0.759 | 0.986 | 0.722 | 0.317 |
| Halibut | 0.005 | 0.738 | 0.538 | 0.122 | 0.696 | 0.512 | 0.953 | 0.407 | 0.058 |

Table B9: Annual absolute variability in yield (AAV) of all fractile rules evaluated over the whole projection period (years 42-80).

| Species | Risk fractile | $f^{C}$ | $f^{F}$ | $f^{FB}$ | $BT_{0.5}f^{C}$ | $BT_{0.5}f^{F}$ | $BT_{0.5}f^{B}$ | $BT_{0.5}f^{FB}$ | $BT_{0.5}f^{CFB}$ |
|---|---|---|---|---|---|---|---|---|---|
| Anchovy | 0.5 | 0.465 | 0.465 | 0.465 | 0.519 | 0.519 | 0.519 | 0.519 | 0.519 |
| Anchovy | 0.45 | 0.44 | 0.442 | 0.418 | 0.482 | 0.486 | 0.529 | 0.493 | 0.465 |
| Anchovy | 0.35 | 0.394 | 0.399 | 0.343 | 0.425 | 0.431 | 0.548 | 0.455 | 0.374 |
| Anchovy | 0.25 | 0.353 | 0.36 | 0.292 | 0.375 | 0.379 | 0.571 | 0.41 | 0.312 |
| Anchovy | 0.15 | 0.314 | 0.321 | 0.259 | 0.323 | 0.332 | 0.6 | 0.373 | 0.273 |
| Anchovy | 0.05 | 0.273 | 0.282 | 0.24 | 0.276 | 0.287 | 0.633 | 0.333 | 0.256 |
| Anchovy | 0.005 | 0.241 | 0.255 | 0.262 | 0.244 | 0.257 | 0.638 | 0.324 | 0.307 |
| Haddock | 0.5 | 0.162 | 0.162 | 0.162 | 0.188 | 0.188 | 0.188 | 0.188 | 0.188 |
| Haddock | 0.45 | 0.155 | 0.156 | 0.149 | 0.178 | 0.177 | 0.196 | 0.181 | 0.172 |
| Haddock | 0.35 | 0.144 | 0.144 | 0.132 | 0.159 | 0.157 | 0.21 | 0.165 | 0.141 |
| Haddock | 0.25 | 0.133 | 0.135 | 0.116 | 0.142 | 0.14 | 0.228 | 0.15 | 0.119 |
| Haddock | 0.15 | 0.122 | 0.128 | 0.105 | 0.127 | 0.129 | 0.245 | 0.135 | 0.102 |
| Haddock | 0.05 | 0.107 | 0.117 | 0.095 | 0.107 | 0.115 | 0.277 | 0.116 | 0.092 |
| Haddock | 0.005 | 0.092 | 0.105 | 0.109 | 0.091 | 0.102 | 0.313 | 0.1 | 0.106 |
| Halibut | 0.5 | 0.109 | 0.109 | 0.109 | 0.122 | 0.122 | 0.122 | 0.122 | 0.122 |
| Halibut | 0.45 | 0.105 | 0.107 | 0.104 | 0.116 | 0.117 | 0.127 | 0.119 | 0.113 |
| Halibut | 0.35 | 0.098 | 0.102 | 0.092 | 0.105 | 0.108 | 0.136 | 0.111 | 0.097 |
| Halibut | 0.25 | 0.091 | 0.098 | 0.084 | 0.095 | 0.101 | 0.147 | 0.104 | 0.083 |
| Halibut | 0.15 | 0.083 | 0.094 | 0.078 | 0.085 | 0.095 | 0.165 | 0.096 | 0.077 |
| Halibut | 0.05 | 0.073 | 0.09 | 0.087 | 0.073 | 0.089 | 0.189 | 0.09 | 0.09 |
| Halibut | 0.005 | 0.063 | 0.097 | 0.124 | 0.061 | 0.095 | 0.21 | 0.104 | 0.119 |

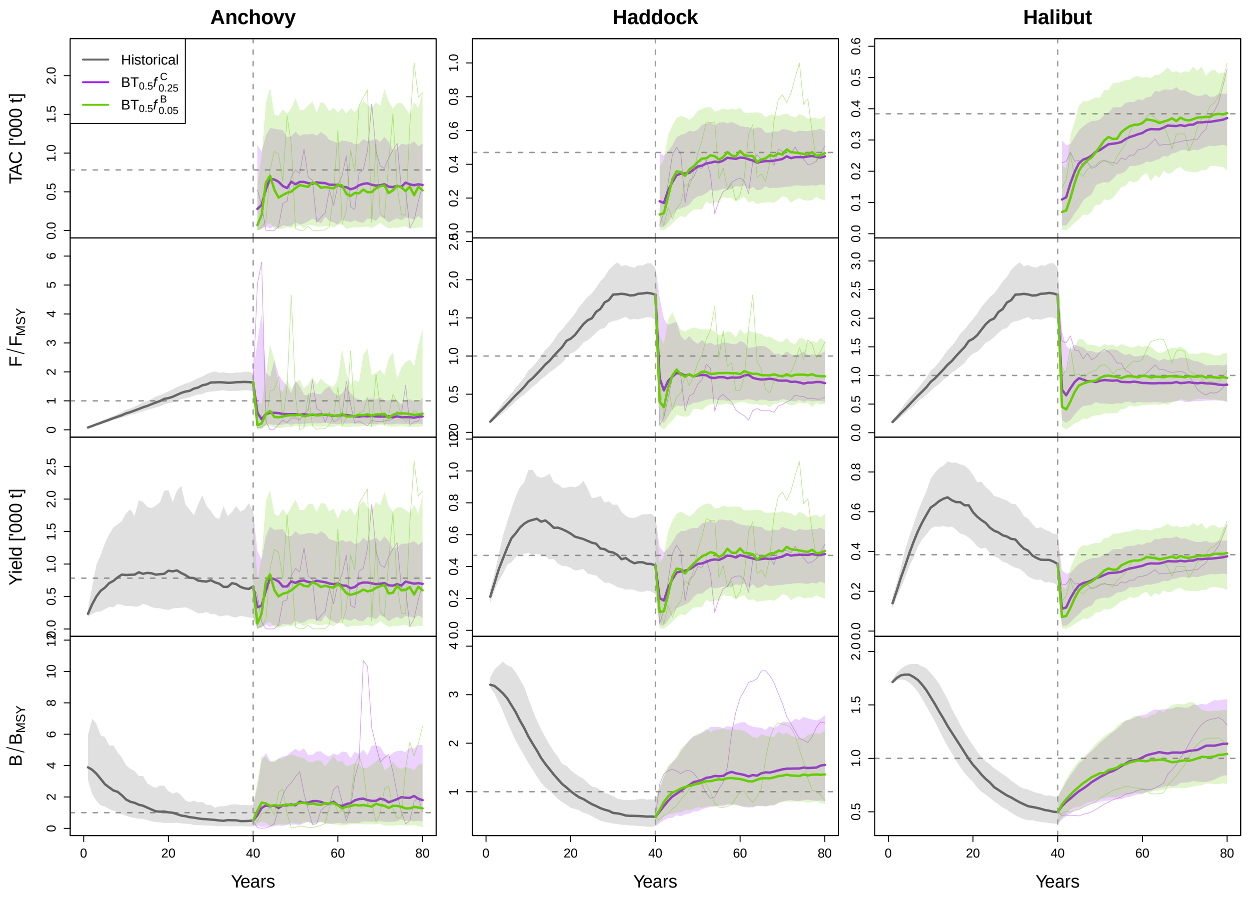


Figure B9: Trajectories of TAC, $F/F_{\mathrm{MSY}}$, yield, and $B/B_{\mathrm{MSY}}$ over historical and projection period. The thick coloured lines represent the median trajectories for the two HCRs, while the thin coloured lines represent the trajectory of a single replicate. The shaded area represents the 10th and 90th percentiles of the specific distributions. The vertical dashed line separates the historical (left of it) from the projection period (right to it). The horizontal dashed lines represent the reference points, i.e. 1 for the relative fishing mortality rate and biomass and MSY for TAC and yield.

Table B10: Median relative error (median($(x_{\text{est}} - x_{\text{true}})/x_{\text{true}}$)) of estimated $B/B_{\text{MSY}}$ and $F/F_{\text{MSY}}$ for scenario 2 (observation SD=0.1), scenario 3 (baseline, observation SD=0.3), and scenario 5 (observation SD=0.7) for selected HCR and all species.

| Species | HCR | $B/B_{\text{MSY}}$ S2 | $B/B_{\text{MSY}}$ S3 | $B/B_{\text{MSY}}$ S5 | $F/F_{\text{MSY}}$ S2 | $F/F_{\text{MSY}}$ S3 | $F/F_{\text{MSY}}$ S5 |
|---|---|---|---|---|---|---|---|
| Anchovy | $f_{0.5}$ | -0.7 | -0.66 | -0.66 | 0.56 | 0.24 | 0.26 |
| Anchovy | $f^{\text{C}}_{0.15}$ | -0.59 | -0.54 | -0.5 | 0.86 | 0.73 | 0.6 |
| Anchovy | $\text{BT}_{0.50}$ | -0.68 | -0.51 | -0.33 | 0.84 | 0.2 | -0.07 |
| Anchovy | $\text{BT}_{50}f^{\text{C}}_{0.25}$ | -0.62 | -0.51 | -0.37 | 0.92 | 0.56 | 0.1 |
| Haddock | $f_{0.5}$ | -0.36 | -0.41 | -0.43 | 0.26 | 0.34 | 0.31 |
| Haddock | $f^{\text{C}}_{0.15}$ | -0.34 | -0.36 | -0.35 | 0.27 | 0.34 | 0.28 |
| Haddock | $\text{BT}_{0.50}$ | -0.36 | -0.39 | -0.37 | 0.28 | 0.3 | 0.27 |
| Haddock | $\text{BT}_{50}f^{\text{C}}_{0.25}$ | -0.35 | -0.37 | -0.35 | 0.3 | 0.33 | 0.33 |
| Halibut | $f_{0.5}$ | -0.11 | -0.15 | -0.19 | 0 | 0.07 | 0.09 |
| Halibut | $f^{\text{C}}_{0.15}$ | -0.07 | -0.07 | -0.13 | -0.01 | -0.02 | 0.02 |
| Halibut | $\text{BT}_{0.50}$ | -0.11 | -0.14 | -0.14 | 0.02 | 0.05 | 0.03 |
| Halibut | $\text{BT}_{50}f^{\text{C}}_{0.25}$ | -0.08 | -0.08 | -0.1 | -0.01 | 0 | 0.01 |

Table B11: Median absolute relative error (median(abs($(x_{\text{est}} - x_{\text{true}})/x_{\text{true}}$))) of estimated $B/B_{\text{MSY}}$ and $F/F_{\text{MSY}}$ for scenario 2 (observation SD=0.1), scenario 3 (baseline, observation SD=0.3), and scenario 5 (observation SD=0.7) for selected HCR and all species.

| Species | HCR | $B/B_{\text{MSY}}$ S2 | $B/B_{\text{MSY}}$ S3 | $B/B_{\text{MSY}}$ S5 | $F/F_{\text{MSY}}$ S2 | $F/F_{\text{MSY}}$ S3 | $F/F_{\text{MSY}}$ S5 |
|---|---|---|---|---|---|---|---|
| Anchovy | $f_{0.5}$ | 0.74 | 0.78 | 0.87 | 0.61 | 0.58 | 0.74 |
| Anchovy | $f^{\text{C}}_{0.15}$ | 0.61 | 0.58 | 0.61 | 0.86 | 0.74 | 0.77 |
| Anchovy | $\text{BT}_{0.50}$ | 0.7 | 0.69 | 0.8 | 0.85 | 0.76 | 0.79 |
| Anchovy | $\text{BT}_{50}f^{\text{C}}_{0.25}$ | 0.63 | 0.59 | 0.62 | 0.92 | 0.72 | 0.74 |
| Haddock | $f_{0.5}$ | 0.36 | 0.42 | 0.5 | 0.27 | 0.41 | 0.6 |
| Haddock | $f^{\text{C}}_{0.15}$ | 0.34 | 0.36 | 0.37 | 0.27 | 0.36 | 0.42 |
| Haddock | $\text{BT}_{0.50}$ | 0.36 | 0.4 | 0.47 | 0.29 | 0.39 | 0.6 |
| Haddock | $\text{BT}_{50}f^{\text{C}}_{0.25}$ | 0.35 | 0.37 | 0.38 | 0.3 | 0.37 | 0.48 |
| Halibut | $f_{0.5}$ | 0.17 | 0.2 | 0.32 | 0.15 | 0.23 | 0.46 |
| Halibut | $f^{\text{C}}_{0.15}$ | 0.16 | 0.2 | 0.26 | 0.15 | 0.23 | 0.34 |
| Halibut | $\text{BT}_{0.50}$ | 0.16 | 0.2 | 0.3 | 0.15 | 0.23 | 0.39 |
| Halibut | $\text{BT}_{50}f^{\text{C}}_{0.25}$ | 0.16 | 0.2 | 0.26 | 0.14 | 0.22 | 0.36 |

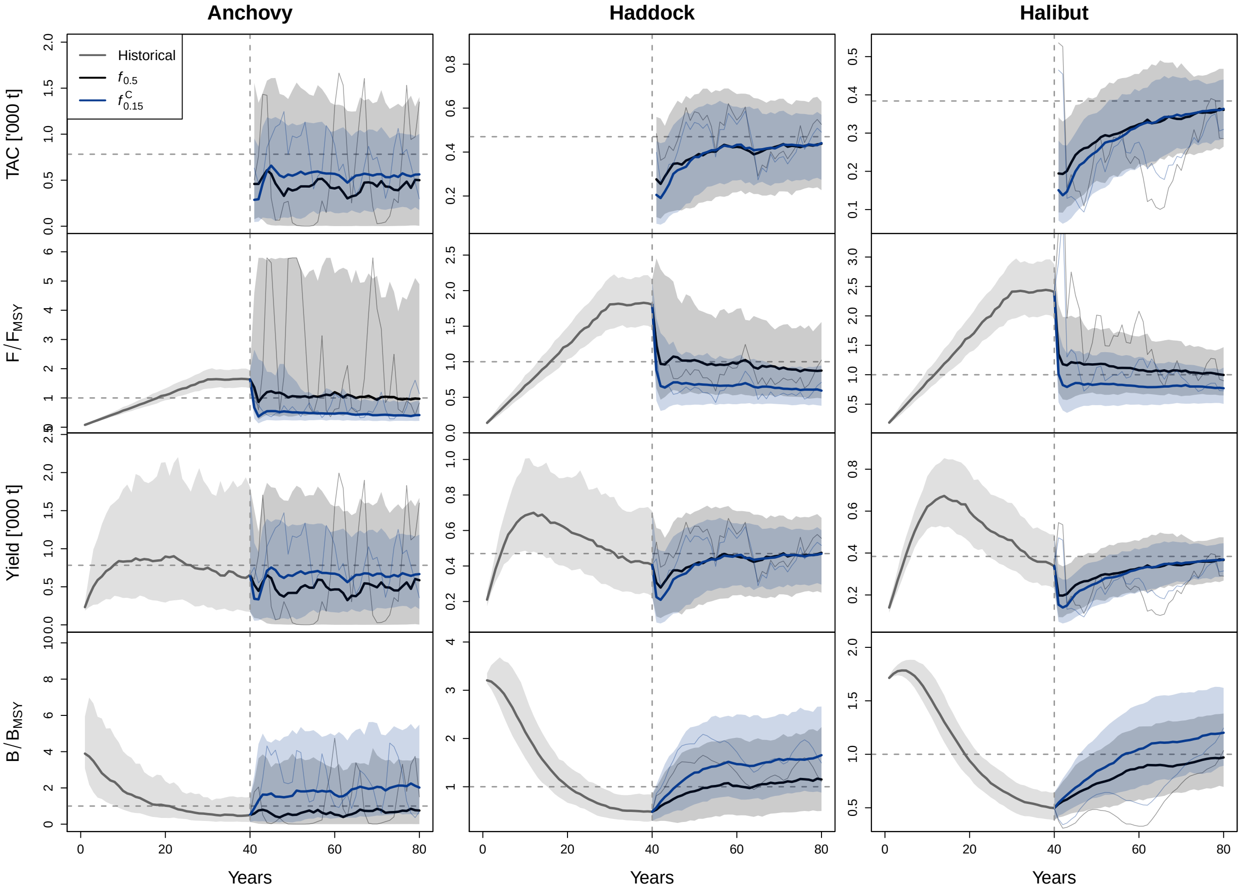


Figure B10: Trajectories of TAC, $F/F_{\mathrm{MSY}}$, yield, and $B/B_{\mathrm{MSY}}$ over historical and projection period. The thick coloured lines represent the median trajectories for the two HCRs, while the thin coloured lines represent the trajectory of a single replicate. The shaded area represents the 10th and 90th percentiles of the specific distributions. The vertical dashed line separates the historical (left of it) from the projection period (right to it). The horizontal dashed lines represent the reference points, i.e. 1 for the relative fishing mortality rate and biomass and MSY for TAC and yield.

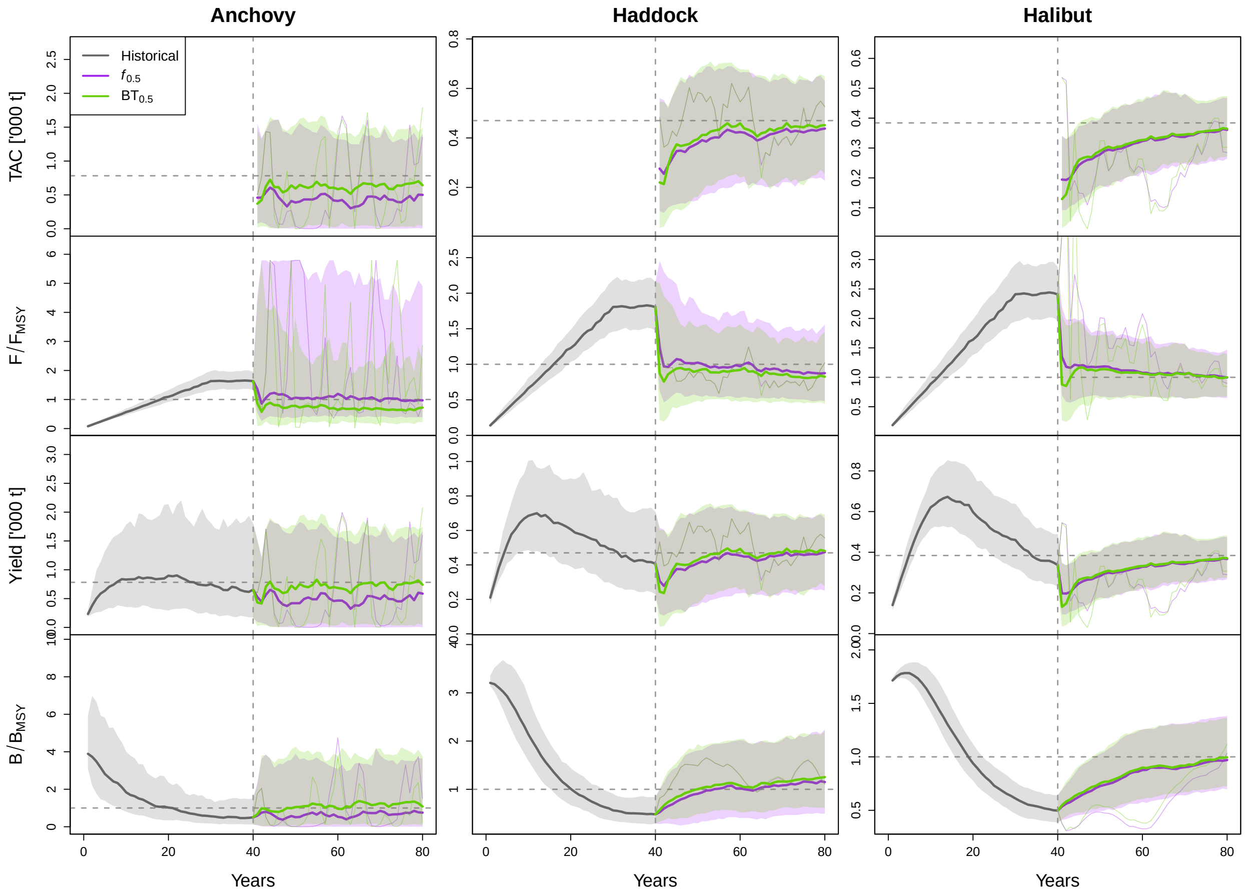


Figure B11: Trajectories of TAC, $F/F_{\mathrm{MSY}}$, yield, and $B/B_{\mathrm{MSY}}$ over historical and projection period. The thick coloured lines represent the median trajectories for the two HCRs, while the thin coloured lines represent the trajectory of a single replicate. The shaded area represents the 10th and 90th percentiles of the specific distributions. The vertical dashed line separates the historical (left of it) from the projection period (right to it). The horizontal dashed lines represent the reference points, i.e. 1 for the relative fishing mortality rate and biomass and MSY for TAC and yield.

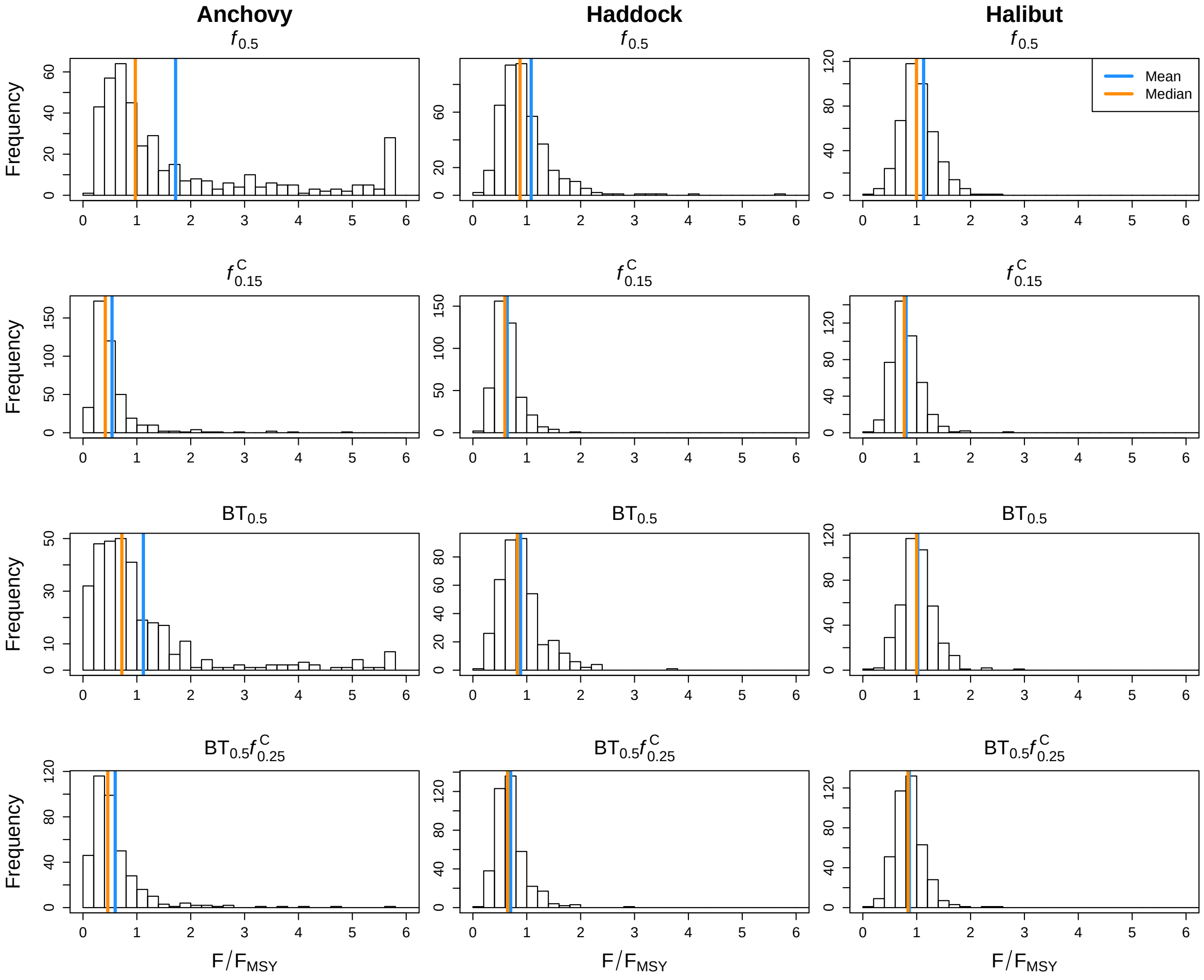


Figure B12: Distribution of $F/F_{\mathrm{MSY}}$ in the last year of the proejction period (year 80) for the selected HCRs: $f_{0.5}$, $f^{\mathrm{C}}_{0.15}$, $\mathrm{BT}_{0.5}$, $\mathrm{BT}_{0.5}f^{\mathrm{C}}_{0.25}$ (rows) for all species (columns). Vertical lines represent the mean (blue) and median (orange). In order to focus on the relevant range around the median and mean of the distribution, replicates above 6 were removed. Thus, up to 3 and 1 0 replicates were removed for anchovy and haddock, respectively.

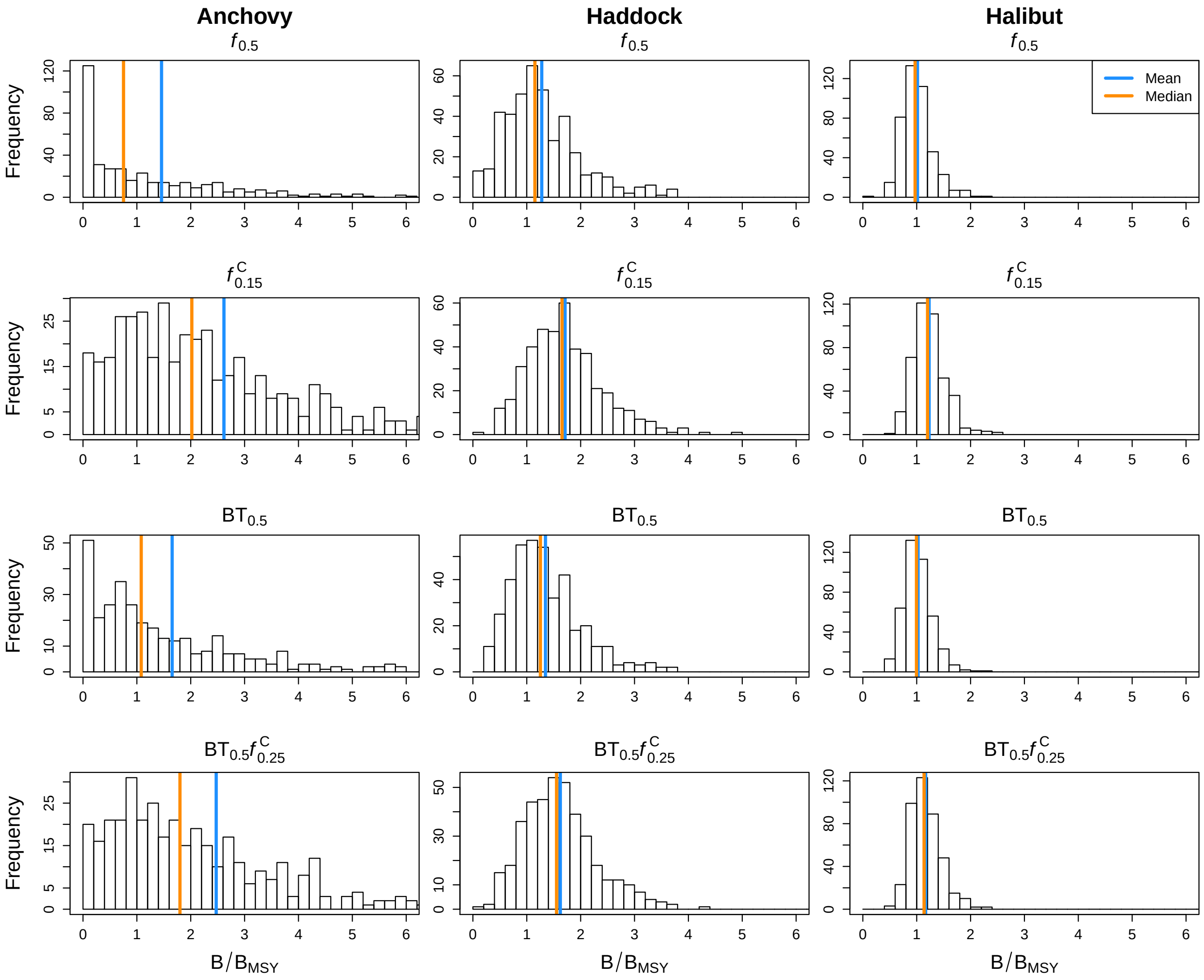


Figure B13: Distribution of $B/B_{\mathrm{MSY}}$ in the last year of the proejction period (year 80) for the selected HCRs: $f_{0.5}$, $f^{\mathrm{C}}_{0.15}$, $\mathrm{BT}_{0.5}$, $\mathrm{BT}_{0.5}f^{\mathrm{C}}_{0.25}$ (rows) for all species (columns). Vertical lines represent the mean (blue) and median (orange). In order to focus on the relevant range around the median and mean of the distribution, replicates above 6 were removed. Thus, up to 35 replicates were removed for anchovy.

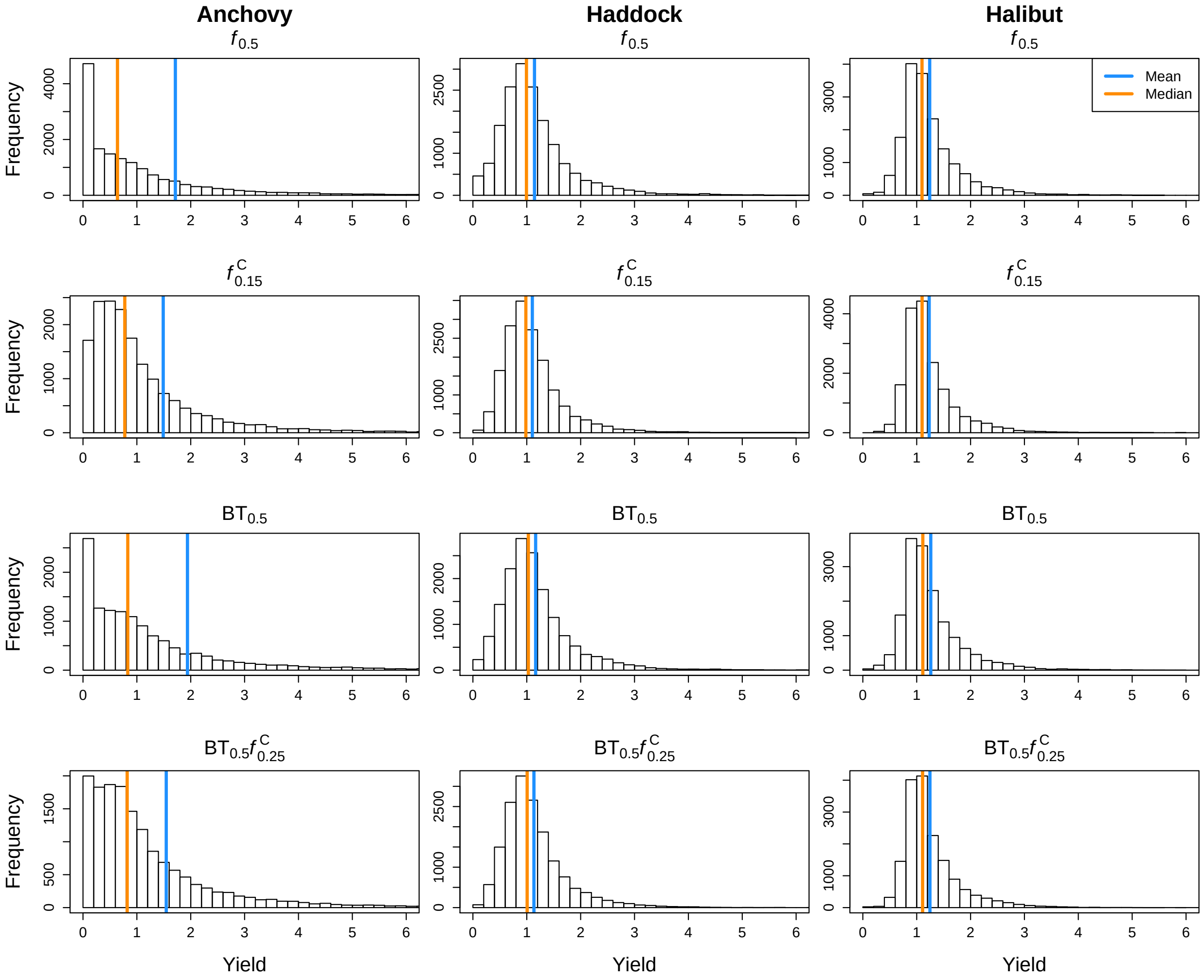


Figure B14: Distribution of the relative yield in the last year of the proejction period (year 80) for the selected HCRs: $f_{0.5}$, $f^{C}_{0.15}$, $\mathrm{BT}_{0.5}$, $\mathrm{BT}_{0.5}f^{C}_{0.25}$ (rows) for all species (columns). Vertical lines represent the mean (blue) and median (orange). In order to focus on the relevant range around the median and mean of the distribution, replicates above 6 were removed. Thus, up to 453 and 26 replicates were removed for anchovy and haddock, respectively.

89 # Supplementary Section C: Sensitivity analysis

Table C1: Proportion of converged replicates in percent for the baseline scenario (S1) and the sensitivity scenarios (S6-S13).

| Scenario | HCR | Anchovy | Haddock | Halibut |
|---|---|---|---|---|
| S1 | $f_{0.5}$ | 81 | 85 | 85 |
| S1 | $f^{C}_{0.15}$ | 86 | 83 | 86 |
| S1 | $BT_{0.50}$ | 66 | 79 | 82 |
| S1 | $BT_{50}f^{C}_{0.25}$ | 77 | 81 | 83 |
| S6 | $f_{0.5}$ | 74 | 59 | 36 |
| S6 | $f^{C}_{0.15}$ | 75 | 57 | 34 |
| S6 | $BT_{0.50}$ | 59 | 52 | 35 |
| S6 | $BT_{50}f^{C}_{0.25}$ | 68 | 54 | 31 |
| S7 | $f_{0.5}$ | 67 | 27 | 4 |
| S7 | $f^{C}_{0.15}$ | 71 | 27 | 4 |
| S7 | $BT_{0.50}$ | 37 | 21 | 3 |
| S7 | $BT_{50}f^{C}_{0.25}$ | 53 | 24 | 3 |
| S8 | $f_{0.5}$ | 78 | 74 | 82 |
| S8 | $f^{C}_{0.15}$ | 81 | 71 | 81 |
| S8 | $BT_{0.50}$ | 65 | 70 | 78 |
| S8 | $BT_{50}f^{C}_{0.25}$ | 73 | 69 | 78 |
| S9 | $f_{0.5}$ | 72 | 84 | 86 |
| S9 | $f^{C}_{0.15}$ | 81 | 82 | 85 |
| S9 | $BT_{0.50}$ | 52 | 78 | 83 |
| S9 | $BT_{50}f^{C}_{0.25}$ | 73 | 80 | 83 |
| S10 | $f_{0.5}$ | 41 | 66 | 78 |
| S10 | $f^{C}_{0.15}$ | 66 | 64 | 73 |
| S10 | $BT_{0.50}$ | 29 | 52 | 63 |
| S10 | $BT_{50}f^{C}_{0.25}$ | 53 | 55 | 62 |
| S11 | $f_{0.5}$ | 11 | 33 | 30 |
| S11 | $f^{C}_{0.15}$ | 14 | 28 | 35 |
| S11 | $BT_{0.50}$ | 2 | 19 | 26 |
| S11 | $BT_{50}f^{C}_{0.25}$ | 8 | 23 | 30 |
| S12 | $f_{0.5}$ | 65 | 90 | 98 |
| S12 | $f^{C}_{0.15}$ | 60 | 92 | 96 |
| S12 | $BT_{0.50}$ | 63 | 90 | 97 |
| S12 | $BT_{50}f^{C}_{0.25}$ | 62 | 91 | 96 |
| S13 | $f_{0.5}$ | 82 | 82 | 90 |
| S13 | $f^{C}_{0.15}$ | 85 | 84 | 89 |
| S13 | $BT_{0.50}$ | 63 | 81 | 89 |
| S13 | $BT_{50}f^{C}_{0.25}$ | 76 | 81 | 87 |

Table C2: Proportion of converged assessments in percent for baseline scenario, the scenario without priors (scenario 7), and the scenario with a shorter time series (scenario 11). In total, 20000 asssessments were conducted for each HCR, scenario and species (projection period of 40 years and 500 replicates).

| Scenario | HCR | Basline | Scenario 7 | Scenario 11 |
|---|---|---|---|---|
| Anchovy | $f_{0.5}$ | 98 | 96 | 85 |
| Anchovy | $f^{C}_{0.15}$ | 98 | 96 | 87 |
| Anchovy | $\mathrm{BT}_{0.50}$ | 95 | 84 | 70 |
| Anchovy | $\mathrm{BT}_{50}f^{C}_{0.25}$ | 98 | 90 | 80 |
| Haddock | $f_{0.5}$ | 98 | 78 | 88 |
| Haddock | $f^{C}_{0.15}$ | 98 | 78 | 86 |
| Haddock | $\mathrm{BT}_{0.50}$ | 97 | 71 | 76 |
| Haddock | $\mathrm{BT}_{50}f^{C}_{0.25}$ | 98 | 70 | 81 |
| Halibut | $f_{0.5}$ | 98 | 46 | 86 |
| Halibut | $f^{C}_{0.15}$ | 98 | 47 | 87 |
| Halibut | $\mathrm{BT}_{0.50}$ | 97 | 46 | 82 |
| Halibut | $\mathrm{BT}_{50}f^{C}_{0.25}$ | 97 | 46 | 84 |

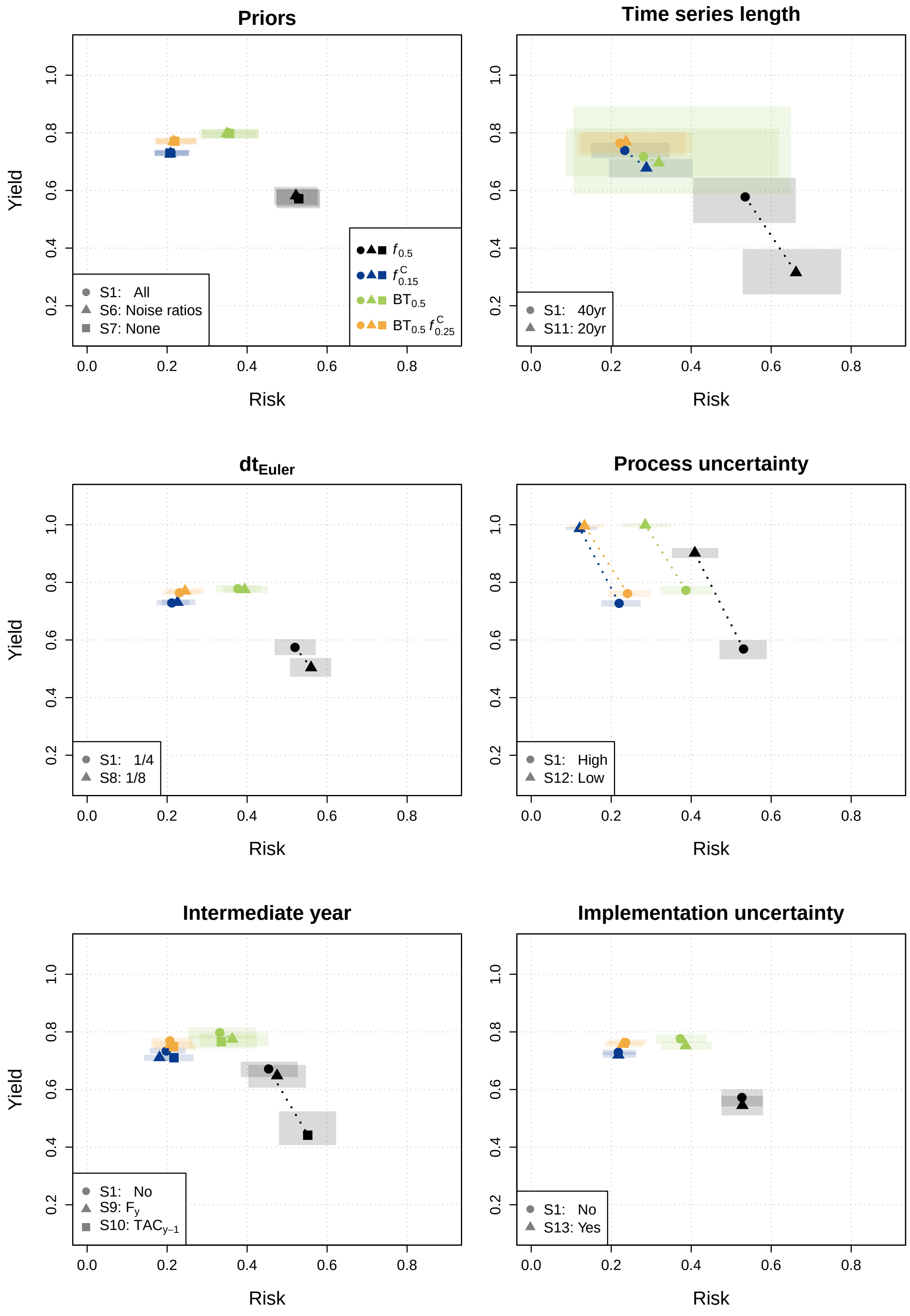


Figure C1: Risk-return trade-off for eight sensitivity scenarios (S6-S13) in comparison to baseline scenario (S1) for **dab**. The colours represent the four selected HCRs: $f_{0.5}$ (black), $f^{C}_{0.15}$ (blue), $BT_{0.5}$ (green), $BT_{0.5}f^{C}_{0.25}$ (yellow) used in all plotting panels. The symbols represents the various scenarios, where the circle represents the baseline scenario and the other symbols are re-defined in each plotting panel. The dotted lines connect the baseline scenario for each HCR with the specific sensitivity scenario.

Table C3: Annual absolute variability in yield (AAV) for sensitivity scenarios and species Anchovy. Scenarios code: S1 = Baseline, S6 = No logn prior, S17 = No priors, S8 = 8 time steps, S9 = Intermediate year with continuing F, S10 = Intermediate year with last year's TAC, S11 = Time series of 20yr, S12 = lower process uncertainty, S13 = Implementation error. For more information, please refer to Table A5.

| HCR | S1 | S6 | S7 | S8 | S9 | S10 | S11 | S12 | S13 |
|---|---|---|---|---|---|---|---|---|---|
| $f_{0.5}$ | 0.459 | 0.457 | 0.458 | 0.468 | 0.438 | 0.446 | 0.56 | 0.258 | 0.492 |
| $f^{\mathrm{C}}_{0.15}$ | 0.312 | 0.311 | 0.312 | 0.322 | 0.249 | 0.283 | 0.354 | 0.141 | 0.358 |
| $\mathrm{BT}_{0.50}$ | 0.499 | 0.497 | 0.498 | 0.516 | 0.465 | 0.481 | 0.471 | 0.316 | 0.55 |
| $\mathrm{BT}_{50}f^{\mathrm{C}}_{0.25}$ | 0.366 | 0.365 | 0.366 | 0.381 | 0.31 | 0.341 | 0.385 | 0.17 | 0.411 |

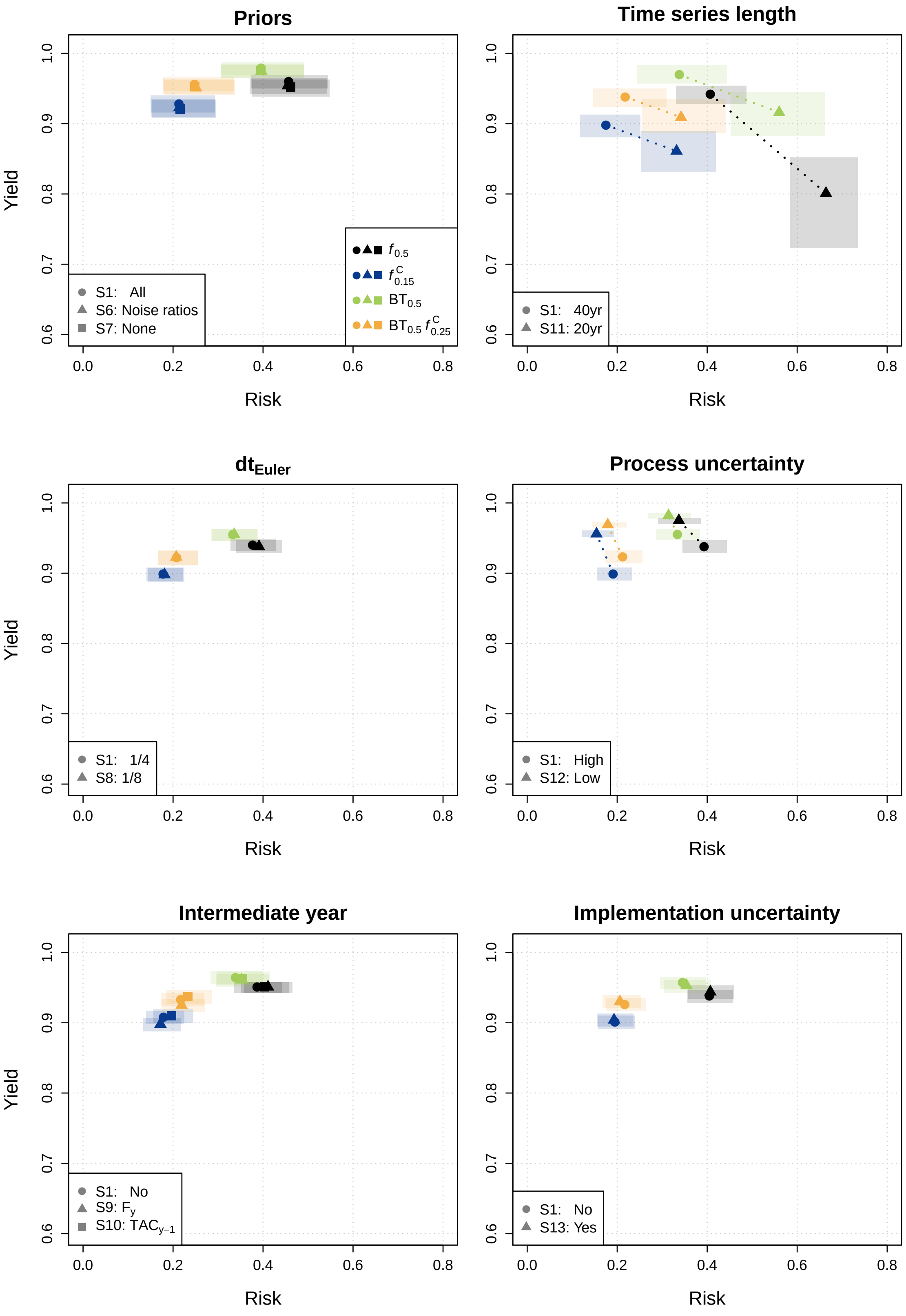


Figure C2: Risk-return trade-off for eight sensitivity scenarios (S6-S13) in comparison to baseline scenario (S1) for **haddock**. The colours represent the four selected HCRs: $f_{0.5}$ (black), $f^{C}_{0.15}$ (blue), $\mathrm{BT}_{0.5}$ (green), $\mathrm{BT}_{0.5} f^{C}_{0.25}$ (yellow) used in all plotting panels. The symbols represents the various scenarios, where the circle represents the baseline scenario and the other symbols are re-defined in each plotting panel. The dotted lines connect the baseline scenario for each HCR with the specific sensitivity scenario.

Table C4: Annual absolute variability in yield (AAV) for sensitivity scenarios and species Haddock. Scenarios code: S1 = Baseline, S6 = No logn prior, S17 = No priors, S8 = 8 time steps, S9 = Intermediate year with continuing F, S10 = Intermediate year with last year's TAC, S11 = Time series of 20yr, S12 = lower process uncertainty, S13 = Implementation error. For more information, please refer to Table A5.

| HCR | S1 | S6 | S7 | S8 | S9 | S10 | S11 | S12 | S13 |
|---|---|---|---|---|---|---|---|---|---|
| $f_{0.5}$ | 0.172 | 0.175 | 0.176 | 0.159 | 0.179 | 0.181 | 0.259 | 0.092 | 0.239 |
| $f_{0.15}^{\mathrm{C}}$ | 0.13 | 0.135 | 0.134 | 0.122 | 0.116 | 0.127 | 0.168 | 0.067 | 0.214 |
| $\mathrm{BT}_{0.50}$ | 0.197 | 0.2 | 0.2 | 0.189 | 0.209 | 0.21 | 0.28 | 0.095 | 0.265 |
| $\mathrm{BT}_{50}f_{0.25}^{\mathrm{C}}$ | 0.149 | 0.152 | 0.151 | 0.143 | 0.14 | 0.15 | 0.181 | 0.075 | 0.226 |

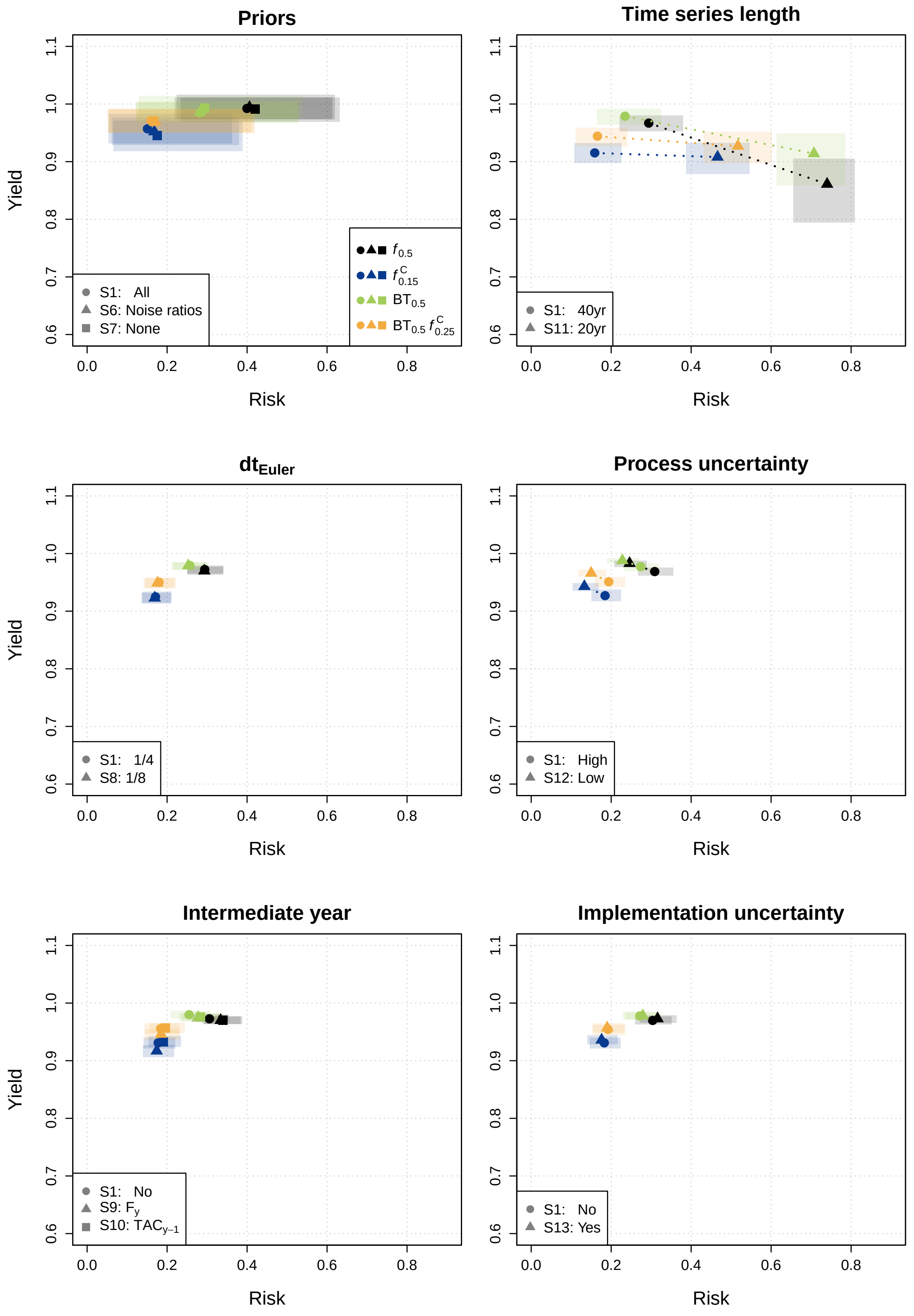


Figure C3: Risk-return trade-off for eight sensitivity scenarios (S6-S13) in comparison to baseline scenario (S1) for **Halibut**. The colours represent the four selected HCRs: $f_{0.5}$ (black), $f^{C}_{0.15}$ (blue), $BT_{0.5}$ (green), $BT_{0.5}f^{C}_{0.25}$ (yellow) used in all plotting panels. The symbols represents the various scenarios, where the circle represents the baseline scenario and the other symbols are re-defined in each plotting panel. The dotted lines connect the baseline scenario for each HCR with the specific sensitivity scenario.

Table C5: Annual absolute variability in yield (AAV) for sensitivity scenarios and species Halibut. Scenarios code: S1 = Baseline, S6 = No logn prior, S17 = No priors, S8 = 8 time steps, S9 = Intermediate year with continuing F, S10 = Intermediate year with last year's TAC, S11 = Time series of 20yr, S12 = lower process uncertainty, S13 = Implementation error. For more information, please refer to Table A5.

| HCR | S1 | S6 | S7 | S8 | S9 | S10 | S11 | S12 | S13 |
|---|---|---|---|---|---|---|---|---|---|
| $f_{0.5}$ | 0.132 | 0.144 | 0.146 | 0.106 | 0.123 | 0.123 | 0.216 | 0.064 | 0.207 |
| $f^{C}_{0.15}$ | 0.105 | 0.108 | 0.108 | 0.082 | 0.084 | 0.089 | 0.15 | 0.05 | 0.193 |
| $BT_{0.50}$ | 0.155 | 0.153 | 0.158 | 0.119 | 0.134 | 0.133 | 0.25 | 0.069 | 0.217 |
| $BT_{50}f^{C}_{0.25}$ | 0.127 | 0.129 | 0.124 | 0.093 | 0.097 | 0.101 | 0.161 | 0.056 | 0.2 |

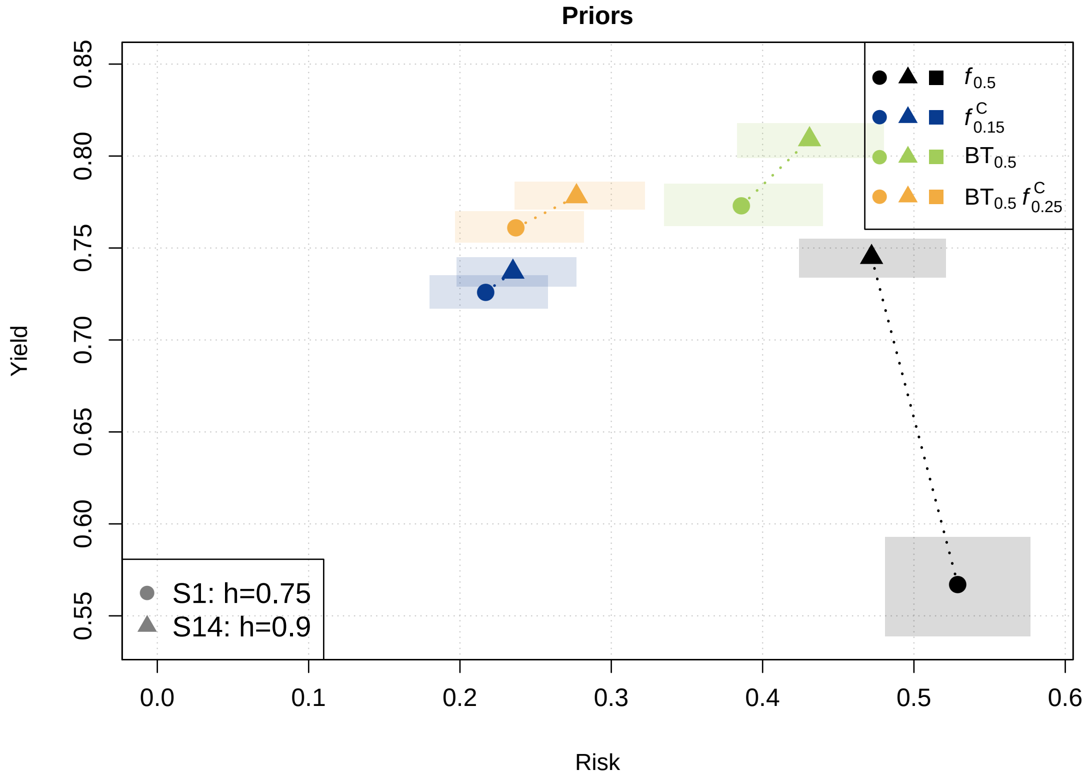


Figure C4: Risk-return trade-off for anchovy when assuming a steepness of the stock-recruitment relationship of 0.9 (Scenario S14) instead of 0.75 (Baseline scenario S1). The colours represent the four selected HCRs: $f_{0.5}$ (black), $f^{C}_{0.15}$ (blue), $BT_{0.5}$ (green), $BT_{0.5}f^{C}_{0.25}$ (yellow) used in all plotting panels. The symbols represents the various scenarios, where the circle represents the baseline scenario (h=0.75) and the triangle represents the higher steepness parameter (h=0.9). The dotted lines connect the baseline scenario for each HCR with the specific sensitivity scenario.

Table C6: Annual absolute variability in yield (AAV) for anchovy and the scenario with different steepness. Scenarios code: S3 = Baseline (steepness parameter (h) of 0.75) and S14 = steepness parameter (h) of 0.9. For more information, please refer to Table A5.

| HCR | S1 | S14 |
|---|---|---|
| $f_{0.5}$ | 0.465 | 0.455 |
| $f^{C}_{0.15}$ | 0.314 | 0.313 |
| $BT_{0.50}$ | 0.519 | 0.531 |
| $BT_{50}f^{C}_{0.25}$ | 0.375 | 0.383 |
